# Transient Non-local Interactions Dominate the Dynamics of Measles Virus N_TAIL_

**DOI:** 10.1101/2024.07.22.604679

**Authors:** Lillian Otteson, Gabor Nagy, John Kunkel, Gerdenis Kodis, Lars V. Bock, Christophe Bignon, Sonia Longhi, Wenwei Zheng, Helmut Grubmüller, Andrea C. Vaiana, Sara M. Vaiana

## Abstract

The RNA genome of measles virus is encapsidated by the nucleoprotein within a helical nucleocapsid that serves as template for both transcription and replication. The intrinsically disordered domain of the nucleoprotein (N_TAIL_), partly protruding outward from the nucleocapsid, is essential for binding the polymerase complex responsible for viral transcription and replication. As for many IDPs, binding of N_TAIL_ occurs through a short molecular recognition element (MoRE) that folds upon binding, with the majority of N_TAIL_ remaining disordered. Though N_TAIL_ regions far from the MoRE influence the binding affinity, interactions between them and the MoRE have not been investigated in depth. Using an integrated approach, relying on photo-induced electron transfer (PET) experiments between tryptophan and cysteine pairs placed at different positions in the protein under varying salt and pH conditions, combined with simulations and analytical models, we identified transient interactions between two disordered regions distant in sequence, which dominate N_TAIL_ dynamics, and regulate the conformational preferences of both the MoRE and the entire N_TAIL_ domain. Co-evolutionary analysis corroborates our findings and suggests an important functional role for the same intramolecular interactions. We propose mechanisms by which these non-local interactions may regulate binding to the measles phosphoprotein, polymerase recruitment, and ultimately viral transcription and replication. Our findings may be extended to other IDPs, where non-local intra-protein interactions affect the conformational preferences of intermolecular binding sites.

## Introduction

The measles virus (MeV) is a non-segmented, single-stranded, negative sense RNA virus, in which the viral genome is wrapped within a helical nucleocapsid, made of thousands of repeats of the nucleoprotein (N) ^1^. This structure is shared among all Paramyxo- and Pneumoviruses, which include common human viruses (e.g., MeV, mumps, respiratory syncytial virus, metapneumovirus, and other para-influenza viruses) and animal viruses (e.g., Sendai virus, Newcastle disease virus and rinderpest virus), as well as zoonotic biosafety level 4 agents of great concern to public health (e.g., Hendra and Nipah viruses, respectively HeV and NiV)^2^. The N protein not only provides the interaction with the RNA and holds the nucleocapsid in its helical structure, but it is the substrate used for viral transcription and replication. Viral transcription and replication in Paramyxoviruses are ensured by a complex formed by the large-polymerase (L) and the phosphoprotein (P), where the latter enables tethering L to the nucleoprotein-RNA (N-RNA) template for transcription and replication^2,3^.

In MeV, N consists of a folded 400 amino-acid (a.a.) domain (N_CORE_) which binds the RNA and forms the structured helical nucleocapsid, and of an intrinsically disordered 125 aa. C-terminal domain (N_TAIL_), which partly protrudes radially outward from the helical nucleocapsid structure^4–7^. The intrinsically disordered N_TAIL_ domain binds to the folded X domain (P_XD_) of the phosphoprotein (P) and is essential for MeV transcription and replication. P in turn, binds as a tetramer to both L and N_CORE_ ^6,8–11^. A similar structure and binding of N_TAIL_ to P_XD_ are found in NiV and HeV^3,12^. Binding of N_TAIL_ to P_XD_ is thought to assist in positioning the L appropriately on the nucleocapsid to read the RNA, while allowing efficient transcription re-initiation at intergenic regions^13^ and hence progression from one reading frame to the next. It may also assist in maintaining contact between L and the nucleocapsid template, preventing it from leaving before transcribing all the viral genome^11,14^. In addition, the presence of N_TAIL_ loosens the nucleocapsid helical structure, possibly facilitating transcription and replication by L^11^.

Because of its crucial role in transcription and replication, binding of MeV N_TAIL_ to P_XD_ has been extensively studied^15–21^. It occurs through coupled folding and binding of a short molecular recognition region (MoRE) of 18 a.a. (grey in Fig. 1)^21^. Upon binding, the MoRE folds into an α-helix, forming a four-helix bundle with three helices from P_XD_ (PDB: 1t6o^8,9^). However, the majority of N_TAIL_ (i.e., the remaining 109 a.a. on the two sides of the MoRE) remains disordered in the bound complex^21^. The N_TAIL_-P_XD_ complex is structurally similar in MeV, NiV and HeV^3,12^. While the regions outside the MoRE (from here on referred to as “flanking regions”) do not fold, or engage in stable binding interactions^16,17^, they modulate binding affinities, as indicated by the binding affinity increase in truncated variants of N_TAIL_ containing the intact MoRE^22,23^. This increase might be due to a pure excluded volume (entropic) repulsion of the flanking regions, as well as to potential interactions involving the flanking regions of N_TAIL_ (either at the intra- or inter-protein level)^22,24,25^.

**Figure 1.**
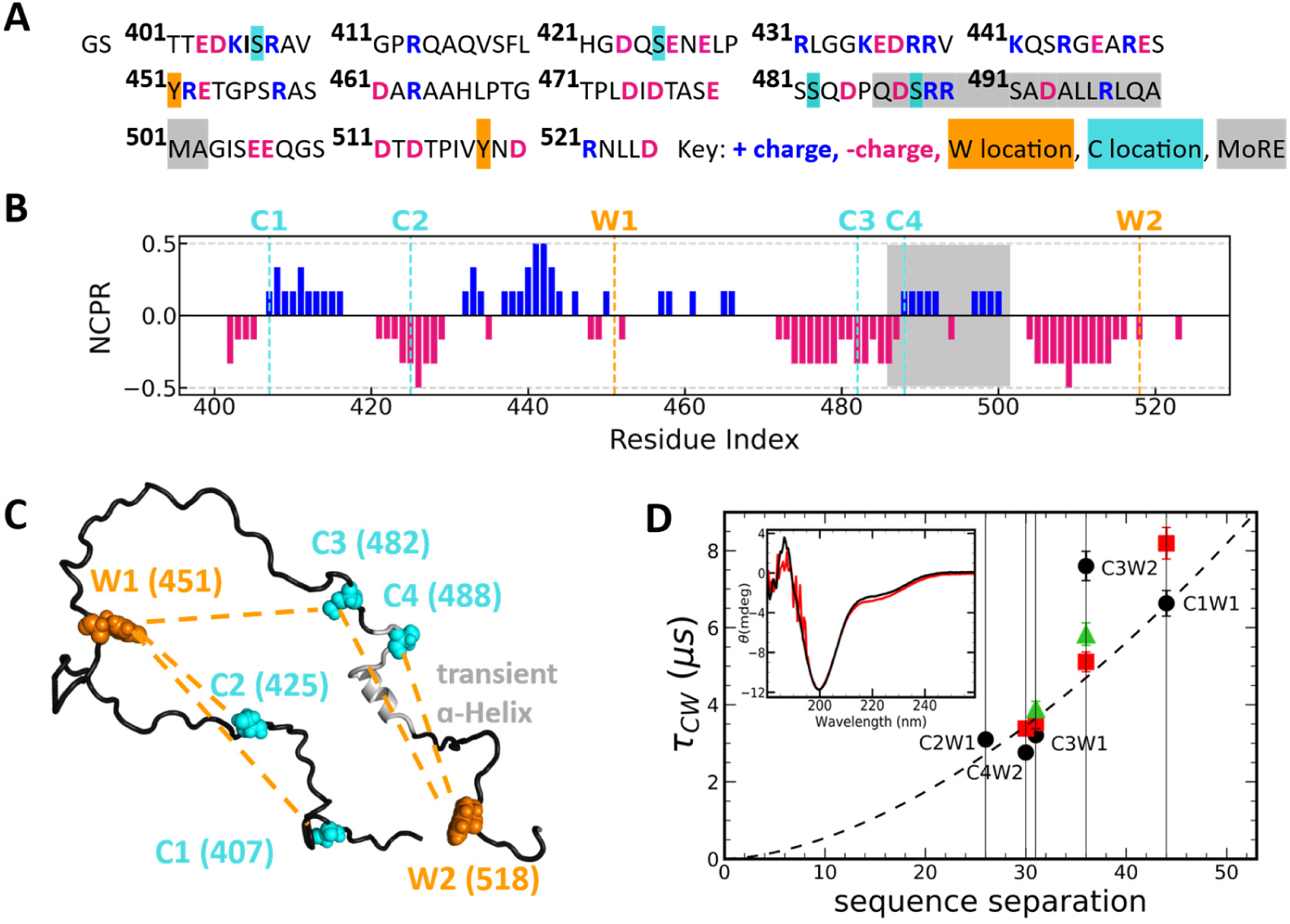
Results of photo-induced electron transfer (PET) experiments at T=20 °C, on NTAIL variants containing a unique C-W pair. A) NTAIL sequence, showing charged residues at pH 7.6, location of Y to W, and S to C substitutions, and location of MoRE, which folds into a helix upon binding to PXD. B) Net charge per residue calculated using a window size of six amino acids. C) W and C positions. Each variant contains a unique C-W pair. The C-W distance in each variant is indicated by orange dashed lines. C3W2 and C4W2 variants encompass the MoRE transient helix. D) C-W relaxation time r_CW,_ due to PET as a function of C-W sequence separation. Experimental conditions: pH 7.6 and 150 mM NaCl (black circles); pH 4.0 and 150 mM NaCl (red squares); pH 7.6 and 500mM NaCl (green triangles). Dashed line: expected scaling of r_CW,_ from a SAW-ν homopolymer model at pH 7.6 (black). Inset: Comparison of NTAIL circular dichroism spectra at pH 7.6 (black) and pH 4 (red).

For N_TAIL_ free in solution, the secondary structure propensity of the MoRE region has been characterized by NMR on full length N ^7^, and by all-atom simulation studies of the MoRE fragment^19,26^. In contrast, we know very little about potential interactions between residues that are distant in sequence (non-local interactions), despite their particular importance for N_TAIL_ binding to P_XD_^24,27^ and, more generally, for IDP function. Large-scale reconfiguration dynamics can regulate IDP binding mechanisms, rates, and affinities; in fact, examples of this are found even in simple models of proteins undergoing two-state conformational transitions^28^. For N_TAIL_, we expect interactions involving the regions flanking the MoRE on either side to confer binding specificity to the IDP by dictating large-scale reconfiguration dynamics. In addition, non-local interactions potentially act as allosteric regulators in IDPs by stabilizing certain conformational states. The resulting shift in the population of states within the ensemble can lead to allosteric regulation, as described in the ensemble allosteric model for IDPs^29,30^.

Probing non-local intra-chain interactions is therefore essential for a molecular understanding of IDP dynamics, binding, and function in general^22,31–33^. Direct experimental probing of such interactions is particularly difficult, due to their transient nature, and the fact that IDPs undergo large conformational changes on sub-microsecond time scales. Atomistic simulations of IDPs, on the other hand, are equally challenging due to the wide conformational space sampled by IDPs. To overcome these experimental and theoretical challenges, we leveraged the high sensitivity of IDP dynamics to non-local interactions. We experimentally quantified the dynamics of full-length N_TAIL_ in solution, by probing intra-molecular contact formation times, using photo-induced electron transfer (PET) between a tryptophan (W) and a cysteine (C) placed in the sequence^34–37^. Using this technique, we measured contact formation rates along the well-defined C-W distance reaction coordinate, probing large-scale backbone motions on the hundreds of nanoseconds to microsecond internal reorganization time scale^36^, which is most relevant to IDP function^38^. With minimal perturbation to the sequence, compared to introducing large prosthetic dyes, we achieved a high sensitivity at short spatial distances. To obtain a structural interpretation and a causal description, we compared the measurements to corresponding observables derived from both simulations and analytical models. Our integrated approach allowed us to identify key intra-protein interactions that extend beyond the MoRE region, and play an important role in N_TAIL_ dynamics and likely also in function.

## Results

### Photo-induced electron transfer reveals the heteropolymeric nature of N_TAIL_ dynamics

To characterize the dynamics of N_TAIL_ in solution, we probed intra-molecular contact formation, using photo-induced electron transfer (PET) between tryptophan (W) and cysteine (C) residues at different positions in the sequence.^35–37^ After exciting W to the triplet state using a nanosecond UV laser pulse, we monitored the triplet state population in time, via transient absorption measurements. As N_TAIL_ undergoes spontaneous conformational changes, W can come into contact with C and PET can occur. During PET, the excited-state (triplet) electron is transferred from W to C, forming a W radical cation which eventually relaxes to the ground state. The relaxation time due to PET, r_CW_, therefore provides information on how often and quickly C and W come into contact after W excitation. If the C-W contact formation rate is of the same order of magnitude, or faster than the natural decay rate of W, r_CW_ can be obtained from the short time behavior of the triplet state relaxation curves (Supporting Fig. S1).

The wild-type N_TAIL_ sequence (Fig. 1A) does not contain native W or C residues. To probe the N_TAIL_ conformational dynamics, we generated five N_TAIL_ variants (PET variants), each bearing a unique C-W pair, to probe the five distances indicated in Fig. 1C. To minimize perturbation of the chemical properties, we substituted native tyrosine (Y) residues with W, and native serines (S) with C residues. To determine the natural decay rate of excited W at the two positions, we also generated two variants with the respective W substitution, but lacking any C substitution. The Supporting Information contains the sequences of all seven N_TAIL_ variants (Table S1) and details about their production and purification (Supporting Methods S1).

Relaxation times r_CW_ were estimated from multiple, independent sample preparations for each N_TAIL_ variant, and a series of repeated measurements to account for W and C photodamage during the measurements (Supporting Methods S2). All measured relaxation curves and fitted functions to obtain r_CW_ are shown in Fig S3-S16, with the fit results summarized in Table S2.

We first performed measurements with the five PET variants of N_TAIL_ under near physiological conditions (i.e. 15mM Tris, 150mM NaCl, 1mM TCEP, pH 7.6, in the following referred to as “reference conditions”). Figure 1D shows the r_CW_ (black circles) estimated from these measurements as a function of the W-C sequence separation (|i-j|).

To test whether the estimated r_CW_ can be explained based on C-W sequence separation alone, we compared them to relaxation times expected from a simple homopolymer model (Fig. 1D, dashed line), where the only non-local interactions between the residues are excluded volume interactions. In this model, both the physical distance in space between C and W at positions *i* and *j* as well as the corresponding relaxation time r_ij_ scale with the sequence separation |i-j|.^39^ The scaling relation for relaxation times r_ij_ = 1_f_ |i − j|^ς1^ depends on a prefactor 1_f_ and an exponent ς_1_. The exponent ς_1_.= 1.69 (corresponding to a Flory scaling exponent v of 0.52) was estimated from small-angle X-ray scattering (SAXS) measurements of N_TAIL_^6^ based on a homopolymer model (SAW-v model^40^, Supporting Methods S4). The prefactor 1_f_ = 0.011 μs was obtained by fitting to the estimated r_CW_ (Fig. 1D) from PET experiments.

The general trend of the estimated r_CW_ at physiological pH (Fig. 1D, black circles) follows the simple homopolymer scaling (dashed curve), however some contacts form faster and others slower than expected. This dynamic heterogeneity (deviation from homopolymer model) observed along the sequence is most striking when comparing the C3W2 (|i-j| = 36) and C4W2 (|i-j| = 30) variants for which the C-W pairs encompass the MoRE region. These variants share the W position and differ by just six residues in sequence separation between W and C (Figs. 1A-1C). According to the homopolymer model, such a difference in sequence separation would result in a r_CW_ increase by a factor of 1.3. In contrast, the r_CW_ estimated from relaxation curves increases by a factor of 2.8. This considerable deviation from the homopolymer behavior is primarily due to the measured 7.6 µs relaxation time for C3W2 that is substantially longer than the 4.7 µs homopolymer estimate. The fact that this anomalously slow relaxation time is not reproduced by the homopolymer model indicates that the N_TAIL_ dynamics near the MoRE region is considerably affected by more complex heteropolymeric interactions, likely secondary structure formation, electrostatic interactions between charged residues, or solvent-induced interactions.

### Charged amino acids partially account for N_TAIL_ dynamic heterogeneity

To test whether interactions between charged residues or secondary structure formation causes the dynamic heterogeneity, we performed additional experiments at low pH and increased salt concentrations, i.e. under conditions that should modulate electrostatic interactions.

The N_TAIL_ sequence contains 40 charged residues (13 Asp, 10 Glu, 14 Arg and 3 Lys), often clustered together (Fig. 1B). If electrostatic interactions between charged residues were markedly contributing to the observed dynamic heterogeneity, we would expect that screening the interactions by increasing the salt concentration would decrease or eliminate the heterogeneity. Indeed, increasing the salt concentration from 150mM (Fig. 1D, black circles) to 500mM (green triangles) results in a r_CW_ decrease from 7.6 μs to 5.8 μs for C3W2, and an increase from 3.2 μs to 3.9 μs for C3W1, bringing both data points close to the homopolymer curve. This suggests that electrostatic interactions are a major contributor to the observed dynamic heterogeneity of N_TAIL_.

Next, we investigated why electrostatic interactions accelerate relaxation for the C3W1 variant, but slow down relaxation for the C3W2 variant. We hypothesized that changes in contact dynamics are caused by the non-uniform distribution of charged amino acids (charge patterning^41,42^) along the N_TAIL_ sequence. However, the increased electrostatic screening at higher salt concentrations could also affect the stability of the transient helix in the MoRE region, and thereby modulate the C3W2 relaxation time. To address these two possibilities, we performed coarse-grained molecular dynamics (CG) simulations that allowed us to explicitly include both charge patterning and the secondary structure contributions.

For these simulations we used a modified version of the recently developed residue-based HPS model^43^ for IDPs. The CG model was adjusted to reproduce the experimental radius of gyration of N_TAIL_ obtained from SAXS^6^ and the helical propensity of the MoRE region from NMR^7^. The effects of varying salt concentrations in the CG model were described by Debye-Hückel electrostatic screening^44^. Additional details on the CG simulations are provided in Supporting Methods S5.

To determine the effect of increased salt concentration (150 mM to 500 mM), we computed the mean distance for each pair of amino acids *i* and *j* from the CG simulations at both salt concentrations. Fig. 2A shows the difference of these means, highlighting the compaction/expansion of each N_TAIL_ segment. In the simulations, the N-terminal part of N_TAIL_ up to C3 expands at higher salt concentrations (red areas), whereas the C-terminal part becomes more compact (blue areas).

**Figure 2.**
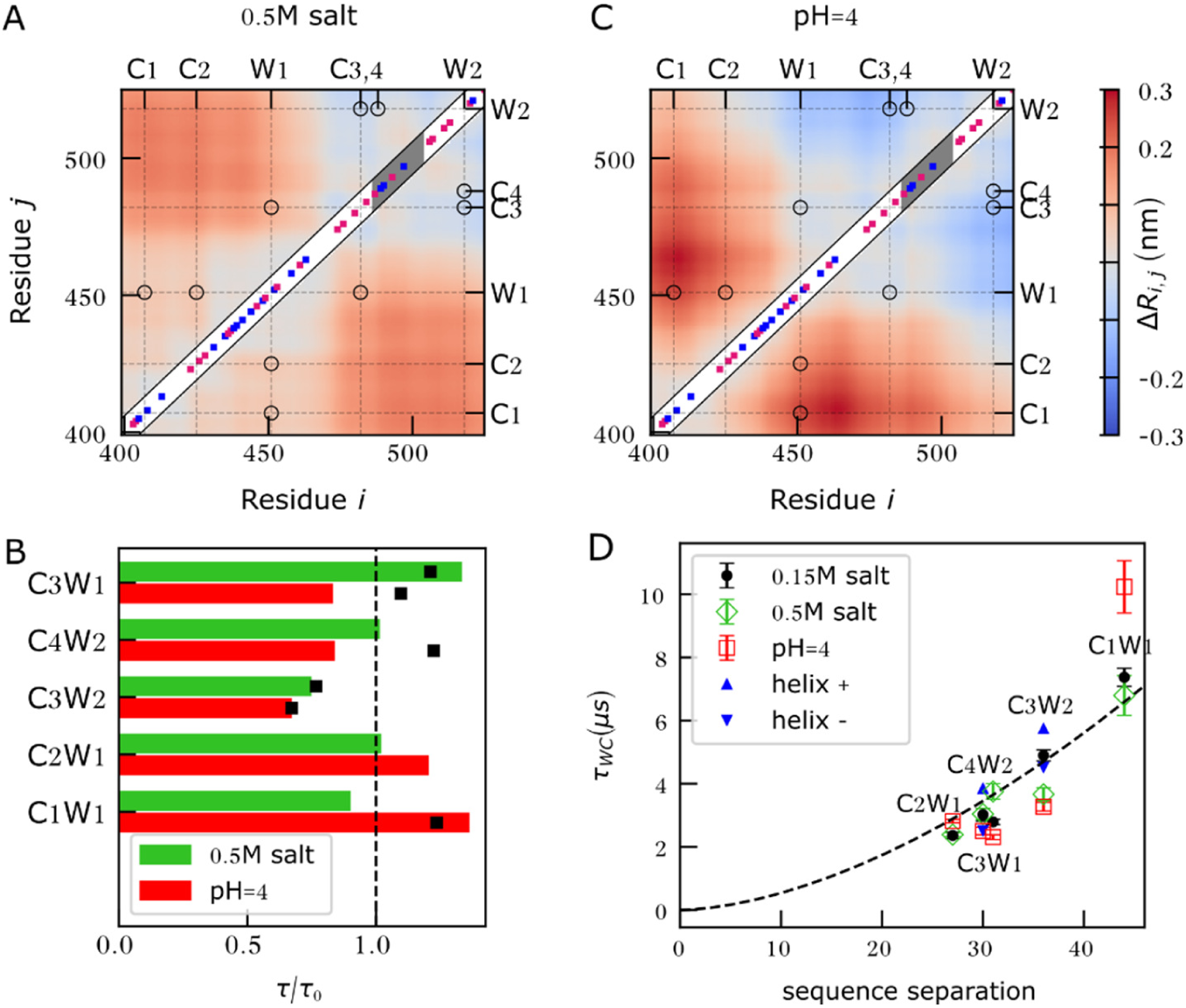
Coarse-grained (CG) simulation of NTAIL. Differences between the mean pairwise distances (Ri,j) of the CG simulations at the reference condition (0.15 M salt and pH 7.6) and simulations A) at high salt concentration (0.5 M salt, pH 7.6) or C) low pH (0.15 M salt pH 4.0), respectively. The diagonals show the charged amino acid positions with positively charged amino acids in blue and negatively charged amino acids in magenta. The MoRE is highlighted in grey. The full amino acid sequence is provided in Fig 1A. B) Ratios of C-W relaxation times due to PET, calculated from CG simulations (bars) at different conditions (high salt: green, or low pH: red), relative to the reference condition. For comparison, experimental ratios are shown where available (black squares). D) C-W relaxation times for all five NTAIL PET variants estimated from CG simulations at different conditions (reference: black, high salt: green, low pH: red). The blue triangles denote the relaxation times for the C3W2 and C4W2 variants after varying the MoRE helix content (helix-35%, helix+: 60%). All CG relaxation times were scaled by a factor, chosen so that the same homopolymer fitting function (dashed line) of Fig 1D fit both experimental and CG data.

The positively and negatively charged amino acids are well-blended in the N-terminal region, with a close to zero net charge. The N-terminus therefore behaves like a polyampholyte, where interactions between oppositely charged residues result in compact conformations and screening of these electrostatic interactions at higher salt concentrations results in more expanded conformations. This expansion in turn increases the average distance between C3 and W1, leading to a slower expected relaxation time at high salt concentration. In contrast, the C-terminal region has more negatively charged amino acids than positively charged amino acids under physiological conditions. This imbalance leads to net repulsive electrostatic interactions that keep this region expanded. Increasing the electrostatic screening at high salt concentrations reduces the repulsion, which leads to the compaction of the C-terminus of N_TAIL_, a decrease of the mean C3-W2 distance, and an expected decrease in the C3W2 relaxation time.

To determine if the salt-induced changes in relaxation times can be fully described by changes in inter-residue distances due to charge patterning alone, we estimated the expected ratio of relaxation times (1⁄1_0_) from the CG simulations at 500 mM and 150 mM NaCl concentrations, respectively. For these calculations, we assumed reaction-limited quenching kinetics, in which case, the relaxation time is inversely proportional to the probability of W and C being at a reactive distance. We used a cutoff distance of 1 nm to determine the probability of reactive conformations (further details in Supporting Methods S5). Figure 2B shows the 1⁄1_0_ ratios upon increasing the salt concentration for each variant (green bars). A ratio larger than one indicates an increase of the relaxation time, exclusively due to the expansion of that segment. Comparison with experimental ratios (Fig. 2B, black squares) shows that the CG simulations accurately predict both the increase in C3W1 relaxation times and the decrease in C3W2 relaxation times, which indicates that charge patterning suffices to explain the salt-induced changes in N_TAIL_ relaxation times. It also corroborates the hypothesis that electrostatic interactions between charged residues are a key factor in the observed dynamic heterogeneity in N_TAIL_.

To further test the effect of electrostatic interactions on the dynamic heterogeneity of N_TAIL_, we measured C-W relaxation times, upon decreasing the pH from 7.6 to 4.0, while maintaining a near-physiological salt concentration (150 mM) (Fig. 1D, red squares). In contrast to increasing the salt concentration which screens all electrostatic interactions, decreasing the pH only alters the charge on specific residues that titrate in the 7.6-4.0 pH range, specifically negatively charged Asp and Glu may become neutral, and neutral His become positively charged. If electrostatic interactions involving these charged residues were the main contributor to the dynamic heterogeneity observed at physiological pH, we would expect a reduction in heterogeneity when decreasing the pH to 4.0. In particular, we expect the N-terminal part, which acts as a balanced polyampholyte at pH 7.6, to become positively charged and therefore to expand at pH 4.0, even beyond what was observed at high salt concentrations. This expansion should then increase the PET relaxation times for the C1W1 and C2W1 variants. Conversely, we expect the negatively charged C-terminal part to become more neutral, resulting in more compact conformations that reduce the C3W2 and C4W2 relaxation times.

To capture the effects of lower pH in CG simulations, the charges of the residues that titrate between 7.6 and 4.0 were adjusted to match the average level of protonation at the simulated pH. To that aim, the charges assigned at pH 7.6 (−1.0 for Asp and Glu and +0 for His residues) were changed to partial charges at pH 4.0 (−0.69 for Asp, −0.36 for Glu and +0.99 for His). Further details on charge assignments are provided in Supporting Methods S5. If electrostatic interactions involving specific charged residues drive the dynamic heterogeneity at physiological pH, lowering the pH to 4.0 should reduce this heterogeneity. The N-terminal region, a balanced polyampholyte at pH 7.6, is expected to become positively charged and expand, while the negatively charged C-terminal region should become more neutral and compact. The CG simulations with simple fixed-charge adjustment at low pH are consistent with the expected behaviors described above, i.e. the N terminal region expands and the C terminal region becomes more compact (Fig. 2C), which is also reflected in the calculated relative relaxation times (red bars Fig. 2B).

The C-W relaxation times of all five PET variants estimated from measurements at pH 4.0 are closer to the homopolymer curve (Fig. 1D, red squares) than those at pH 7.6. Lowering the pH also reduces the observed dynamic heterogeneity between the C3W2 and C4W2 variants, with the ratio of relaxation times reduced from 2.8 at pH 7.6 to 1.5 at pH 4.0, which is closer to the homopolymer prediction of 1.3. However, in contrast to the reduction of the C-W relaxation time expected from compaction based on charge patterning and predicted by CG simulation, the measured relaxation time for C4W2 increases slightly upon lowering the pH (Fig. 1D).

In Fig. 2B the ratios of relaxation times at pH 4.0 and pH 7.6 estimated from the CG simulations (red bars) were compared to those estimated from PET experiments (black squares). The CG model correctly captures the trend of C1W1 and C3W2, but not that of C4W2 and C3W1. Further, the measured increase of r_CW_ for C4W2 is not only contrary to the CG simulation results, but also contrary to the expectation based on charge patterning, suggesting that other unaccounted factors influence the relaxation times. In the following, we will consider three factors that can affect relaxation times in a pH dependent manner: the MoRE helicity, W to C electron transfer efficiency, and residue specific changes in protonation equilibria.

#### pH-dependence of MoRE helix content

The CG model suggests limited difference in helical propensity upon changing pH (Supporting Fig. S20). We further measured and compared the far-ultraviolet circular dichroism (CD) spectra of the N_TAIL_ variant Y518W (W2) at pH 7.6 (Fig 1C, inset, black curve) and 4.0 (red curve). CD spectroscopy in this wavelength range of 180-260 nm is highly sensitive to the presence of α-helices and the two spectra appear to be very close to each other, indicating that the helical content of N_TAIL_ is not affected significantly by pH. In addition, analysis of the measured spectra by the SESCA software^45^ (Supporting Methods S13) predicts at most 8% global helix content, which is compatible with the NMR estimates used to set the MoRE helicity in CG simulations. Together, these results suggest the MoRE helix content is not affected by the change in pH.

#### Changes in electron transfer efficiency

Because PET involves formation of W radical cations, the pH may directly affect the electron transfer process in solution, independent of pH-induced changes in the protein conformational ensemble and dynamics. To address this possibility, we performed a bimolecular quenching study between N-acetyl-tryptophan-amide (NATA) and cysteine molecules, freely diffusing in solution, at pH 7.6 and 4.0 (Supporting Methods S2). In these experiments the bimolecular quenching rate was determined by measuring the W triplet relaxation times in samples at fixed NATA concentration and increasing cysteine concentrations. These measurements showed that the C-W bimolecular quenching rate (determined from the slopes of Stern-Vollmer plots) at pH 7.6 and 4.0 are very similar (Fig. S19). This result shows that the efficiency of electron transfer from the excited state W to C, under the conditions of our PET experiments is not directly affected by pH, and the pH dependence of r_CW_ in N_TAIL_ is due to changes in the protein structure and dynamics.

#### Residue-specific protonation

N_TAIL_ is rich in Asp and Glu, amino acids that reach protonation equilibrium close to pH 4.0. Under these conditions, the protonation state of individual acidic residues can be influenced, e.g., by the spatial vicinity of similarly or oppositely charged residues, resulting in a dynamic, conformation-dependent charge distribution.^46^ This charge distribution would also allow electrostatic interactions to stabilize certain conformational states, increase or decrease the observed r_CW_ for individual N_TAIL_ variants compared to those predicted by our fixed-charge CG simulations.

### Dynamic heterogeneity at the N_TAIL_ C-terminus

Although dynamic protonation may explain deviations from CG and homopolymer models, this effect is an unlikely explanation for the dynamic heterogeneity between C3W2 and C4W2 under reference conditions at pH 7.6. To address this heterogeneity with CG simulations, we assumed these processes are reaction limited, and that r_p_ the proportionality between C-W contact probabilities (P_rij<rO_) and corresponding r_CW_ relaxation times are the same for all PET variants, and thus r_ij_ = r_p_⁄P_rij<rO_. Under these assumptions, it is possible to compare relaxation times of different C-W pairs obtained from CG simulations relative to one another and address the dynamic heterogeneity at the N_TAIL_ C-terminus.

To this aim, we fitted a homopolymer model to these CG-derived relaxation times at the reference condition, and chose r_p_ such that the scaling equation is identical to the one obtained from experimental measurements in Fig. 1D. The resulting relaxation times from CG simulations for all three conditions (reference, high salt, and low pH) are shown in Fig. 2D. The figure shows that all CG-derived relaxation times at the reference condition (black circles) follow homopolymer scaling (dashed black line), with only small deviations for individual CW pairs. Although the CG simulations accurately predict many of the changes upon increasing salt concentrations or lowering the pH, they do not show either the considerable slowdown for the C3W2 relaxation time, or the pronounced dynamic heterogeneity measured at pH 7.6 between C3W2 and C4W2 (ratio 2.8). In fact, the CG-derived r_C3W2_/r_C4W2_ ratio of 1.6 is close to the 1.3 ratio predicted by homopolymer models, suggesting that the CG simulations do not capture the main cause of the dynamic heterogeneity observed in PET measurements.

An additional reason why the CG simulations may not have captured the C3W2/C4W2 dynamic heterogeneity is that we underestimated or overestimated the MoRE helix propensity (∼45%) under the conditions of PET experiments. This can alter the CG-derived heterogeneity if the MoRE helix in its folded state affects C3W2 and C4W2 relaxation differently. To test this hypothesis, we performed additional CG simulations in which we altered dihedral potentials that control the MoRE helicity and computed the r_CW_ for C3W2 and C4W2 variants at high (60%) and low (35%) helix contents (blue triangles on Fig. 2D). Relaxation times from these simulations show that considerable changes in the helix content can indeed shift the relaxation times obtained from CG simulations, but this shift affects both C3W2 and C4W2 similarly. Therefore, even a significant error in the imposed MoRE helicity would have only a small (±0.1) effect on r_C3W2_/r_C4W2_ ratio, certainly not sufficient to explain the dynamic heterogeneity observed in PET experiments.

Overall, our results show that N_TAIL_ dynamics is heteropolymeric in nature, especially in the region encompassing the MoRE, with electrostatic interactions playing an important role. It appears that the dynamic heterogeneity of N_TAIL_ cannot be explained by a simple polymer scaling model (Fig. 1D) or by CG models including helix formation within the MoRE and explicit interactions between charged amino acids (Fig. 2). These observations suggest that the cause of the dynamic heterogeneity is either side-chain specific transient interactions, such as salt-bridges and hydrogen bonds^47^, or interactions between the protein and solvent. To study the effects of these interactions on N**_TAIL_** dynamics in more detail, we turned to all-atom explicit-solvent molecular dynamics simulations.

### Disordered N_TAIL_ is characterized by a conformational ensemble with distinct states

In addition to specific interactions between amino acids and protein-solvent interactions, all-atom simulations include the backbone and side-chain dynamics required to accurately compute the contact formation rates. The relaxation times r_CW_ estimated from these contact formation rates can be compared to r_CW_ derived from PET experiments. However, this increased level of detail comes with two drawbacks; considerably higher computational costs that limit conformational sampling and the need for a larger number of accurate interaction parameters (force field). IDP simulations are particularly sensitive to variations in the used force field^48–51^, and force field accuracy for IDPs often varies in a system-dependent manner^52^.

To maximize conformational sampling, we limited all-atom simulations to the reference conditions (pH 7.6 at 150 mM NaCl concentration) and focused on the C3W2 and C4W2 variants, which showed large dynamic heterogeneity (Fig. 1D). Further, to ensure the reliability of all-atom simulations, we first validated eight force fields, characterized by different combinations of protein and water parameters, against independent experimental SAXS, CD, and NMR^6,16^ data on the wild-type (WT) N_TAIL_ sequence (shown in Supporting Table S3). To evaluate these force fields, we first performed simulations of WT N_TAIL_ using each force field, and then compared the resulting trajectories to the SAXS data, which reports on protein compactness, using the software CRYSOL^53^. Next, we compared the trajectories to CD and NMR chemical shift measurements, which report on secondary structure propensities, using SESCA^45^, and SPARTA+^54^. For further details of the force field validation, see Supporting Methods S8.

The two force fields that reproduced the WT validation data most accurately were the CHARMM36m (C36M)^51^ force field with optimal point charge (OPC) water model^55^, and Amber99SB-disp (A99SB-d) force field^49^. Both of these force fields produced extended N_TAIL_ ensembles with transiently forming helices in the MoRE region, that were in good agreement with the available experimental data (Supporting Table S3), and suggested helix propensities similar to those reported in computational studies of MoRE region^19,26^ (Supporting Methods S23).

The two best force fields were used to perform longer all-atom simulations of the C3W2 and C4W2 variants. As a second force field validation step (Supporting Table S4), the variant trajectories were tested against PET relaxation measurements of the same variants, and the CD spectrum of the W2 variant (Fig. S21B). Simulations of both force fields estimated C3W2 and C4W2 relaxation times showing considerable dynamic heterogeneity. However, the C36M-OPC force field was the only parameter set that reproduced measurements of both WT N_TAIL_ and the two variants within the experimental and computational uncertainty, including the two measured W relaxation curves (Supporting Fig. S22). Thus C36M-OPC trajectories were used for all subsequent analysis. The relaxation times r_CW_ estimated from these simulations were 2.0 ± 0.5 μs for C4W2 and 5.3 ± 3.3 μs for C3W2, in good agreement with the corresponding measured values of 2.3 μs and 7.6 μs. Although the uncertainty on the C3W2 relaxation time is large due to limited sampling, the ratio of the relaxation times estimated from the simulations is 2.7 ± 1.8 close to the experimentally determined ratio of 2.8. Together, these results suggest that the all-atom simulations accurately describe the N_TAIL_ dynamics causing the C3W2/C4W2 dynamic heterogeneity.

To study the origin of the dynamic heterogeneity in detail, we computed the free energy landscape of the N_TAIL_ variants along the C-W distances for positions C3W2 and C4W2. Variants C3W2 and C4W2 only differ in the exchange of an oxygen and a sulfur atom between two nearby residues at position 482 (C3) and 488 (C4), respectively. We therefore assumed that the free-energy landscape of the two variants is similar and combined their trajectories for computing it. The C-W distances were defined as the distance between the gamma sulphur of C (or gamma oxygen of S) and the nearest heavy atom of the W side-chain ring. These serve as intuitive reaction coordinates for describing C-W contact dynamics in the N_TAIL_ C-terminal region.

Unexpectedly, the computed free-energy landscape (Fig. 3) is characterized by four local minima corresponding to four N_TAIL_ conformational states (S1 to S4). Approximately 60% of the sampled N_TAIL_ conformations are within the respective energy wells, including almost all conformations with C-W contacts necessary for PET. S1 corresponds to the conformational state around the absolute free-energy minimum and is occupied for approximately 24% of the simulation time. S2 and S3 are less populated conformational states, occupied for 14% and 8% of the simulation time, respectively. S4 is the least populated state, with approximately 2% of conformations. The landscape shows only small barriers (< 1 kT) separating S1, S2, and S3, and a low free-energy barrier between S1 and S4 (< 2.0 kT). This shallow and accessible free energy landscape suggests nearly-free diffusion between states, which is in line with previous SAXS results suggesting N_TAIL_ to be a pre-molten globule with residual structure^6^.

**Figure 3.**
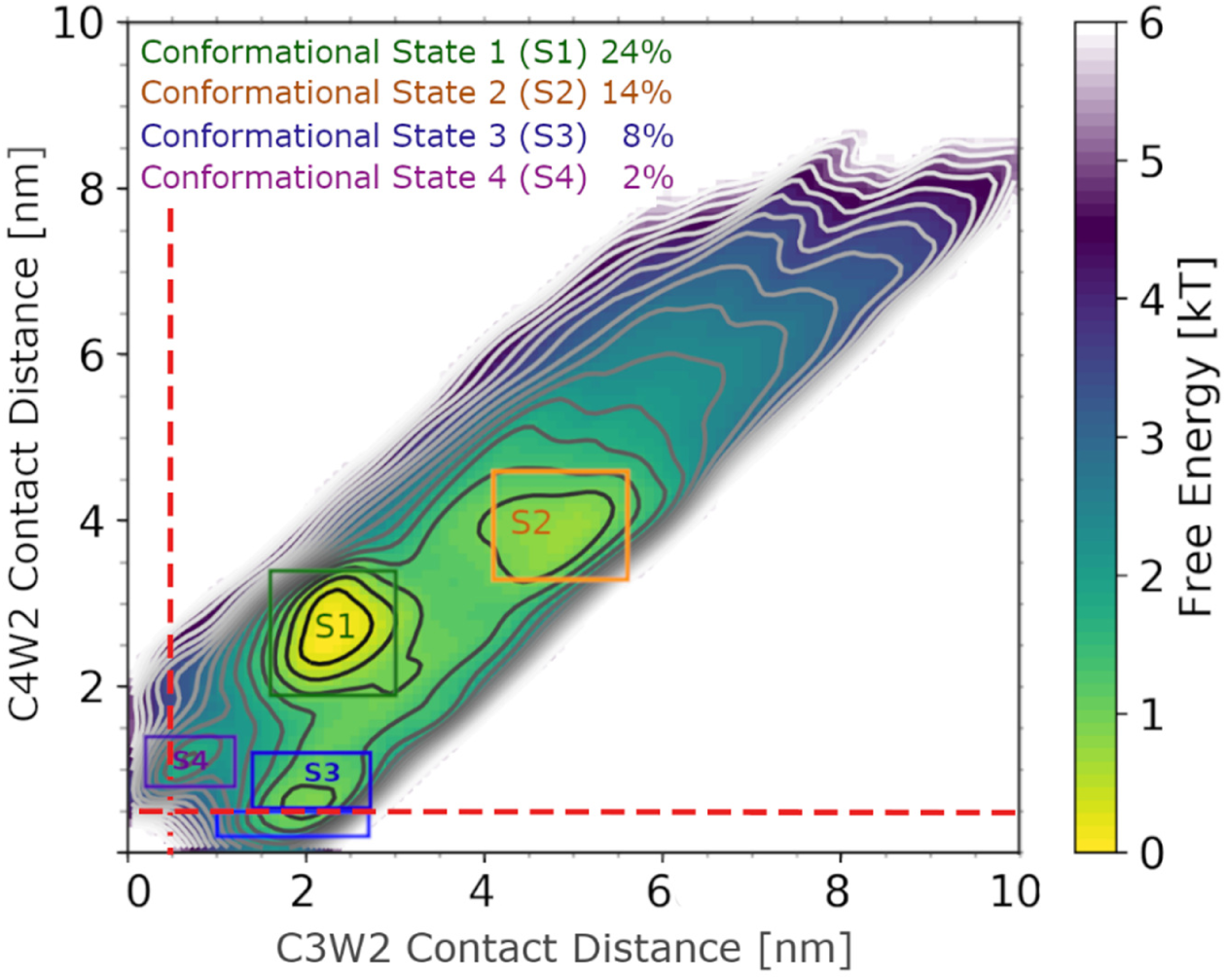
Conformational states revealed by all-atom MD simulations. The free-energy landscape of sampled NTAIL conformations along the calculated C-W distances from C36M-OPC MD simulations. Contour lines denote a 0.3 kT difference in the estimated free energy landscape. Conformational states are highlighted by colored labels and rectangles. The population of conformations within each rectangle is shown in the legend. Red dashed lines indicate a contact distance of 0.4 nm. Conformations below these cutoffs would allow quenching of the W triplet state, due to electron transfer to C, for the corresponding variant (C4W2 or C3W2).

The four conformational states influence C-W relaxation times as well. PET cannot occur in the two most populated states (S1 and S2), because the distances between electron acceptors C4 or C3 and electron donor W2 are more than 2 nm. In state S3, C4-W2 distances below 0.4 nm are observed, allowing PET in the C4W2 variant, while in state S4, C3-W2 distances below 0.4 nm are reached, which allows PET in the C3W2 variant. Further, from the most populated S1 state the free-energy barrier that N_TAIL_ has to cross to reach S4 is about 0.9 kT larger than the barrier to reach S3. This suggests that S3 is more easily accessible than S4, with an approximately 2.5-fold difference between the respective transition rates. This result is consistent with and provides a possible explanation for the measured 2.8 ratio between the C3W2 and C4W2 relaxation times.

### Non-local N_TAIL_ interactions explain dynamic heterogeneity

To determine the molecular basis of the observed dynamic heterogeneity, we investigated which specific interactions stabilize the four conformational states identified above. To this aim, Fig. 4 shows intramolecular contact probability maps for each conformational state, with backbone (C_α_) contacts shown in yellow to green. Because both the pH and salt concentration dependence of PET experiments and CG simulations indicated an important role of electrostatic interactions in N_TAIL_ dynamics, we also calculated contact probability maps for salt bridges between charged amino acids shown in purple to blue. For the sake of clarity when discussing the contacts, we defined five regions along the N_TAIL_ sequence (Fig. 4 bottom, color-coded and labelled A-E). Regions A-C are subdivisions of the polyampholyte N-terminal part identified in our CG simulations, regions D and E constitute the negatively charged C-terminal part, with the MoRE located in region D.

**Figure 4.**
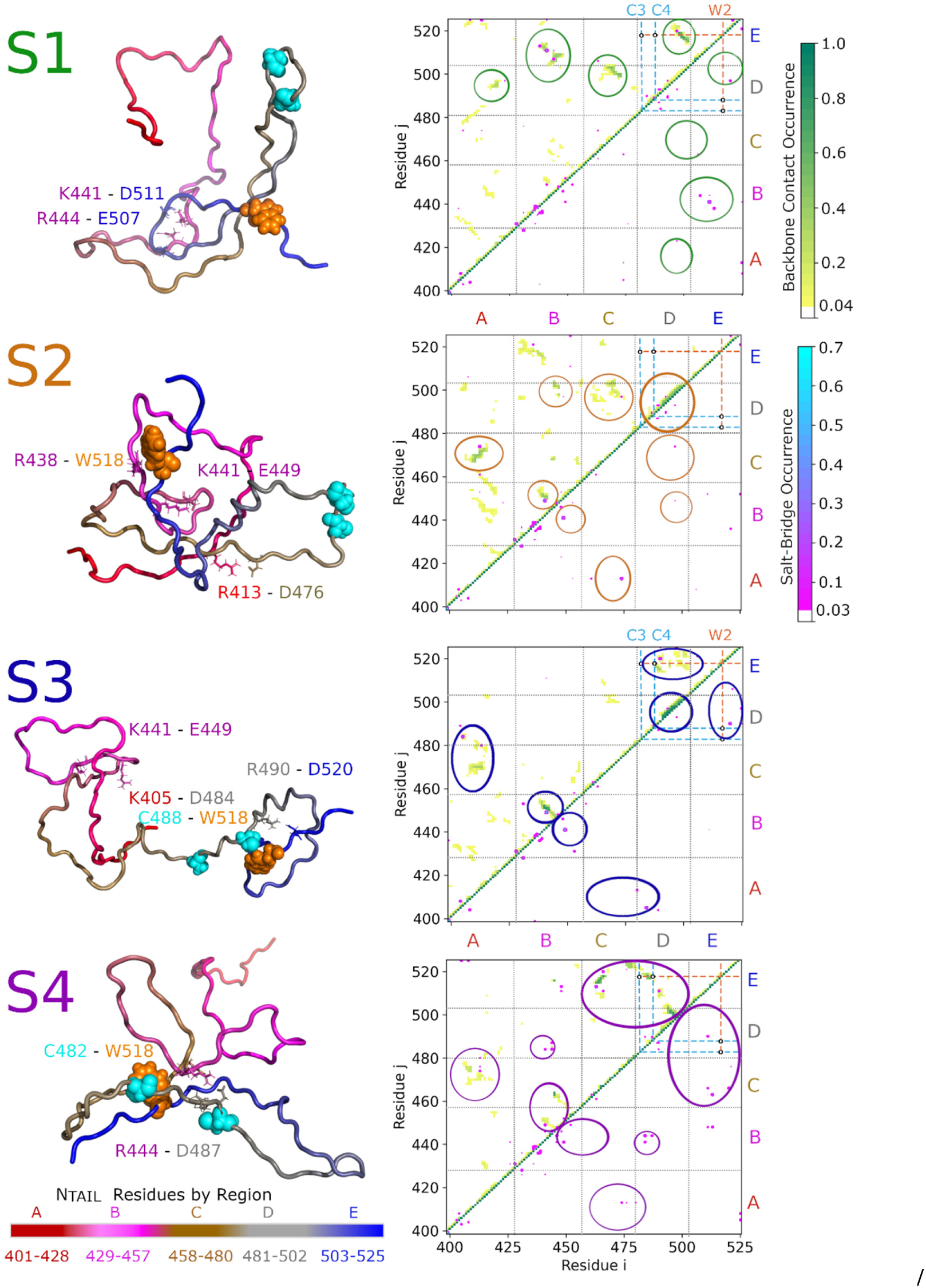
Contact maps and representative conformations in NTAIL conformational states. For each conformational state (S1-S4, shown in Fig.3), a representative conformation (left) and contact probability map (right) is shown. Each map shows Cα contact probabilities (above the diagonal, yellow to green) and salt bridge probabilities (above and below the diagonal, purple to blue). Cysteine acceptor positions C3 and C4 (residues 482 and 488) as well as tryptophan donor position W2 (residue 518) are indicated by cyan and orange dashed lines, respectively. Potential C-W contacts are indicated by black circles. The boundaries of identified interaction regions A-E and their color-coding for representative conformations are shown in the bottom-left side of the figure. The same regions are separated by black dotted lines on the contacts maps. The most prominent interactions in each state are encircled. The pixel size of salt-bridge interactions was increased for visibility, using a Bessel interpolation scheme. On the left side, the backbone trace of a representative conformation from each state is shown in cartoon representation, colored according to regions. The donor and acceptor atoms are highlighted as orange and cyan spheres, respectively. Residues with prominent interactions in the conformational states are labelled and shown in stick representation. Individual contact maps in high resolution, as well as the average contact map for all NTAIL simulations are shown in Figs. S23-S27.

The contact maps (Fig. 4, right) clearly show that the four states are stabilized by distinct sets of local (within a region) and non-local (between regions) interactions, although some interactions occur in several conformational states. The maps also provide information on the secondary and tertiary structure preferences of N_TAIL_ in these conformational states. Helical secondary structure elements and tight turns appear on contact maps as continuous sets of interactions, parallel and close (within 5 residues) to the diagonal. β-strands, β-hairpins or longer loops appear as sets of interactions that are further away and either parallel or perpendicular to the diagonal. Disordered regions typically have few or sporadic interactions.

The four conformational states show markedly different structures (Fig. 4, left), including the MoRE in region D. In state S1, region D is largely non-helical but forms several transient local salt bridges. Several non-local interactions (encircled in green) may prevent donor-acceptor contacts for both variants (C3W2 and C4W2). S1 conformations are also stabilized by salt bridges formed between regions B and E. Salt bridges involving K441, R444 (region B) and E507 and D511 (region E) are particularly frequent and probably play an important role in stabilizing this conformational state.

State S2 includes conformations where region D is non-helical and conformations with a folded α-helix (residues 492-502). The helical structures appear on the contact map as an interaction patch close to the diagonal of region D (encircled in orange). Non-local interactions in S2 appear to be less stable than in S1 and include contacts between regions A and C, C and D, as well as B and D. Stable interactions in S2 also involve two salt bridges between R413 and D476 (regions A and C) and K441 and E449 (local in region B). Many sporadic interactions (both backbone contacts and salt bridges) were observed within the N-terminus (between regions A and B). Taken together, the interactions suggest that S2 consists of multiple conformational substates and most interactions are linking central regions (B and C) with both termini (A, D, and E).

Compared to S1 and S2, the conformations of S3 comprise fewer non-local interactions and are instead characterized by two very stable local interaction sites. These local interactions correspond to a shorter helix between residues 492-497 present in all S3 conformations and to a stable hairpin or zipper in region B, stabilized by a salt bridge (K441-E449) observed previously in S2. Further, there are two sets of notable non-local interactions. First, a patch of interactions between regions D and E folds the C-terminus into a loop, stabilized by the salt bridge between R490 and D520. These interactions also bring the electron donor W2 (518) very close to the potential acceptor at position C4 (488), thus promoting electron transfer in the C4W2 variant. At the same time, the zipper in region B brings the N-terminal close to regions C and D, where a salt bridge between K405 and D484 may prevent electron transfer to position C3 (482) in the C3W2 variant.

The S4 state allows electron transfer to position C3 (482) in the C3W2 variant. It is structurally very different from states S1-S3. The most defining structural feature of this state is a long loop reminiscent of an anti-parallel β-sheet stretching over the entire C-terminus (regions D and E). This loop is stabilized by interactions between regions C and E including a salt bridge between R463 and D513. The N-terminus of N_TAIL_ in this conformational state has only weaker interactions, mostly limited to contacts between regions A and C as well as between regions B and C.

To determine if frequent contacts observed in one of the four conformational states (Fig. 4) are specific to that state and if they are correlated with other contacts, including C-W contact formation for the C3W2 and C4W2 variants, we applied normalized pointwise mutual information analysis (NPMI, see Supporting Methods S22).

The NPMI analysis revealed that many salt bridges between regions B, D, and E occur selectively in their respective conformational states. These stable salt bridges often co-occur with clusters of backbone contacts (encircled in Fig. 4), possibly nucleating the interaction clusters that stabilize the states.

Further, all frequent contacts observed in the S1 state were anti-correlated with C-W contact formation (Table S5), indicating that the specific interactions that stabilize the S1 state would increase the expected relaxation times compared to the flat free-energy landscape of a homopolymer model. In contrast, all frequent contacts of states S3 and S4 are positively correlated with C-W contact formation for variants C4W2 and C3W2, respectively. Thus, interactions in these two states selectively reduce the relaxation times of either C4W2 or C3W2 variants, contributing to the dynamic heterogeneity observed in PET experiments.

Finally, the same charged residues (e.g., K441, R497, and D501) in regions B, D, and E form salt-bridges with different residues in different conformational states (Table S5). Switching between these sets of interactions is necessary to transition between the most populated conformational states, and therefore likely define the free-energy barriers between the states (Fig. 3). Further, a conformational switch based on these competing intramolecular N_TAIL_ interactions of region B may have a functional role, by regulating the helical propensity of the MoRE in region D, which in turn modulates affinity for the measles phosphoprotein.

### Functional relevance of non-local interactions is supported by coevolutionary analysis

Having identified the most important interactions that dictate N_TAIL_ structure and dynamics (Fig. 4), we asked whether they also have a functional role. To this aim, we performed a coevolutionary analysis of N_TAIL_ using EVcouplings^56^, which identifies and quantifies correlated mutations of residue pairs within the sequence of homologous proteins from different species or strains, and which often indicates pairwise-interactions of functional importance. A total of 12,072 sequences similar to N_TAIL_ have been aligned and selected for this analysis (Supporting Methods S21).

Most non-local residue pairs with positive EVcouplings scores connect regions B and D (Fig. S28), suggesting evolutionarily conserved interactions between these two regions. Here we specifically highlight the four non-local residue pairs with the highest EVcoupling scores (Fig. S28, top left, Table S5). These pairs include hydrophobic contacts involving three leucine residues (L491, L496 and L498) interacting with S443 and A447, and one salt bridge between R444 and D493. Correlated mutations of residues in the distant regions B and D suggest a biologically relevant, functional interaction between these two regions of N_TAIL_. Given that both L491 and L498 are located in the MoRE region (in region D) and are directly involved in binding to P_XD_, it is possible that their interactions with residues of region B serve the function of regulating binding. We note that the contact analysis of our simulations does not identify these specific coevolving residue pairs as frequent interactions. However, all-atom simulations independently identify non-local interactions between regions B and D (Fig. 4, S2 state), including A447 contacts with P_XD_ binding residues L498 and M501. These differences between all-atom simulations and the co-evolution analysis may be due to insufficient sampling or the lack of biological context in all-atom simulations (i.e. the absence of the N_CORE_ domain or P_XD_ itself).

## Discussion

To understand the role of disordered regions flanking the MoRE on either side in N_TAIL_ dynamics and conformational preferences, we probed non-local interactions of the full-length protein in solution, combining photo-induced electron transfer experiments, analytical polymer models, molecular dynamics simulations, as well as coevoluionary analysis. With PET experiments, we measured and compared contact formation times between tryptophan and cysteine residue-pairs introduced in different regions of the N_TAIL_ sequence, under varying solution conditions. The experiments revealed a pronounced dynamic heterogeneity within the protein (Fig. 1) under physiological conditions, especially at the N_TAIL_ C-terminal region, close to the MoRE, which is involved in binding to P_XD_. PET experiments also showed a reduced heterogeneity under conditions that weaken electrostatic interactions, such as increased salt concentration or lower pH, pointing to an important role of electrostatic interactions between charged residues in the dynamics of N_TAIL_. Based on our molecular dynamics simulations, we attribute this heterogeneity to salt-bridge induced non-local interactions between specific regions of N_TAIL_.

### Interpretation of the general role of electrostatics

This finding is supported and explained in structural terms by our simulations (both coarse-grained and all-atom). Even without including atomic-level details, our CG simulations suggest that when increasing the electrostatic screening or changing the charge of acidic residues to emulate pH related protonation, the N-terminal region expands while the C-terminal region compacts. In addition, CG simulations performed with different helical propensities in the MoRE indicate that changes in helicity of the MoRE do not give rise to the dynamic heterogeneity within the N_TAIL_ C-terminal region (i.e. C3W2 and C4W2). These results indicate that side-chain specific transient interactions, not described by the CG model, contribute to the large difference between the C3W2 and C4W2 relaxation times.

### Specific non-local interactions dictate N_TAIL_ conformation and dynamics

Our all-atom simulations, performed with a carefully validated force-field, describe interactions with atomic-level detail. These simulations reproduce the dynamic heterogeneity of C-W pairs encompassing the N_TAIL_ MoRE (C3W2 compared to C4W2). We found that, while N_TAIL_ is clearly disordered, a few key non-local interactions stabilize four distinct conformational states which in total represent 48% of the structural ensemble. The two most populated states do not allow PET for either C3W2 or C4W2 variants, while the third and fourth states allow PET for the C4W2 and C3W2 variants, respectively. The free energy landscape of N_TAIL_ (Fig. 3) reveals rapid interconversions between the three most populated states. In contrast, structural rearrangements required to reach the C3W2 contact state are slower due to a small, though comparatively higher free-energy barrier (∼4.4 kJ/mol) under physiological conditions. This free-energy barrier and the lower equilibrium population of the C3W2 contact state contributes to the dynamic heterogeneity of the N_TAIL_ C-terminal region.

Analysis of the state-specific contacts (Fig. 4) reveals that competing clusters of interactions stabilize the four conformational states. These interaction clusters involve (and are possibly nucleated by) stable salt bridges between charged amino acid side chains. Further, focusing on the two most populated conformational states S1 and S2, our all-atom simulations show that specific charged residues in region B (residues 435-451) can form either non-local salt bridges with C-terminal regions D-E (484-525, close to the MoRE), or local salt bridges with other region B residues. Switching between these sets of interactions (as sketched in Fig. 5) is necessary for N_TAIL_ to transition between the two dominant states, affecting its large-scale dynamics.

**Figure 5.**
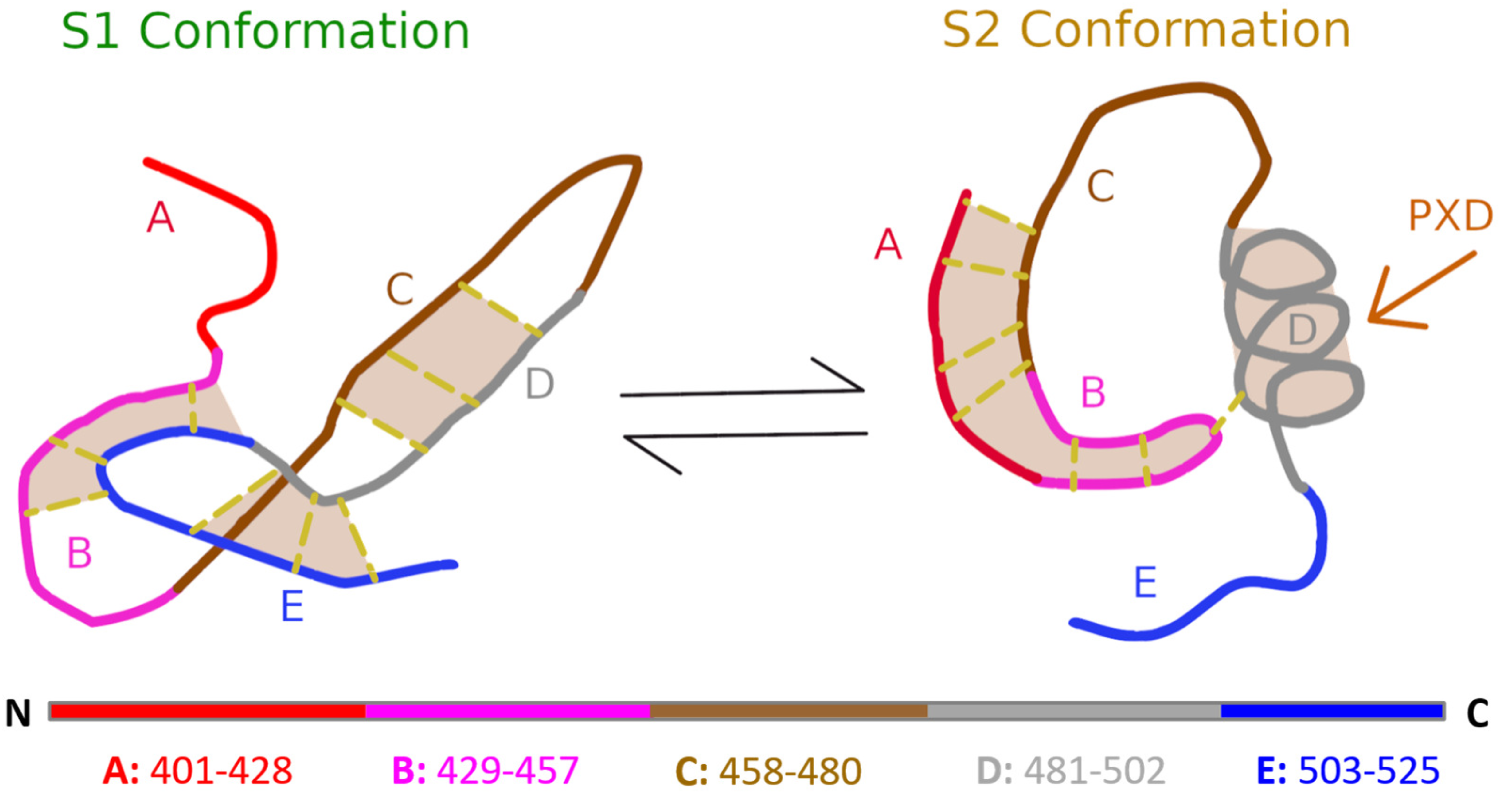
Schematic representation of the two major NTAIL conformational states (S1 and S2). Different regions of NTAIL are color-coded according to Figure 4. Non-local interactions between the regions are represented as dashed lines. Key interactions between regions are highlighted by orange shading. The helical binding site of the phosphoprotein X-domain (PXD) in region D is marked by an orange arrow.

Co-evolutionary analysis independently shows that correlated mutations across regions were only found between regions B and D. Given the distance in sequence between these two regions, this not only corroborates the presence of non-local interactions between region B and D, but also suggests that they play a functionally important role.

In summary, our simulations and coevolutionary analysis independently highlighted region B of N_TAIL_ as a potential allosteric interaction partner to region D, which includes the binding site (MoRE) for the phosphoprotein X-domain (P_XD_). Additionally, fluorescence-based binding kinetics experiments on truncated N_TAIL_ variants^23^, indicated that intramolecular interactions of N_TAIL_ involving residues 435-451 (region B) have a significant impact on P_XD_ binding in vitro, despite being located far away in sequence from the MoRE.

### Identified interactions in the context of N_TAIL_ to P_XD_ binding

We hypothesize that the intramolecular N_TAIL_ interactions observed in our simulations not only govern the N_TAIL_ dynamics but also play a role in regulating P_XD_ binding to the MoRE. Mutational studies by Bignon *et al.*^57^ have shown that the helix content of the MoRE correlates with the binding affinity, and that N-terminal truncation of N_TAIL_ increases its affinity for P ^22,24,27^. Similarly, Gruet *et al.*^27^ have identified mutations in several regions of the N sequence, on either side of the MoRE, which increase P_XD_ binding affinity. The crystallographic structure of the N_TAIL_ MoRE bound to P ^9^ (PDB: 1t6o, Fig. 6B) shows that the binding interface consists mainly of a hydrophobic edge along the MoRE helix (region D), between residues S491 and M501 (hereafter D1-site, shown in dark green), which interacts via intermolecular contacts with the α2-α3 face of the P_XD_ triple helix (Fig. 6A). The other side of the MoRE helix is a hydrophilic edge formed by charged and polar residues (D2-site, dark red) between Q486 and R497. The N_TAIL_-P_XD_ complex is further stabilized by intermolecular interactions involving R497 and salt bridges formed by D487 and R490 which are located at the beginning of the MoRE helix between the two edges (D3-site, light green)^27^. Many of these key amino acids are involved in intramolecular N_TAIL_ interactions in one or more conformational states, as shown by the contact probabilities obtained from all-atom simulations (Fig. 4).

**Figure. 6.**
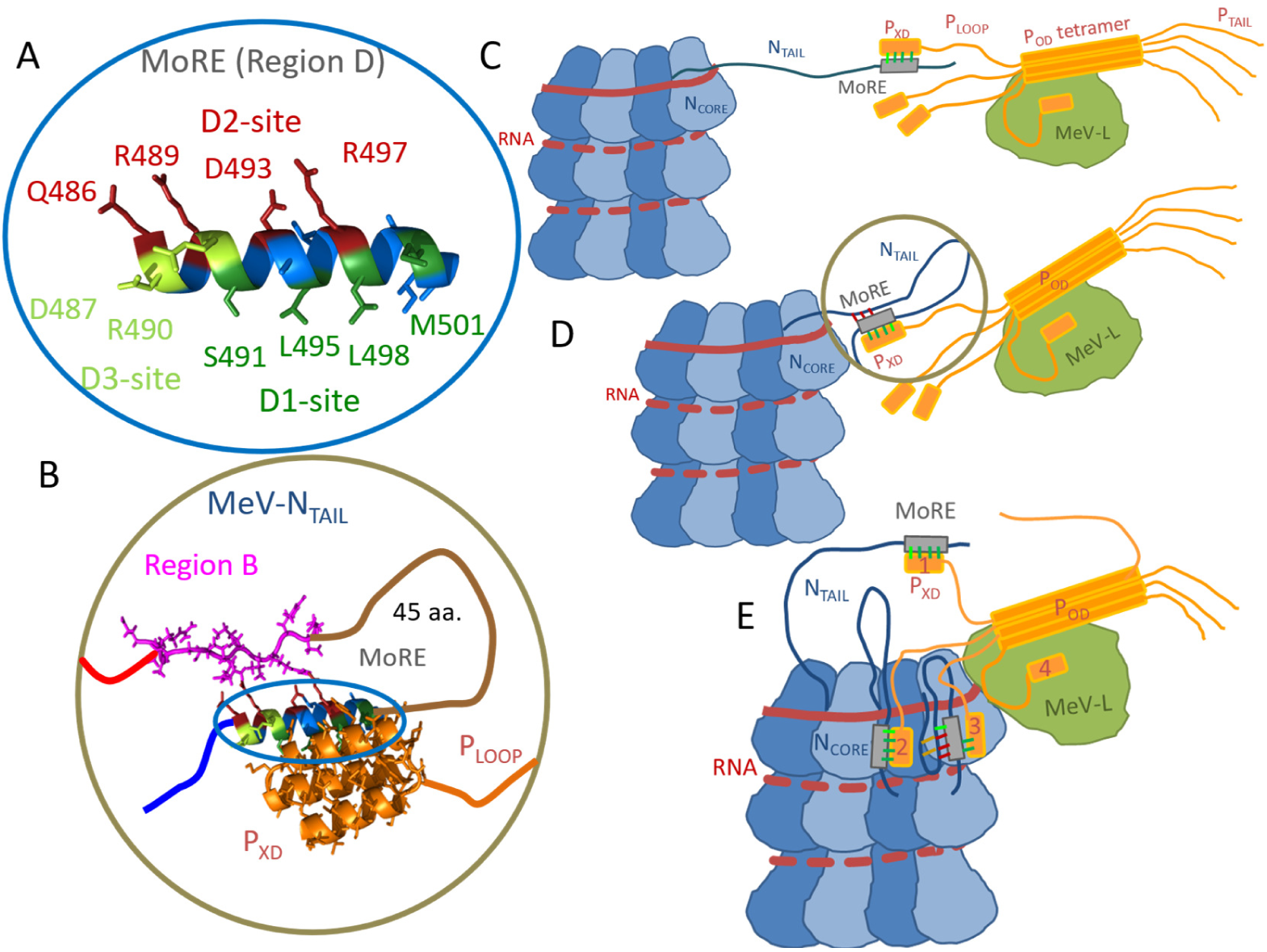
Speculated biological context of non-local NTAIL interactions. (**A**) NTAIL MoRE residues in region D constituting the hydrophobic PXD-binding interface are highlighted (D1 site, dark green) along with charged residues possibly involved in intramolecular NTAIL interactions (D2 site, dark red), and residues possibly involved in both intra- and intermolecular interaction (D3 site, light green). **(B)** Cartoon representation of the PXD-MoRE complex based on the available crystal structure (PDB: 1t6o). **(C)** During the initial steps of the virus replication the MeV nucleocapsid protein (N, blue) recruits the large polymerase (L, green), by binding to the phosphoprotein (P, orange). Here NTAIL is extended. (**D**) When Brownian motion allows L to approach the RNA (red) in the nucleocapsid groove, NTAIL adopts a hairpin conformation, which promotes intramolecular interactions between the MoRE and the N-terminal region of NTAIL. **(E)** Once the contact between nucleocapsid and L is established, additional complexes can be formed between free PXD and downstream N monomers. Further, intramolecular NTAIL interactions help release bound P to allow polymerase progression, by weakening the binding to PXD when L is nearby. Numbers (1-4) on PXD denote which step individual domains are in a “cartwheeling” L progression mechanism. In panels C, D, and E, only 1-3 NTAIL domains have been depicted for the sake of clarity. The RNA molecule is shown as a solid line on its first helix turn around the nucleocapsid to indicate that it is more solvent exposed, and as a dashed line on downstream turns to highlight that it is partially buried in the RNA-binding ridge.

Based on these interactions, we propose two independent mechanisms that provide a possible explanation for the observed changes in P_XD_ binding affinity upon N_TAIL_ mutation or truncation.

#### Direct competition for MoRE

In all-atom simulations, the D1-site residues (Fig. 6A, dark green) form intramolecular interactions with residues of regions B and C in state S2 as well as with residues of region E in state S3 (see Supporting Table S7), both of which are states with considerable helical propensity of the MoRE region. Assuming that a folded MoRE helix is a necessary part of the biologically active N_TAIL_ conformation, these intramolecular interactions would directly compete with the intermolecular interactions with P_XD_.

First, the intramolecular interactions would reduce the free helix population of the MoRE, thereby decreasing the binding rate of P_XD_. Second, the intramolecular interactions may also displace P_XD_ from the binding site without exposing the hydrophobic D1 site to water, thereby lowering the P_XD_ unbinding activation barrier and increasing the unbinding rate. Both effects would regulate MoRE-P_XD_ binding by reducing the binding affinity, and explain why the binding affinity increases when flanking regions of the MoRE are removed^27,57^.

#### Indirect allosteric regulation

Helical conformations of the N_TAIL_ MoRE appear to have higher affinity for P_XD_^57^. Therefore, changing the population of helical MoRE conformations via intramolecular interactions would also affect the MoRE-P_XD_ binding affinity. This mechanism is similar to the ensemble allosteric model for IDPs described by White *et al*^29^. Specifically, intramolecular interactions that stabilize non-helical MoRE conformations, such as the contacts seen in state S1, would decrease the population of helical conformations, reducing both N_TAIL_ binding affinity and the rate of binding to P_XD_. The observed contacts (Fig. 4) suggest that the ratio of states S1 and S2 is controlled by the competing interactions of region B, which acts as a conformational switch. Specifically, in state S1, the interactions between regions B and E stabilize a loop-like, non-helical conformation of the N_TAIL_ C-terminus, which is likely binding-incompetent towards P_XD_. In contrast, in state S2, region B residues engage in local interactions, allowing the N_TAIL_ C-terminus to form the high-affinity helical structure in the MoRE at region D.

### Potential functional role of N_TAIL_ intramolecular interactions in viral replication

Non-local interactions between regions B and D/E of N_TAIL_ may also play a functional role in RNA polymerization and viral replication. In the following we present three established mechanisms where intramolecular interactions between the B and D regions of N_TAIL_ may contribute to optimizing the binding kinetics for viral replication.

#### Tether shortening

Initial polymerase complex recruitment likely involves the formation of one or more disordered tethers between the polymerase complex and the nucleocapsid. By limiting the volume available for diffusion, tethering increases the local concentration of the polymerase complex around the nucleocapsid, thereby increasing the rate of transcription/replication initiation. There are two experimentally established tethers between N and P. The first tether consists of the disordered N-terminal tail of P (1-304, P_TAIL_) binding to two opposite sides of N_CORE_, and its primary function appears to be the prevention of nucleocapsid association in the absence of the viral RNA^58^. The second, tether is formed by regions B and C of N_TAIL_ and the C-terminal domain (P_LOOP_, 376-459) of P, through coupled binding and folding between the MoRE of N and P_XD_ (Fig. 6C). This tether regulates L progression on the nucleocapsid.^59^ The two tethers may coexist and in fact work in concert to form N-P condensates that serve as the basis of viral factories within the host cell^60,61^, and to securely attach the polymerase complex to the nucleocapsid during transcription/replication initiation^11^.

During replication initiation, it is preferable to have long tethers to recruit polymerase complexes from a larger volume around the nucleocapsid, but later, when L has to be guided to the 3’ end of the RNA, shorter tether lengths may be preferable. Intramolecular N_TAIL_ interactions between the MoRE D2-site and region B would shorten the tether by 45 amino acids (Fig. 6D), further reducing the volume available to the polymerase complex, and increase the rate of initiation of genome transcription and/or replication.

#### P_XD_ unbinding during polymerase progression

Sourimant *et al.*^10^ carried out recombinant minigenome bioactivity assays found that polymerase progression is slowed down considerably by a deletion of regions B and C of N_TAIL_ and proposed a mechanism for polymerase progression. This mechanism is based on a newly discovered binding site on N_CORE_ for the α1-α2 face of P_XD_, at which compensatory point mutations could weaken P_XD_-N_CORE_ binding and restore bioactivity. Briefly, the mechanism suggests that the L-P polymerase complex is tethered to the nucleocapsid by forming the N_TAIL_ MoRE− P_XD_ complex. Then, the P_XD_-MoRE complex binds the recently discovered N_CORE_ site. P_XD_ binding to this N_CORE_ site likely facilitates P_XD_-MoRE unbinding, possibly after compaction of the N_TAIL_ loop to allow polymerase passage. P_XD_ unbinding in turn allows subsequent binding to MoRE of the downstream N monomer, and polymerase translocation.

The mechanism proposed by Sourimant *et al.* provides previously unprecedented molecular insight into the measles virus replication. However, it mostly considers the deleted regions of N_TAIL_ as passive linkers or “roadblocks” for L, and it does not give an in-depth explanation on how weakening the N_CORE_-P_XD_ binding affinity compensates for reduced bioactivity caused by the deleted N_TAIL_ segments. Our current results and previous binding efficiency studies on truncated N_TAIL_ constructs^22^ suggest that non-local interactions between MoRE and N_TAIL_ regions B and C can in fact actively compete with P_XD_ for its binding site and may do so in the context of a MoRE-P_XD_-N_CORE_ ternary complex as well. This implies that deleting regions B and C does not just shorten N_TAIL_ but also eliminates intramolecular interactions that promote MoRE-P_XD_ unbinding. This would explain why the deletion hampers viral replication, why mutations at the N_CORE_ interface compensate for the deletion, and provides a possible reason for region B residues to co-evolve with the MoRE (Table S5). In addition, based on the available cryo-EM structure of the nucleocapsid^1^ (PDB: 4uft), region B would emerge from the RNA binding ridge close to the N_CORE_ binding site, with sufficient space for region B to fold into a loop and interact with the MoRE region (Fig. S24).

#### Polymerase progression by cartwheeling

Our findings are also compatible with a modified version of the cartwheeling mechanism^62^ for polymerase progression (Fig. 6E). Here, P is recruited (Fig. 6E, step 1) by N_TAIL_ and bound to the N_CORE_ binding site (step 2) of the same monomer as described by Sourimant *et al.*^10^. However, in this mechanism P_XD_ remains bound to the same N monomer until the L polymerase approaches. Then, due to steric hindrance from L or conformational changes at the N-N interface, N_TAIL_ folds back onto the nucleocapsid (step 3) in a conformation that allows region B to weaken P_XD_-MoRE interactions. This, in turn, allows P_XD_ to unbind and be transferred to its binding site on L (step 4) as L progresses. Subsequently, P_XD_ is either released to the solvent or directly binds to the N_TAIL_ of a downstream N monomer in the next cartwheeling cycle.

As discussed above, evidence suggests intramolecular N_TAIL_ interactions weaken P_XD_ binding. Tight binding to P_XD_ is beneficial for tethering the viral polymerase to the nucleocapsid during polymerase recruitment and progression, and thereby ensures that the transcription and replication of the genome is completed. However, unbinding of individual P_XD_-N_TAIL_ complexes is required for polymerase progression during viral replication^63^. Therefore, MeV and related viruses may be at an evolutionary disadvantage if the binding between P_XD_ and N_TAIL_ is so tight that it limits the replication rate, as supported by mutational studies of P ^64^. It is thus tempting to hypothesize that a possible functional role of non-local intra-protein N_TAIL_ interactions is to provide a conformation-dependent optimization of P_XD_ binding affinity.

## Conclusions

Our combined experimental and simulation approach shows that N_TAIL_ is clearly disordered, contains very little secondary structure, and shows nearly diffusive dynamics. However, it still displays some structural features which cause its dynamics to deviate significantly from that of a homopolymer. In particular, we identified several key transient interactions between disordered regions distant in sequence as the main reason for this deviation. These interactions affect the overall conformation and dynamics of N_TAIL_ in solution. Interestingly, similar interactions also emerge from our independent co-evolutionary analysis, corroborating our simulation results and suggesting that they not only govern the conformational dynamics of the essential N_TAIL_ domain, but also are functionally important. We therefore propose possible mechanisms by which these non-local interactions regulate binding to the phosphoprotein X domain (P_XD_), and consequently, recruitment and progression of the polymerase complex onto the nucleocapsid template. Our results for N_TAIL_ corroborate the importance of flanking regions in IDPs that bind via short molecular recognition elements (MoREs)^64^, and suggest a regulatory role of regions that may be far in the sequence from MoREs. These non-local interactions within N_TAIL_ can regulate binding *via* an ensemble allosteric model^29,30^ without requiring the binding of a third molecule. Further studies targeting the N_TAIL_-P_XD_ interaction may provide additional insight into the underlying mechanisms and effects of these interactions.

It is plausible that similar mechanisms are also at work for other IDPs that share sequence characteristics, such as charge patterning, with N_TAIL_. Specifically, non-local intra-protein interactions in IDPs may regulate the conformational preferences and dynamics of both the MoRE and of the entire protein, either via direct competition or indirectly by shifting the population of free-energy minima.

## Methods

### Protein expression, purification, and sample preparation

Wild type (WT) N_TAIL_ and the seven variants listed in Supporting Table S1, were expressed in *E. coli*, and further purified as described in detail in Supporting Methods S1. Briefly, after expression of tagged protein in *E. coli*, cell pellets were resuspended in 8 M urea, 50 mM Tris pH8, 0.3 M NaCl and frozen. After thawing, the solution was sonicated, spun, and tagged proteins were purified using Ni-Sepharose fast flow beads (Cytiva). After dialyzing the eluent in 50 mM Tris pH8, 0.3 M NaCl, TEV protease was used to remove tags from the desired protein. TEV protease as well as uncut tagged proteins were eventually removed using Ni-sepharose beads. Purified N_TAIL_ proteins, free of histidine tags, were dialyzed against 15 mM Tris, 150 mM NaCl, 1 mM TCEP buffer at neutral pH, checked using SDS-PAGE, UV absorption, and far-UV circular dichroism (CD), and frozen for shipping. After shipping, samples were further purified via HPLC (semipreparative Vydac C18 column) and lyophilized. Before PET experiments, the lyophilized protein was dissolved directly into filtered buffers (for reference conditions: 15 mM Tris, 150 mM NaCl, 1 mM TCEP pH 7.6; for low pH measurements: 20 mM NaAc, 150 mM NaCl, 1 mM TCEP pH 4.0; for high salt concentrations: 15 mM Tris, 500 mM NaCl, 1mM TCEP pH 7.6). Protein concentration was adjusted to ∼100 μM as determined by UV-absorbance at 280 nm (extinction coefficient 6990 cm^-1^ M^-1^). Samples (∼350 μL) were placed in 5×10×30(h) mm Spectrocell quartz gas tight cuvettes with screwcap (equipped with Teflon coated silicon membrane), and bubbled with USP grade nitrous oxide for at least one hour, to reduce the concentration of dissolved oxygen (a quencher of excited tryptophan triplet state), and to introduce solvated electron scavenger (to prevent reactions with electrons, produced by water decomposition under UV laser pulses).

### Photo-induced electron transfer (PET) experiments

To probe intra-molecular contact formation, we used a technique based on photo-induced electron transfer (PET) between a tryptophan (W) and a cysteine (C)^34–36^ placed at different positions within the sequence (sequences in Supporting Table S1). Details are described in Supporting Methods S2. Briefly, we used a homebuilt nanosecond transient absorption apparatus to excite the W to the triplet state, and to monitor the excited state population as a function of time. When, C comes into contact with W (within Van der Waals distance) due to stochastic collisions, an excited-state electron is transferred from the triplet state of W to C. As explained in Supporting Methods S2, and in Sizemore et al. 2015^36^, the measured relaxation time is related to the intra-molecular contact formation time between W and C in the protein. The contribution to the triplet state relaxation times due to C-W quenching via electron transfer, are reported in Fig. 1D. These were obtained by fitting W triplet state relaxation curves as illustrated in Supporting Figs. S1 and S2, and taking into account the natural lifetime of the triplet state in the absence of C, measured on single W variants (see Supporting Methods S2). Each value is the result of globally fitting multiple, repeated measurements, under given solution conditions. All raw data and their fits are shown in Supporting Figs. S3 through S16. The corresponding fitting parameters, obtained for each variant under each solution condition, are reported in Supporting Table S2. Solution conditions used in PET experiments of Fig. 1 were, for reference conditions: 15 mM Tris, 150 mM NaCl, 1 mM TCEP pH 7.6; for low pH measurements: 20 mM NaAc, 150 mM NaCl, 1 mM TCEP pH 4.0; for high salt concentrations: 15 mM Tris, 500 mM NaCl, 1mM TCEP pH 7.6.

### Circular Dichroism

Circular Dichroism (CD) spectra reported in Fig. 1D inset were measured as described in Supporting Methods S3. To improve signal to noise, and access low wavelengths which are particularly important for the interpretation of IDP spectra, at each pH we carried out measurements for a series of protein concentrations and combined them. Measurements at high salt concentration were not possible due to the high absorbance of these samples below 210nm, a range which is most important for the interpretation of IDP spectra. Solution conditions used for spectra of Fig. 1 were: 10mM NaPO_4_, 150 mM, NaF for pH 7.6 measurements, and 20 mM NaAc, 150 mM NaF for pH 4.0 measurements.

### Coarse-grained molecular dynamics

We applied a coarse-grained model based on the original HPS model^43^ to N_TAIL_. Each amino acid is represented by a bead with charge (+1, 0, −1) and hydropathy. There are three types of interactions in the HPS model: bonded interactions, electrostatic interactions, short-range pairwise interactions. The electrostatic interactions are modeled using a Coulombic term with Debye-Hückel^43^ electrostatic screening to account for salt concentration. The short-range pairwise potential accounts for both protein-protein and protein-solvent interactions, which was optimized using the experimental radius of gyration of N ^6^. We further added additional terms for angle and dihedral preference so that the secondary structure of the MoRE region can be captured (see Supporting Methods S5). The HOOMD-Blue software v2.9.3^65^ together with the azplugins (https://github.com/mphowardlab/azplugins) were used for running the molecular dynamics simulations. All simulations were run using a Langevin thermostat with a friction coefficient of 0.01 ps^-1^, a time step of 10 fs and a temperature of 298 K for 5 μs. The first 0.5 μs were dumped for equilibration.

### All atom molecular dynamics

We performed all molecular dynamics (MD) simulations on three full length N_TAIL_ variants. The wild type (WT) N_TAIL_ (401-525) simulations were performed including an N-terminal hexa-histidine (His6) tag, while the latter was not included in the case of variants C3W2 (i.e. S482C-Y518W) and C4W2 (i.e. S488C-Y518W) to match the available experimental data as much as possible. All simulations were performed using the GROMACS 2019 simulation package^66^. Each N_TAIL_ variant was simulated using two different force fields (CHARMM36m^51^ and AMBER99SB-disp^49^) and at least three replicas per force field started from different initial conformations. The WT N_TAIL_ simulations were performed only with the CHARMM36m force field. For the Amber force field validation, we used previously published wild type N_TAIL_ simulation trajectories^26^. The simulations of the two variants in Amber was carried out using force field parameters adapted to the GROMACS simulation package. Both the WT trajectories and the force field parameters were kindly provided by Piana et al. Each simulation was carried out using periodic boundary conditions in a box filled with explicit solvent molecules consisting of either optimal point charge (OPC^55^, for CHARMM36m) or a modified four point-charge (TIP4P^50^ for AMBER99SB-disp) water models as well as Na+ and Cl-ions corresponding to an ion concentration of 150 mM. The total size of simulated systems was approximately 120,000 to 200,000 atoms. Total simulation time was more than 40 µs per variant with conformations recorded after every nanosecond. Hydrogen bond vibrations were constrained using virtual atom sites to enable a 4 fs time step during simulations. All simulations were kept at 1 atm pressure and 298 K temperature. Further details on the MD simulation parameters and system preparation are provided in the Supporting Methods S7 and S9. Analysis methods related to the comparison to experimental data, calculation of mutual information, contact maps, and free-energy landscapes are described in the Supporting Methods S11, S19-21.

## Data availability

The implementation of the coarse-grained HPS model can be downloaded from https://github.com/wzhenglab/ntail. Other data supporting the findings of this study are available from the corresponding authors upon reasonable request.

## Acknowledgements

L.O, J.K., G.K., S.L. and S.M.V. acknowledge the support from the National Institutes of Health (R01GM120537). A.C.V., H.G., L.V.B. and G.N. were supported by the Max Planck Society and by the German Science Foundation (DFG), Excellence Strategy Grant MBExC 2067/1; computer time was provided by the Max Planck Computing and Data Facility. G.N. was additionally supported by the Alexander von Humboldt-Foundation. A.C.V. was additionally supported by the European Union’s Horizon 2020 Framework Programme for Research and Innovation, Specific Grant Agreement No. 945539 (Human Brain Project SGA3). W.Z. acknowledges the support from the National Institutes of Health (R35GM146814) and the research computing at Arizona State University.

