## Supporting Information for "Transient Non-local Interactions Dominate the Dynamics of Measles Virus N_TAIL_"

### Supplementary Information for "Transient Non-local Interactions Dominate the Dynamics of Measles Virus $N_{TAIL}$"

#### A. Supporting Methods

**S1: $N_{TAIL}$ Protein Samples**

In addition to wild type $N_{TAIL}$, seven variants were studied in this work, namely Y451W (W1), Y518W (W2), S407C-Y451W (C1W1), S425C-Y451W (C2W1), Y451W-S482C (C3W1), S482C-Y518W (C3W2), and S488C-Y518W (C4W2) (Supporting Table S1). In these, one of the two tyrosines of wild type $N_{TAIL}$, at positions 451 and 518, was selectively substituted with tryptophan. And one of the serines, at positions 407, 425, 482 or 488 was substituted with cysteine. These mutations were chosen because they introduce minimal perturbations to the sequence, and because several of them had already been introduced and tested in previous studies. Specifically, single-site Y518W, S407C, S482C and S488C $N_{TAIL}$ variants were used in $N_{TAIL}$-$P_{XD}$ binding studies[1–4]. Proteins were expressed in *E. coli*, extracted, and further purified. In the end, all sequences studied here were free of hexahistidine tag.

**Generation of the bacterial expression constructs**

In a first step, Y451W and Y518W mutations were separately introduced by overlap extension PCR using wild-type $N_{TAIL}$ coding sequence borne by the Gateway expression vector pETG20A[5] as template and, respectively, mutagenic forward (F) and reverse (R) primer pairs Y451WF / Y451WR and Y518WF / Y518WR. After DpnI treatment, each of the two PCR products was inserted in the Gateway shuttle vector pDONR using BP clonase (Invitrogen). The sequence of the insert was checked by DNA sequencing, and inserts were transferred from pDONR to pETG20A using LR clonase (Invitrogen). In a second step, S407C, S425C, S482C and S488C mutations were individually introduced as described above using either the Y451W or the Y518W construct as template and, respectively, mutagenic primer pairs S407CF / S407CR, S425CF / S425CR, S482CF / S482CR and S488CF / S488CR. All the resulting constructs drive the expression, under the T7 promoter, of a thioredoxin (TRX) tagged fusion protein in which the $N_{TAIL}$ protein is preceded by a hexahistidine (His6) tag and a Tobacco Etch virus (TEV) protease cleavage site (TRX-His-Tev-$N_{TAIL}$). The TRX tag, the hexahistidine tag and the AttB1-encoded amino acid stretch can be cleaved off by TEV protease digestion thus leading to a protein bearing only two non-native N-terminal residues (GS).

**Protein expression and purification**

T7 cells (New England Biolabs) were transformed with the plasmid bearing the sequence coding for the fusion protein, and transformed cells were selected on agar plates containing 100µg /mL ampicillin and 34µg /mL chloramphenicol. After one night at 37°C, a few colonies were used to seed 50 ml of LB containing the same antibiotics. The preculture was incubated overnight at 37°C with constant shaking at 200 rpm. The next day, four to six liters of terrific broth (TB) containing 100µg /mL ampicillin and 34µg /mL chloramphenicol were seeded with 10mL of preculture per liter of culture and grown at 37°C with constant shaking at 200 rpm until OD at 600 nm reached 0.5 to 0.8. IPTG was added at the final concentration of 0.5 mM and expression was allowed to proceed overnight at 27°C under constant shaking. The culture was then centrifuged for 30' at 4000g. The supernatant was discarded and the cell pellet was resuspended in 10 mL of 8 M urea, 50 mM Tris pH8, 0.3 M NaCl per liter of culture, and frozen. After thawing, the cell lysate was sonicated at room temperature and spun for 30' at 15,000g at 25°C. The supernatant was supplemented with 1 mL per liter of bacterial culture of Ni-sepharose fast flow resin (Cytiva) pre-equilibrated in 8 M urea, 50 mM Tris pH8, 0.3 M NaCl and incubated for 30' on a rotating wheel at room temperature. Beads were recovered by centrifugation for 1’ at 5000g and washed five times with one volume (i.e. the same volume of beads) of 8 M urea, 50 mM Tris pH8, 0.3 M NaCl. Fusion proteins were eluted from the beads with five volumes of 8 M urea, 50 mM Tris pH8, 0.3 M NaCl, 250 mM imidazole. Pooled eluted fractions were dialyzed overnight at room temperature against five volumes of 50 mM Tris pH8, 0.3 M NaCl. The next day, the dialysis buffer was discarded and His-tagged TEV protease was added at 1/26 dilution (weight / weight, with respect to the amount of dialyzed protein) in the dialysis bag. A second dialysis was then performed against five volumes of 10 mM Tris pH8, 0.3 M NaCl at 4°C for seven hours. The content of the dialysis bag was then supplemented with 1 mL per liter of culture of pre-equilibrated Ni-sepharose resin and incubated for 30’ on a rotating wheel at 4°C. Ni beads were recovered by centrifugation for 1’ at 5000g and the supernatant containing the protein of interest was frozen. This Ni affinity step allows His-tagged proteins (i.e., uncut fusion proteins and TEV protease) to be removed, and the protein of interest to be recovered in the unbound material. After thawing, the protein was concentrated using a centricon (cutoff: 10 kDa) and the concentrated sample was kept at -20°C and shipped in dry ice.

**Further protein purification**

Initial protein purity before shipping was analyzed by electrophoresis (SDS-PAGE), absorption, and circular dichroism (CD). For storage and transportation between labs, purified proteins were dialyzed into neutral pH 15 mM Tris, 150 mM NaCl, 1 mM TCEP buffer and kept frozen in several small Eppendorf tubes. After shipping and before experiments, variants were repurified by reverse phase HPLC on a semipreparative C18 column (Vydac) and lyophilized. In all cases, protein purity and photophysical properties of W was confirmed after suspension of the lyophilized powder, by analytical HPLC, ESI-QTOF mass spectrometry (Agilent 6530 Quadrupole TOF LC-MS) and absorption spectroscopy (Cary 50 UV-Vis spectrophotometer from Agilent). Unless otherwise noted, all reagents and solvents used in this study were analytical grade obtained from Sigma. Deionized (DI) water was Millipore grade, from Evoqua Water Technologies filtration system. Lab-made buffers were pH adjusted with acid titration (Accumet AB 15, Fisher Scientific). To prevent dimerization due to C-C disulfide bridge formation, all samples contained 1 mM TCEP (freshly added to samples before measurements, in a way to ensure final pH values reported in manuscript).

**S2: Photo-induced Electron Transfer (PET) experiments**

**Measuring tryptophan-cysteine quenching times due to PET**

To quantify the intra-protein dynamics of full-length $N_{TAIL}$ in solution, we probed intra-molecular contact formation, using a technique based on photo-induced electron transfer (PET) between a tryptophan (W) and cysteine (C)[6–9]. For these experiments, we used the $N_{TAIL}$ variants of Supporting Table S1, produced as described above, containing a unique W and C placed at different positions within the sequence. A nanosecond UV excitation is used to excite W to its triplet state, and the population of this state is monitored as a function of time through transient absorption[8]. In the absence of quenchers, the W triplet state is long-lived and eventually decays to the ground state via its normal relaxation routes. In the presence of C, the W triplet state relaxes via an additional route, due to the transfer of an excited state electron from W to C. This occurs when the two amino acids come within Van der Waals distance of each other, via stochastic collisions, as the disordered protein samples its conformational states in solution. The observed relaxation rate (the inverse of the triplet state lifetime) is the sum of the natural relaxation rate in the given solution conditions, $k_o$, and the C-W quenching rate due to contact formation, $k_{CW}$:

$$k_{obs} = k_o + k_{CW}$$

At the protein concentrations used in this study, intra-molecular collisions between W and C placed on the same protein, are much faster (100ns-us time scales) than inter-molecular ones (between W and C placed on different proteins). The C-W quenching rate $k_{CW}$ therefore reflects the rate at which those two positions in the IDP contact each other[8]). We note that for simple and short IDP model peptides near physiological conditions $k_o \ll k_{CW}$ (as in our previous works) and the observed relaxation rate can be approximated by $k_{CW}$. For larger and more complex IDPs, such as $N_{TAIL}$, this is not necessarily the case. On one hand $k_{CW}$ may be significantly smaller (slower) in larger and more complex IDPs, due to larger C-W sequence separations, and to intrinsically slower intra-chain dynamics caused by interactions; on the other, the relaxation rates $k_o$ may be larger (faster), depending on pH, temperature, additional quenchers, and the position of W in the sequence. Therefore, $k_o$ should be determined on variant containing W and no C, under the given solution conditions. Throughout this work, C-W quenching times (plotted in main text Fig 1D) were obtained as:

$$\tau_{CW}^{-1} = \tau_{obs}^{-1} - \tau_{o}^{-1}$$

**PET sample preparation**

Samples for photo-induced electron transfer (PET) measurements were prepared by dissolving the lyophilized protein directly into filtered buffers (for reference conditions: 15 mM Tris, 150 mM NaCl, 1 mM TCEP pH 7.6; for low pH measurements: 20 mM NaAc, 150 mM NaCl, 1 mM TCEP pH 4.0; for high salt concentrations[1]: 15 mM Tris, 500 mM NaCl, 1mM TCEP pH 7.6). Protein concentration was adjusted to ~100 uM as determined by absorbance at 280 nm (extinction coefficient 6990 $cm^{-1}$ $M^{-1}$). Samples (~350 uL) were placed in 5x10x30(h) mm Spectrocell quartz gas tight cuvettes with screwcap (equipped with Teflon coated silicon membrane), and bubbled with USP grade nitrous oxide for at least one hour, to reduce the concentration of dissolved oxygen (a quencher of excited tryptophan triplet state), and to introduce solvated electron scavenger (to prevent reactions with electrons, produced by water decomposition under UV laser pulses).

**PET experimental setup**

We measured the lifetime of the excited W triplet state after UV pulse excitation, using a homebuilt nanosecond transient absorption apparatus. Our apparatus uses two coherent light sources, a 10 Hertz pulsed 289 nm excitation beam, and a 457.9 nm continuous probe beam, arranged in a pump–probe optical setup, similar to that in a previous work[8]. The 3 mW excitation pulse train is provided by the fourth harmonic of a Surelight Nd:YAG laser (266nm), shifted to 289 nm using a deuterium gas Raman Converter (Light Age). The excitation, Stokes, and anti-Stokes components from the Raman Converter are separated by a Pellin-Broca prism. The 289nm component is selected via a spatial pinhole filter, reduced to ~ 3.5 mm diameter, and eventually blocked by an aperture after passing through the 1 cm long and 5 mm wide sample cell equipped with a magnetic stirrer. The 200 mW, 457.9 nm Argon Ion laser beam (Coherent Innova 90C) is split into two beams. One beam, used as the probe light source, is directed to pass into the sample cell collinearity (with small angle offset) with the excitation light beam, and the other is used as the reference light source. Optical probe and reference signals are each filtered using interference filters and focused onto two separate diode detectors (New Focus, Model 2032). The outputs from each detector are directly recorded by two digital storage oscilloscopes (DPO 3032, Tektronix) to simultaneously record kinetics in both short (800 ns; with 150 MHz bandwidth, 1.25 GS/s sampling rate) and long (1 ms; with 20 MHz bandwidth, 1.00 MS/s) timescales, as to maintain high time resolution and high dynamical range. High speed shutters are employed to control the probe and excitation laser beams prior to entering the sample to protect it against unnecessary radiation exposure between measurements and to obtain background noise data.

[1] The pH was adjusted by adding amounts of acetic acid (this could be significantly different from the one estimated by weight).

With each sample, multiple measurements, each averaged over 256 laser pulses, were taken. Digital data from oscilloscopes were initially processed to reduce background noise, produce transient absorption values, and to combine the short and long timescale data using in house MATLAB (MatchWorks) scripts. Measurements, noise reduction, and initial data processing were automated using LabView. An example of the resulting raw data, monitoring the population of the W triplet state as it decays in time, is shown in S1 (all PET data presented in this manuscript can be found in Supporting Figs. S3 through S16).

**Transient absorption data analysis**

Relaxation times, $\tau_{obs}$, were obtained by fitting transient absorption data (Fig. S1) using locally written software (ASUFIT) [https://www.public.asu.edu/~laserweb/asufit/asufit.html] developed in a MATLAB environment. Multiple measurements (from multiple sample preparations as well as from repeated measurements of the same sample), taken under given solution conditions, and for a given $N_{TAIL}$ variant, were globally fit to a sum of exponentials including a rise time (Eq 1) convoluted with the Gaussian function (FWHM ~4 ns) associated with the excitation pulse time profile:

$$\Delta A(j,t) = \sum_{i=1}^{n} A_i(j) exp(-t/\tau_i) \quad (1)$$

Where ΔA(j,t) is the transient absorption signal for an individual measurement *j* as a function of time *t* and *n* is the number of kinetic components used in the fitting (n=5, including a rise time, delivered the best fit), $A_i(j)$ is the amplitude of the i-th kinetic component for the given measurement *j*, and $\tau_i$ is the global fitting parameter representing the lifetime of the i-th kinetic component, optimized over all repeated measurements j (these included multiple measurements of the same sample, as well as measurements from multiple sample preparations). An example of raw data obtained from PET measurements, highlighting the contribution to the fitting function from individual kinetic components can be found in Supporting Fig. S1**.** The kinetic component with the shortest lifetime ($\tau_1$, orange) has the largest amplitude $A_1$ (i.e., is the main component of the signal) and corresponds to the intact protein population where the excited W triplet state is quenched by electron transfer to C. The following components, $\tau_2$ (green), $\tau_3$ and $\tau_4$ (not shown) have much smaller amplitude and correspond to contributions from photochemical decomposition products (as explained in the following). Kinetic component $\tau_5$ (~20-50 ns) has negative amplitude $A_5$ and represents the initial rise in W triplet state population (formation of the lowest triplet excited state). All fitting parameters for each variant, under each solvent condition, are reported in Supporting Table S2. The corresponding raw data and fits can be seen in Supporting Fig. S3 through S16.

To understand the nature of components $\tau_2$, $\tau_3$, and $\tau_4$, we note that under intensive UV light, water decomposes and produces free solvated electrons and various chemically reactive radicals[10,11]. Electrons can reduce C, thus preventing electron transfer from W to C, and radicals can damage W and other parts of the protein. The effects of these photochemical reactions on the transient absorption signal are evident in Supporting Fig. S2. The total amplitude (the sum of all amplitudes associated with each kinetic component) decreases with each measurement (not shown). This is expected and indicates the chemical destruction of some W after long intense UV light exposure (this is more evident when small sample volumes are used, as in the present study). The amplitudes of each individual component (shown in Supporting Fig. S2) reveal which one is due to photodamage, and which is due to the actual quenching process between intact W and C. The figure shows that the largest amplitude, corresponding to the protein population where W is quenched by C ($\tau_1$ blue), decreases steadily as a function of UV laser irradiation, due to gradual depletion of intact W and C pairs. The next significant component ($\tau_2$, orange) starts at much lower amplitude and increases steadily, indicating that this decay results from a photodamage. The remaining two components ($\tau_3$, and $\tau_4$, green and red), have negligeable amplitudes and increase slightly as a function of UV irradiation, indicating that these too result from photodamage.

To avoid including measurements that were dominated by photodamage, in our global fits we only included data for which the amplitude $A_1$ of the main decay ($\tau_1$) contributed 35% or more to the total signal amplitude $\Delta A(j,0)$ (Eq (1)). This cutoff was chosen to allow enough measurements for sufficient statistics, while avoiding unwanted effects due to photodamage. All transient absorption kinetics raw data with their fits are plotted in Supporting Figs. S3 through S16. The corresponding fitting parameters are reported in Supporting Table S2. Finally, $\tau_{CW,}$ the actual contribution to the observed relaxation time due to quenching by C (as opposed to the contribution from natural relaxation), was obtained as explained above from:

$$\tau_{CW}^{-1} = \tau_{obs}^{-1} - \tau_o^{-1},$$

where $\tau_{obs}^{-1}$=$\tau_1^{-1}$ measured for the variant containing both W and C, and $\tau_o^{-1}$=$\tau_1^{-1}$ measured for the corresponding reference variant, containing only W, under given solution conditions. Values of $\tau_{CW}$ are reported in Table S2 and in the main text Fig.1D. Only values for $\tau_{CW}$ (corresponding to relaxation due to C-W electron transfer) are reported and discussed in the main text. Errors on the observed times (obtained by globally fitting multiple transient absorption signals, obtained from repeated measurements) were estimated from block averaging on mutants with the most measurements. Namely, repeated measurements were divided into groups of different bin size and globally fit within each bin. The standard error of the mean of the main decay time was calculated within each group, and plotted as a function of bin size. The plateau value yielded an error of about ±5% on the observed times obtained from global fits.

**Bimolecular quenching experiments**

To check whether the differences in C-W relaxation times $\tau_{CW}$, observed between pH 4.0 and pH 7.6 (shown in Fig. 1D main text), were due to actual conformational changes of the protein as opposed to changes in the electron transfer efficiency between W and C, we measured the bimolecular quenching rates of W and C freely diffusing in solution, at the two pH values. For this purpose, we carried out a series of PET measurements on 100 uM N-Acetyl Tryptophan Amide (NATA) solutions, containing varying cysteine concentrations, at both pH 4.0 and pH 7.6. The buffers were identical to those used for $N_{TAIL}$ in Fig. 1D (20 mM NaAc, 150 mM NaCl, pH 4.0 and 15 mM Tris, 150 mM NaCl pH 7.6). The smaller cysteine concentration range used at pH 7.6 (0 mM – 300 mM) compared to pH 4.0 (2 mM - 10 mM), was due to the lower solubility of cysteine at pH 7.6. Cysteine concentrations were determined using Ellman's reagent protocol. All sample preparation was carried out under oxygen free atmosphere in a glovebox, to avoid disulfide formation. All other details for PET experiments and data analysis were identical to those described in the above sections. The corresponding Stern-Vollmer plots of the observed relaxation rates (from the largest component of the multiexponential fit) measured at 20ºC as a function of cysteine concentration are shown in Supporting Fig. S19. The slopes of the linear fits yield a bimolecular quenching rate of 1.97 (±0.20) × $10^8$ $(M•s)^{-1}$ at pH 7.6 and 1.67 (±0.17) × $10^8$ $(M•s)^{-1}$ at pH 4.0, which are the same within experimental error (error calculated by propagating the ±5% error on $k_{obs}$ to ±10% error on the slopes).

**S3: Circular dichroism measurements**

The CD spectra of Fig. 1D inset were measured using a Jasco J-815 spectropolarimeter, in a 1mm quartz cuvette (Starna Cells), a wavelength range 190-250 nm, with a 0.4 nm step resolution, 50 nm/min speed, 2 s response time, and 1 nm bandwidth. Each sample spectrum was obtained by averaging eight spectra. Solvent conditions were 10mM $NaPO_4$, 150 mM NaF for pH 7.6 measurements, and 20 mM NaAc 150 mM NaF for pH 4.0 measurements. Protein stock solution concentration was calculated from UV-VIS absorption spectrum. To ensure high quality data at low wavelengths, where spectral shape is most significant for IDPs, we carried out measurements on samples at multiple protein concentrations, obtained by stock solution dilution (9.5 uM, 7.0 uM, 5.0 uM at pH 7.4, and 8.2 uM, 6.0 uM at pH 4.0). We used these to check measurement consistency, and signal intensity optimization at different wavelength (requiring linear concentration dependence at all wavelengths included). The data obtained then were buffer subtracted and baseline corrected, spectrally truncated according to maximum high tension (HT) voltage limitation, renormalized based on concentration, and averaged together to extend spectral range and reduce the overall signal-to-noise ratio.

**S4: Scaling of PET rates using a polymer model.**
In order to understand the scaling behavior of C-W quenching rates, $\tau_{CW}^{-1}$, measured in PET experiments, as a function of sequence separation (|*i-j*|), we utilized a polymer model, SAW-$\boldsymbol{\nu}$ model [12], in which the distance distribution function $p(r, \boldsymbol{\nu})$ is written as

$$p(r, R, \nu) = A\frac{4\pi}{R}(\frac{r}{R})^{2+g}\,exp[-\alpha(\frac{r}{R})^{\delta}\,].$$

Here $\nu$ is a varying scaling parameter, $g = (\gamma - 1)/\nu$, $\gamma \approx 1.1615$, $\delta = 1/(1 - \nu)$,[13–15] the constant $A$ and $\alpha$ are determined by normalizing the integration of $\int p(r)dr$ and $\int r^2\, p(r)dr$ to 1, and $R$ is the RMS distance. We then calculated for a polymer chain, using the SAW-$\boldsymbol{\nu}$ model, the expected C-W quenching rates for a range of sequence separation values as

$$1/\tau_{CW} = \int k_q\,(r)p(r, R, \nu)dr,$$

in which

$$k_q\,(r) = k_0 H(r - r_0),$$

a step function with $r_0$ = 0.4 nm [16]. This is the estimated reaction limited rate contributing to the relaxation times exclusively due to chain compaction/expansion, and the diffusion limited rate due to dynamics of the chain is not considered. We note that $k_0$ is not important here, since it will cancel out when investigating the scaling behavior or ratio of rates. For a given Flory scaling exponent $\nu$, the RMS distance is estimated using the scaling law

$$R_{i,j} = b|i - j|^{\nu}$$

with $b$=0.55nm [17]. By fitting the theoretically calculated values of $\tau_{CW}$ above, as a function of |*i-j*| using

$$\tau_{CW} \propto |i - j|^{\nu_{\tau}}$$

we obtain the scaling exponent $\nu_{\tau}$ for the $\tau_{WC}$, corresponding to a given Flory scaling exponent $\nu$, for $R$. As shown in Supporting Fig. S17, we find a linear relation between the distance scaling exponent $\nu$ and quenching time scaling exponent $\nu_{\tau}$. For the scaling exponent of ~0.52 estimated using the radius of gyration from SAXS experiment[18] and the SAW-$\boldsymbol{\nu}$ model[12], we obtained a corresponding scaling exponent of ~1.69 for C-W quenching time. Using this value, we fit the experimental values $\tau_{CW}$ at pH 7.6 in Fig 1D (main text), and obtain a best fit value of 0.011μs for the prefactor.

**S5: Coarse-grained models for $\mathbf{N_{TAIL}}$**
We applied the coarse-grained HPS model to $N_{TAIL}$. In the original HPS model[19], each amino acid is represented by a bead with charge (+1, 0, -1) and hydropathy. There are three types of interacions: bonded interactions, electrostatic interactions and short-range pairwise

interactions. The bonded interactions are characterized by a harmonic potential with a spring constant of 10 kJ/Å$^2$ and a bond length of 3.8 Å. The electrostatic interactions are modeled using a Coulombic term with Debye-Hückel[20] electrostatic screening to account for variation of salt concentrations. Specifically, the strength of this screening effect decays exponentially with distance, and the range is governed by the Debye length, which decreases with increasing salt concentration. The short-range pairwise potential accounts for both protein-protein and protein-solvent interactions with an adjustable parameter epsilon for the interaction strength, which was optimized using the experimental radius of gyration of IDPs and can be tuned specifically for $N_{TAIL}$ based on its radius of gyration.

Here for $N_{TAIL}$, we further added additional terms for angle and dihedral preference so that the secondary structure of the MoRE region can be captured. Specifically, for the angle potential, we used a statistical potential from a previous literature[21] for all types of amino acids. For the dihedral potential, we used a statistical dihedral potential from a previous literature[22], in which the parameters are dependent on the types of the two centering amino acids. We found that even with the statistical angle and dihedral potentials, the helical propensity of the MoRE region is still underestimated (Supporting Fig. S20). Therefore, we further introduced a harmonic dihedral potential

$$E_{harmonic-dih} = k_{dih}\,[1 + cos(\theta - 28)],$$

just to the amino acids in the MoRE region (residues 489 to 499), Here $k$ is scanned in a variety of values (Supporting Fig. S20) to also test the role of secondary structure of MoRE region to the dynamics of $N_{TAIL}$. We determined $k_{dih}$=0.2 kcal/mol so that the secondary structure of the MoRE region is close to that from NMR measurement[23]. With these angle and dihedral terms, we further scanned the interaction strength ε of the pairwise interaction term and found that ε=0.16 kcal/mol best captures the radius of gyration of $N_{TAIL}$.

The C-W quenching rate is calculated using the same step function as shown in Supporting Methods S4,

$$k_q\,(r) = k_0 H(r - r_0),$$

in which $r_0$ = 0.4 nm but now averaging over all the conformations in the simulation. We do not need an explicit $k_0$ here, since it will cancel out when investigating the scaling behavior or ratio of rates.

In order to simulate $N_{TAIL}$ at a low pH, we calculated the average charge of D and E based on their pKa values and changed the charge of these two amino acids in the CG model accordingly. For example, the pKa values of D and E are 3.65 and 4.25, respectively. At pH 4.0, the averaging charge of D and E are therefore -0.69 and -0.36, respectively.

The HOOMD-Blue software v2.9.3[24] together with the azplugins (https://github.com/mphowardlab/azplugins) were used for running the molecular dynamics simulations. All simulations were run using a Langevin thermostat with a friction coefficient of 0.01ps$^{-1}$, a time step of 10fs and a temperature of 298 K for 5 µs. The first 0.5 µs were dumped for equilibration. Further simulation conditions and lengths are provided in Table S9.

To estimate the contribution to the C-W relaxation time, exclusively due to changes in the C-W distance from the CG simulation, we first obtained the distances between the two measured amino acids (i.e. W and C) from the simulation, then we calculated the distance dependent quenching rate $k_q\,(r)$ for each conformation in the trajectory as $k_q\,(r) = k_0 H(r - r_0)$ (as described above), and at last estimated the $\tau_{WC}$ as the inverse average of $k_q\,(r)$ from all conformations (i.e. $< k_q\,(r) >^{-1}$). We note that for an all-atom model, the same distance dependent quenching rate $k_q\,(r) = k_0 H(r - r_0)$ as previously used in the polymer model can be used with a characteristic distance $r_0$ of 4 Å[16]. For a CG model, both $r_0$ and $k_0$ are unknown. The cutoff distance $r_0$ is expected to be larger than an all-atom model considering the missing atomistic interactions. We used an $r_0$ of 10Å for all the analysis. Regarding $k_0$, we always compared the ratio of the PET rates instead of an individual PET rate so determining $k_0$ for the CG model is not necessary.

**S6: All-atom Molecular Dynamics Simulations**

Molecular dynamics (MD) simulations on three Measles Nucleocapsid protein C-terminal domain ($N_{TAIL}$) variants (WT, C3W2, C4W2) were performed using the GROMACS 2019 package[25] under periodic boundary conditions, constant temperature and pressure.

**S7: General simulation setup**

The temperature of 298 K was maintained separately for solute (protein) and solvent (water and ions) atoms using a velocity rescaling algorithm[26] with a coupling constant of 0.1 ps for both atom groups. For constant pressure simulations a Parinello-Rahman barostat[27] was used to maintain a pressure of 101 kPa using a 0.1 ps coupling constant and the isothermal compressibility of water 4.5 10–5 bar−1. All simulations were propagated using a leapfrog integrator[28] in 4 fs time steps. Fast vibrational degrees of freedom were removed by using LINCS algorithm[29] using a sixth order iterative restraint on the bond angles. Apolar hydrogen positions were constrained using virtual atom sites to eliminate hydrogen bond vibrations. Electrostatic and van der Waals interactions were explicitly calculated within 1.0 nm cutoff distance, long-range electrostatic interactions beyond the cutoff were calculated by particle-mesh Ewald summation[30] with a grid spacing of 0.12 nm. Long-range van der Waals dispersion corrections[31] to the total energy of the system were applied in all simulations.

**S8: Force field parameters**

Initial simulations of WT $N_{TAIL}$ were carried out using multiple simulation parameter sets of the and Amber force field families. The tested parameter sets included two modified AMBER99 force fields, one with a rescaled TIP4P water model[32,33], and a parametrized version by Robustelli *et al*[34], AMBER03 with a scaled TIP4P water model[32], the CHARMM22* force field[35] either using a modified TIP3P water model[36] or a TIP4P water model[37], and the CHARMM36m[38] force field with either a TIP3P, TIP4P, or OPC[39] water parameters. Production MD simulations of all three $N_{TAIL}$ variants were performed using the CHARMM36m protein parameter set with OPC water parameters (C36M-OPC), and the AMBER99SB-disp (A99SB-d) parameter set including modified TIP4P water parameters optimized for intrinsically disordered protein simulations by Robustelli *et al*[34].

**S9: System preparation**

Three wild type $N_{TAIL}$ (aa. 399-525, with an N-terminal His6-tag) conformations including a helical alpha-More region were selected from a structural ensemble deposited in the protein ensemble databank[40] (PED-ID: 00020), and three further extended conformations were used to start all production runs. All conformations were solvated in a dodecahedral box of approximately 12 nm unit cell size. The box was filled with water molecules, Na+ and Cl- ions appropriate for a neutral 150 mM salt solution. All amino acids were protonated according to their dominant protonation state at pH = 7.2.

Solvent molecules around the protein were energy minimized, then initial velocities to all atoms in the system were assigned according to a Maxwell-Bolzmann distribution appropriate for 298 K temperature. The systems were then allowed to equilibrate under constant temperature and volume (*NVT*) using a Berendsen[41] thermostat for 100 ps, where the solute atomic positions were restrained by a harmonic potential and a force constant of *2500* kJ/mol/nm. In a second equilibration step, 100 ps MD simulation performed on all systems under constant temperature and pressure (*NPT*) using a velocity rescaling thermostat and a Parinello-Rahman barostat as described in the general simulation setup but maintaining the protein harmonic restraints. Finally, a 250 ps *NPT* MD simulation was performed on all systems with position restraints, and the atomic positions and velocities of the systems were used to start production MD simulations.

**S10: Force field comparison**

The behavior of intrinsically disordered proteins in all-atom MD simulations depends strongly on the choice of the parameter set used (force field)[42]. Additionally, MD force field performance for IDPs often varies in a system-dependent manner[43]. We therefore chose to validate several force fields and water models against existing independent experimental data on the wild type $N_{TAIL}$ sequence. The two best performing parameter sets were then selected to perform longer production runs of the two $N_{TAIL}$ variants used in our PET experiments. The validation data includes information on protein compactness based on small angle X-ray scattering (SAXS), secondary structure information based on circular dichroism (CD) and nuclear magnetic resonance (NMR) chemical shifts. Supporting Table S3 summarizes the computed deviations between measured observables and those predicted from MD simulations using various force fields of the AMBER and CHARMM families. For reference, the table also shows the deviations of a molecular ensemble of $N_{TAIL}$ which was obtained by fitting to NMR chemical shifts, and was deposited in the protein ensemble databank (PED).

A more detailed discussion of force field accuracy is provided below, but in summary: two all-atom force fields reproduced both $N_{TAIL}$ compactness and secondary structure propensities accurately. The most accurate simulations were obtained with the C36M force field[38] combined with optimal point charge (OPC) water[39] parameters. The second-best accuracy was achieved by the A99SB-d force field. For validation of this AMBER force field variant, we used previously published wild type $N_{TAIL}$ simulation trajectories[34], and force field parameters adapted to the GROMACS simulation package, both kindly provided by Piana et al.

We therefore selected the C36M-OPC and A99SB-d force field variants to generate a new set of simulations to sample the dynamics of two $N_{TAIL}$ variants. These variants, S488C-Y518W (C4W2) and S482C-Y518W (C3W2), displayed considerable dynamic heterogeneity in PET experiments (*i.e.* they deviated from homopolymer scaling behavior in Fig. 1D main text), and we sought to understand the structural causes of such heterogeneity through MD simulations. A summary of observables estimated from the variant simulations is shown in Supporting Table S3.

In both force fields the average radius of gyration for variants is smaller (2.5 and 2.2 nm) compared to the WT (3.0 and 2.7 nm). This is most likely due to the presence of a hexa-histidine (His6) purification tag at the N-terminus of the WT $N_{TAIL}$ sequence, which was absent in the variant sequences used in all experiments and MD simulations. Unfortunately, no SAXS measurements of the C4W2 or C3W2 sequences were available at the time of writing to confirm or disprove this hypothesis.

However, we performed CD measurements on a similar variant, Y518W (W2), without a His6 tag to estimate the expected change in secondary structure. Bayesian secondary structure estimates using the SESCA analysis package[44] indicated a significant reduction in beta-strand content (from 13±6 % to 2±6 %), even after compensating for side chain contributions of the tag. As shown in Supporting Fig. S21, the C36M-OPC simulations of both WT and variant were consistent with the measured CD spectra, whereas the A99SB-d simulations reported a significantly higher helix content (12% vs. 5±5 %) than the CD estimates.

Further, as explained in detail below, we also computed the predicted relaxation curves for the excited tryptophan triplet state population (measured in PET experiments) from simulations with both force fields, according to the method established by Zerze et al[16]. The predicted relaxation curves, shown in Supporting Fig. S22, suggest a good qualitative agreement with those measured in PET experiments. The rather large uncertainty of predicted curves is due to sparse sampling of the electron donor-acceptor (C-W) contact conformation on the timescale of the performed MD simulations.

In conclusion, our unbiased MD simulations using the C36M-OPC parameter set reproduce the experimental observables, within uncertainty, for both WT $N_{TAIL}$ and the selected $N_{TAIL}$ variants used in PET experiments. In fact, the deviations between experimental observables and those estimated from the WT $N_{TAIL}$ simulations are on par with the $N_{TAIL}$ ensemble model, deposited in the protein ensemble databank (PED.) The A99SB-d simulations also predict the wild type observables accurately but seem to overestimate the helicity of the variants. Therefore, we decided to investigate the structural reasons for the dynamic heterogeneity observed in PET experiments, based on the C36M-OPC simulations.

**S11: Comparison between all-atom molecular dynamics simulations and experimental data**

To assess the accuracy of all MD simulations, CD spectra, SAXS curves, and NMR chemical shift profiles were predicted from MD simulation trajectories conformation by conformation, then the ensemble averaged observables were compared to the measured experimental data as described below.

**S12: CD spectrum predictions and comparison**

Circular dichroism spectra were predicted from 190 to 250 nm using the SESCA package version 0.95[45]. The predicted CD spectra were computed using the DS-dTSC3 basis set, which includes three secondary structure (SS) signals (helix, beta-strand and other), and base-line corrections for side chain contributions. The SS composition $W_i$ of the protein in each conformation of simulated MD trajectories was determined by the DISICL algorithm and averaged over all trajectories of a given force field. The predicted CD spectrum is computed as

$$S_\lambda^{calc} = \sum_i W_i \cdot B_{i\lambda},$$

where $B_{i\lambda}$ is the CD signal of SS element $i$ at wavelength $\lambda$. Side chain corrections for the protein are calculated in a similar manner, based on the extracted mean CD signal of amino acid side chains ($B_{k\lambda}$), weighted by fraction of amino acid $k$ in the sequence of the protein ($W_k$).

The root means squared deviation (RMSD) of the predicted and measured CD intensities in 1000 mean residue ellipticity (kMRE) units were calculated and used as a metric to validate the MD trajectories. The global intensity of the measured CD spectrum was automatically rescaled by the factor $\alpha$ to minimize this RMSD and eliminate potential errors in the estimated experimental protein concentration (as described in a previous work[45])

$$RMSD = \sqrt{\sum_\lambda^L \frac{(S_\lambda^{calc} - \alpha \cdot S_\lambda^{exp})^2}{L}}.$$

The accuracy of MD simulations in terms of $N_{TAIL}$ SS composition was determined by comparing the RMSD between predicted and measured CD spectra to the average deviations obtained for a set of eight disordered proteins with known CD spectra and validated structural ensembles[46]. The mean *RMSD* of these model IDP-s was 2.2 kMRE with a standard deviation of 1.1 kMRE. Based on these values, the CD predictions were considered exceptionally good if they were below 1.1 kMRE (mean-sigma) and of acceptable accuracy below 3.3 kMRE (mean+sigma).

We note that four of the IDP models were taken from the protein ensemble databank (PED)[47], including a deposited model of WT $N_{TAIL}$. The mean RMSD of the IDP set is close to the 2.07 kMRE observed for globular proteins using the same parameter set, with a similar standard deviation of 1.0 kMRE.

The CD-based validation of $N_{TAIL}$ simulation ensembles relied on two CD spectra; a previously measured spectrum of WT $N_{TAIL}$ with a hexa-histidine (His6) tag attached, kindly provided by Longhi *et al*, and a more recent CD measurement of the $N_{TAIL}$ variant Y518W (without a His6 tag) measured in this work, as described in Supporting Methods S3 (pH=7.6 phosphate buffer,).

In Figure S21we compare our MD simulation ensembles of the C36M-OPC and A99SB-d force fields to the measured CD spectra. Figure S21-A shows that for the WT $N_{TAIL}$ both force fields predict the CD spectrum with similarly good accuracy (1.4 and 1.5 kMRE, respectively). In contrast, the predicted CD signal of C36M-OPC variant simulations in Supporting Fig. S21B show a much better agreement with the measured spectrum (0.9 kMRE) than those computed from A99SB-d simulations (3.3 kMRE). This difference is attributed to the differences between measured CD spectra themselves, and the significantly higher α-helix content in Amber-based simulations. To investigate further, we estimated the SS composition of $N_{TAIL}$ variants based on their measured CD spectra (as discussed in Supporting Methods S13), and compared these estimates with SS composition obtained from all-atom MD simulations.

**S13: Secondary structure predictions**

The secondary structure (SS) of $N_{TAIL}$ was estimated based on the measured CD spectra described in using a bayesian SS estimation algorithm[44] of the SESCA package version 0.95 and the CD basis set DS-dTSC3. To estimate most likely SS composition, the possible SS composition space is sampled by 50 Monte Carlo - Hastings chains started from random initial SS compositions, yielding a total 100 000 sampled SS compositions, from which the first 20 000 were discarded to exclude very unlikely initial SS. The posterior probability of sampled SS compositions $P(SS_j | CD_\lambda)$ given the measured CD spectrum is determined according to

$$P(SS_j | CD_\lambda) = P(CD_\lambda | SS_j) \cdot P(SS_j) \quad ,$$

where $P(SS_j)$ is prior probability of SS compositions *j*, taken from a uniform distribution, and $P(CD_\lambda | SS_j)$ is the likelihood of the measured CD spectrum given that SS composition. This likelihood is the joint probability of the deviation $RMSD_j$ and scaling factor $\alpha_j$ required to compare the measured CD spectrum and the predicted spectrum of the SS composition *j, as* described in Supporting Methods S12. The likelihood was estimated previously[44] from approximately 300 SS predictions of globular proteins with known CD spectra and structures.

Using the SESCA basis set DS-dTSC3, the expected SS composition of was determined from the CD spectra measured for the WT $N_{TAIL}$ with a His6 tag, as well as for the $N_{TAIL}$ variant Y518W (W2) without a tag (shown in Supporting Fig. S21). This basis set provides estimates for the protein's helix, β-strand, and random coil contents, while applying baseline corrections for amino acid side chain CD signals.

SS estimation of the measured CD spectra shows that both $N_{TAIL}$ variants remain disordered, but whereas the $N_{TAIL}$ Y518W (W2) variant shows only a small amount periodic SS (1±3 % helix and 2±6% β-strand) content, the WT variant likely has a higher propensity for β-strands (14±6%). The difference in estimated β-strand content suggests that the N-terminal His6 tag may have significant impact on the overall $N_{TAIL}$ structure, and the experimental data available for the WT $N_{TAIL}$ should not be used to validate our variant simulations. The estimated SS composition also defines a maximum helical propensity of the MORE region, which is approximately 58% for the WT $N_{TAIL}$, and 32% for the variant.

The estimated SS compositions were also compared to the SS compositions observed in MD simulations for both the WT and variant $N_{TAIL}$ variants (in Supporting Tables S3 and S4). The comparison reveals that the unexpectedly large deviation between the measured CD spectrum and the predicted spectrum of the $N_{TAIL}$ variants based on Amber99SB-disp simulations is due to an over-estimation of α-helix content (11% vs. 3±4%).

**S14: SAXS predictions and comparison**

Small angle X-ray scattering between 0.01 and 0.05 1/Å was calculated using the program CRYSOL (from ATSAS version 2.7.2.5)[48]. SAXS curves from each conformation of simulation ensembles were averaged and compared to the measured SAXS curve using a modified ² value, calculated as

$$\chi^2 = \frac{1}{N} \sum_q \left( \frac{I_q^{exp} - \alpha \cdot I_q^{calc}}{\sigma_q^{exp}} \right)^2 \quad .$$

Here $I_q^{exp}$ and $I_q^{calc}$ are the measured and calculated SAXS intensities, and $\sigma_q^{exp}$ is the experimental uncertainty of the intensity at scattering vector $q$, respectively. The scaling factor $\alpha$ rescales the predicted SAXS curve to match the experimental intensity, which is expressed in arbitrary units and depends on the experimental setup. To this aim, the $\mathcal{X}^2$ deviation is minimized as a function of α , and this minimized value quantifies the deviation between the predicted and measured SAXS curves. Because $\mathcal{X}$ describes the average deviation in relation to the experimental uncertainty, simulation ensembles were considered of good accuracy if $\mathcal{X}$ is below 1.0 and still acceptable when $\mathcal{X}$ is between 1.0 and 2.0. The SAXS-based evaluation of simulation ensembles relies on previously measured SAXS curve of the WT $N_{TAIL}$ provided by Longhi *et al*[18].

**S15: NMR chemical shift predictions and comparison**

$C\alpha$ chemical shifts for each residue in the protein was calculated for each conformation using the SPARTA+ software (version 2.6)[49]. The calculated primary chemical shifts were averaged for each $C\alpha$ atom, and compared to measured chemical shifts using the equation

$$RMSD = \sqrt{\sum_j^N \frac{(S_j^{calc} - S_j^{exp})^2}{N}}.$$

Here, $S_j^{calc}$ and $S_j^{exp}$ are the predicted and measured values of chemical shift $j$, respectively. To analyze the data, the predicted primary chemical shifts were also converted to secondary chemical shifts. To this aim, the difference of primary and secondary chemical shifts of the predicted data were subtracted from the measured primary chemical shifts. This procedure applies the same chemical shift corrections for random coil values for both measured and computed shifts.

For the $C\alpha$ chemical shift-based evaluation of $N_{TAIL}$ simulation ensembles, chemical shifts provided by Blackledge *et al* for the PED entry. This entry was later updated into PED entry 00020. The chemical shifts are identical to those published by Gely et al.[50] The agreement with these $C\alpha$ chemical shifts was considered good if the RMSD between the measured and predicted chemical shifts is smaller than the typical error of the chemical shift prediction for IDP ensembles 0.48 ppm with a standard deviation of 0.27. [46] Based on these results we considered exceptional accuracy for $C\alpha$ chemical shift predictions and below 0.21 ppm of acceptable accuracy below 0.75 ppm. Similarly, we set the accuracy for $C\beta$ chemical shifts based on the typical error of 0.52 ppm and SD of 0.1 ppm, with exceptionally good predictions below an RMSD of 0.42 ppm and acceptable accuracy below 0.62 ppm.

**S16: Triplet-state survival predictions and comparison to PET experiments**

Relaxation curves for the excited tryptophan triplet-state population, due to electron transfer between W and C, were calculated from MD trajectories according to Zerze et al[16]. The C-W relaxation times due to electron transfer $\tau_{WC}$, can be estimated from the survival probability curve $S(t)$ of the tryptophan triplet state.

The tryptophan triplet-state survival curves are calculated from the minimum donor-acceptor distances $r_{CW}$, which in turn are computed from atomic coordinates at every timestep of the MD simulation as $r_{CW} = \min(|\underline{R}_{C\gamma} - \underline{R}_{Wi}|)$. Where $\underline{R}_{Wi}$ are the positions of tryptophan ring atoms (electron donor) and $\underline{R}_{C\gamma}$ is the $S_\gamma$ atom position of the cysteine (acceptor). Note that in order to increase sampling efficacy, the donor-acceptor distances were also estimated using the tryptophan ring atom positions and the $O_\gamma$ atom position of serines in alternative acceptor positions. This allowed us to determine both C3W2 and C4W2 distances for all variant simulation trajectories, regardless of which serine-to-cysteine mutation was actually applied to $N_{TAIL}$ in the trajectory.

The survival probability curve of the triplet-state tryptophan, is calculated as

$$S(t) \; = < e^{-\int_{t0}^{t0+t} q_c \cdot H[\, r_{CW}(t\prime) - r_c \,]\, dt\prime} >_{t0},$$

where the donor-acceptor distance $r_{CW}$, $q_c = 8 \cdot 10^{-8}\, s^{-1}$, and $r_c =$ 0.4 nm are the parameters that determine the probability of quenching upon contact, and $H$ denotes a Heaviside step function that describes the number of contact conformations encountered within a time lag $t$ after the initial conformation at time $t_0$. A conformation is considered to be in donor-acceptor contact if the $r_{CW}$ distance is smaller than the contact cutoff distance $r_c$. To compute the predicted survival probability curves of $N_{TAIL}$, this calculation was averaged for every $t_0$ timestep where the integral until $t_0 + t$ can be fully evaluated within any one of the $N = 6$ simulation trajectories performed using the same force field parameters. A bootstrapping algorithm was used to assess the uncertainties of $S(t)$ by drawing 1000 sets of $N$ trajectories (with replacements) and computing the predicted survival curves. The expected value and uncertainty of $S(t)$ at each $t$ was estimated by computing the mean and standard deviation of the predicted values from each set drawn. This approach allowed us to obtain $S(t)$ curves up to 2.5 µs shown in Supporting Fig. S22 for both the C3W2 and C4W2 $N_{TAIL}$ variants. These curves were compared directly to the tryptophan triplet state relaxation curves measured in PET experiments (Supporting Methods S2) and were used to estimate the timescales of the dynamic processes in MD simulations that would influence measured transient absorptions due to C-W quenching.

For a direct comparison with the relaxation curves computed from the MD simulations, the primary data (transient absorption $\Delta A$ vs. $t$) measured in PET experiments, were transformed into survival probability curves ($S$ vs. $t$). To this aim, the initial data points before $t_{ex} = 360$ ns of the measurements were discarded. During this period, tryptophan molecules in the sample are excited, and ΔA undergoes a rapid increase to a plateau. The remaining ΔA values are transformed into survival probabilities by the equation $S(t'') \; = \; \Delta A(t) \,/\, \Delta A_{ex}$, where $\Delta A_{ex}$ is the average $\Delta A$ of the plateau, and $t'' \; = \; t \; - t_{ex}$ is the adjusted time after reaching maximum excitation. This transformation is necessary, because the initial excitation depends on the experimental setup, and MD simulations can only provide information on the relaxation decay after maximum excitation is reached. Finally, we note that (as explained in Supporting Methods S2, and further discussed below), the W triplet-state relaxation curves measured in PET experiments are the result of two types of relaxation processes: one due to C-W quenching from electron transfer to C (computed in MD), and one due to all other processes occurring naturally in solution in the absence C (not computed in MD).

**S17: Analysis of the predicted relaxation curves**

All MD-based survival curves shown in Fig. S22 panels A and B indicate several dynamic processes, occurring on multiple timescales, associated with C-W quenching upon contact. These processes can be explained using the free-energy landscape shown in Fig. 3 (main text). The first of these processes for both variants (in both force fields) are fast processes on the 10 ns timescale that affect only a small fraction of the excited $N_{TAIL}$ population. These processes are most likely associated with side chain rearrangements within the contact conformational states (S4 for C3W2 and S3 for C4W2, respectively), which bring the electron donor and acceptor in and out of contact without major structural changes of $N_{TAIL}$. Unfortunately, these processes are too fast to be captured by PET experiments, because they would occur during the triplet-state excitation process by the laser pulse.

The second fastest process is likely associated with crossing the first free-energy barrier between the free-energy minimum in the S1 conformational state and the respective contact state (S3 or S4). Time scales for these processes were determined by fitting the initial part of the survival probability curve (shown as process 2 on Supporting Fig. S22) with a single exponential function $S(t) = e^{-t/\tau_{CW}}$. We applied a Bayesian fitting method that takes into account the uncertainty of $S(t)$ to provide a mean and uncertainty estimate of the average quenching time $\tau_{CW}$.

In addition, predicted triplet-state survival curves of both force fields indicate one or two slower dynamic processes for both $N_{TAIL}$ variants, likely associated with the transitions from S2 to the contact state and the diffusion of an extended $N_{TAIL}$ C-terminus to a more compact conformation (S1 or S2). Estimating precise time scales of these processes from the MD simulations is difficult due to the sparse sampling of these slow events, and due to the expected time scales being comparable or longer than the typical trajectory length (10-20 μs) of our MD simulations.

The obtained estimated C-W quenching times from the first barrier crossing are shown in Supporting Table S4, where they are compared with the C-W quenching times ($\tau_{CW}$) of the measured PET relaxation processes. $N_{TAIL}$ MD simulations using the CHARMM36m/OPC force field predict $\tau_{CW}$ values of 5.3 (sd: 3.3) and 2.0 (sd: 0.5) μs for the C3W2 and C4W2 variants, respectively. These agree remarkably well with the $\tau_{CW}$ estimates of 7.6 and 2.8 μs from the measured PET relaxations. In comparison, crossing the first free-energy barrier appears to be a slower process using the Amber99SB-disp force field simulations, for which $\tau_{CW}$ values of 13.3 (sd: 3.2) and 4.7 (sd: 1.0) μs were computed. We note that even though the CHARMM36M/OPC simulations agree better with the measured PET relaxation times the two force fields estimate similar 2.7 (CHARMM36M/OPC) and 2.8 (Amber99SB-disp) ratios between C3W2 and C4W2 relaxation times, in very good agreement with the measured 2.8 ratio.

In Supporting Figs. S22C and S22D, we compare the predicted W triplet-state survival curves (dashed lines) with the measured PET relaxation (dots). To this aim, we corrected the predicted decay curves to account for the measured natural relaxation $\tau_0 = 39.4$ μs of the $N_{TAIL}$ variant Y518W (W2), which does not have a cysteine quencher. Assuming that the natural and quenched decays are independent of each other, the corrected survival probability is calculated as $S(t)_{corr} = S(t)_{CW} \cdot S(t)_0$, where $S(t)_0 = e^{-t/\tau_0}$. For the simulations using the CHARMM36m/OPC force field, the corrected survival curve for the C3W2 $N_{TAIL}$ variant shows an excellent agreement with the measured data (RMSD: 0.007), whereas for the fast C4W2 variant triplet-state relaxation appears to be slightly over-estimated, although this deviation (RMSD: 0.03) does not exceed the computational uncertainty (shaded areas). Natural relaxation corrections also improve the agreement between PET experiments and simulations from the Amber99SB-disp force field, which appear to underestimate the measured triplet-state relaxation for both C3W2 (RMSD: 0.04) and C4W2 (RMSD: 0.05), but these deviations are only slightly larger than the considerable computational uncertainty.

As an additional probe of how well the simulations agreed with experimental data, we also computed the relaxation times for the C1W1, C2W1, and C3W1 variants from Charm36m/OPC force field trajectories. These simulations had the wild-type tyrosine (Y) residue in position 453. This of course limits the accuracy of PET relaxation time calculations, because the Y side chain is smaller and shaped differently than a W side chain, which in turn alters the short-range dye-quencher distance distribution. To partially compensate for the differences, the cutoff-distance for a reactive contact $r_c$ was increased 0.6 nm for these calculations. With this modification, triplet-state relaxation curves and times were estimated as described above, and the relaxation times of all five variants were plotted as the function of sequence separation in Fig S30, similarly as was done for Fig 1D. The plot indicates that within the significant computational uncertainty, three out five relaxation times of the $N_{TAIL}$ PET variants are reproduced. The relaxation time for the C4W2 variant is slightly underestimated (2.0 ± 0.5 μs vs. 2.7 μs), whereas the relaxation time for C3W1 is overestimated (5.7±1.8 μs vs. 3.2 μs) by the simulations. The latter deviation may suggest that non-local interactions and contact frequency between $N_{TAIL}$ regions B and D (seen state S2) are under-sampled in Charmm36m/OPC simulations.

**S18: Data Processing**

Basic plots and images in this work were generated using Grace v5.1.25 and Python environment v3.7.3, with the Numeric Python (NumPy) v1.16.4, Matplotlib v3.1.0, Scientific Python (SciPy) v 1.3.0, and Seaborn v0.9.0 extensions. Images were processed into finalized figures using GIMP v2.10.22 and Microsoft Office 2019.

**S19: Free energy landscape calculations**

The free energy surface shown in Fig. 3 of the main manuscript was computed from the minimum PET donor-acceptor distances described in Supporting Methods S16. Donor-acceptor minimum distance trajectories were converted into histograms between 0 and 100 Å, with bin width of 1 Å. Histogram counts were normalized and converted into probability densities using the Gaussian kernel density estimator from scipy v1.3.0. The free energy of bin *i* (in Boltzmann units) was determined as $F_i = -\log(P_i)$. The free energies of each bin were adjusted by an offset $F_0 = \min(F_i)$ to set the global free energy minimum to zero, and all free energy values above the upper limit $F_{max}$= 6 kT was set to $F_{max}$ to provide an easy visualization of the landscape.

**S20: Contact map calculations**

The contact maps shown in Fig. 4, were calculated from ensembles of conformations extracted from all-atom MD simulations that are within free-energy minima of Fig. 3 of the main manuscript (see highlighted areas). Backbone contact maps of $N_{TAIL}$ variants were calculated based on Cα - Cα pair distance distributions of these conformational ensembles. For each conformation, residues $i$ and $j$ were considered to be in a backbone contact if the distance between their Cα positions was smaller than the cutoff distance $D_{C\alpha}$ = 0.6 nm. Salt bridge contact maps were calculated in a similar manner, with $D_{salt}$ = 0.35 nm cutoff distance between amide hydrogen atoms of positively charged side chains donor residues (arginines and lysines) and negatively charged acceptor residues (Oγ of aspartate, Oδ of glutamate, or terminal carbonyl oxygen). Additionally, salt bridges were required to have N-H-O angle of 110°or larger.

**S21: Coevolutionary analysis**

Coevolutionary analysis was performed using EVcouplings (https://evcouplings.org/) with default parameters[51]. The statistical inference method is Pseudo-likelihood maximization. The Bitscore sequence inclusion threshold is 0.1. The sequence coverage is from residue 443 to 501

(59 amino acids). A total of 12072 sequences have been aligned and selected for analysis. Therefore, there are 204 sequences per amino acids with a run quality score of 8 (10 best, high quality). The coupling score instead of the folding model has been used to generate the contact map (Fig. S28) since for an IDP, these strongly coupled sites might not always be in contact.

**S22: Mutual information calculations**

To quantify correlations (including non-linear correlations) between the occurrence of PET donor-acceptor contact formation and other interactions, we calculated normalized pointwise mutual information (NPMI) scores based on the MD simulation trajectories. NMPI scores of Table S5 between PET donor-acceptor contacts (C3W2 or C4W2) and either backbone contacts or salt-bridge contacts (as described in Supporting Methods S20) were calculated from every conformation of all simulation trajectories of a given force field (*i.e.* not just within conformational states). PET donor-acceptor pairs were considered to be in contact if their minimum distance (see Supporting Methods S16) was below 0.4 nm. The NPMI scores were calculated according to:

$$NPMI(x,y) = -\frac{1}{\log(P(x,y))} \cdot log\left(\frac{P(x,y)}{P(x) \cdot P(y)}\right)$$

Where $P(x)$ and $P(y)$ are, respectively, the probabilities of a PET donor-acceptor contact and of intramolecular contact (either backbone or salt-bridge) observed throughout the simulations. $P(x,y)$ is the probability of the two contacts observed simultaneously.

To establish if there is a direct correlation between the presence of identified contacts and the C-W electron transfer, we selected several contacts that are prominent in at least one of the conformational states, and computed the normalized pointwise mutual information (NPMI) between them and the two C-W donor-acceptor contacts as shown in Supporting Table S5. NPMI describes the co-occurrence of two events (see Supporting Methods S21). NPMI values range on a logarithmic scale between -1 (never co-occurring) and 1 (always co-occurring), with independent occurrences resulting in an NPMI value of zero.

As Table S5 shows, all NPMI values for S1 non-local interactions are negative, indicating that the observed interactions in this conformational state rarely co-occur (i.e., are anti-correlated) with donor-acceptor contacts. This is expected because, as mentioned above in the contact map analysis, these interactions stabilize a non-contact conformational state. The negative NPMI values indicate that these interactions would indeed hinder electron transfer for both C4W2 and C3W2. There are few prominent interactions observed in the S2 state, with interaction between regions A, B, C often having positive correlations to C-W contact formation, and interactions between regions B and D showing negative correlations with C-W contact formations of both variants. Interactions observed in the S3 and S4 states have considerable positive NMPI-s with C4W2 and C3W2 donor-acceptor contact formation, respectively, confirming that these interactions co-occur and likely promote electron transfer.

Interestingly, the salt bridge R413-E480, which connects regions A and D, also has a positive NPMI with both C-W contacts, possibly stabilizing conformations that enable electron transfer for both C4W2 and C3W2. Further, the table highlights certain charged residues, like R413 from region A, K441 and R444 from region B, R490 from region D, as well as E511 and D520 from region E that apparently form different prominent stabilizing interactions in multiple conformational states. The mutual information between salt bridges occurring in different states shows strong negative correlations (NPMI of -0.25 – -0.50), indicating a set of competing interactions that may drive transitions between conformational states. The highlighted charged residues in region B apparently can form either non-local salt bridges with residues distant in sequence close to the MoRE region, or local salt bridges with other charged residues within region B. We observe that many of these salt bridges are accompanied by clusters of other interactions (see encircled clusters in Fig. 4). It is possible that the formation of these clusters is nucleated by the salt bridges. In any case, switching between these sets of interactions is necessary for the system to transition between the most populated conformational states S1 and S2 in Fig. 3, and competition between these interactions is what determines the populations of the conformational states themselves.

We also used mutual information analysis to determine if certain non-local interactions are correlated to helix formation in the MoRE region, which may be of functional importance. To this aim, we computed NPMI scores between non-local contact formation and three substates of helix folding observed in out MD simulations. The low helix substate is defined by less than 20% of the $N_{TAIL}$ MoRE residues found in a helical conformation which is the dominant helix sub-state both in the S1 and S4 states. The medium helix substate is intermediate folding state, where 20-50% of the MoRE residues are helical. This substate appears frequently in the S3 state, where the N-terminal part of the MoRE is folded more often than the C-terminal part. Finally, the high helix substate refers to conformations where 50-100% of MoRE folded, which is observed in (but not limited to) S2 conformational state.

Computing NPMI values between helix substates and interactions of region D, with particular attention paid to the interactions of residues that participate in binding the phosphoprotein X-domain ($P_{XD}$) shown in Supporting Table S7. The NPMI scores correlating selected prominent non-interactions with helix substates (low, medium, or high) are shown in Supporting Table S6**.** Most of the interactions can placed in one of three broader categories.

The first group (red) interactions are negatively correlated with both medium and high helix contents, which includes several salt bridges from the S1 and S4 conformational states. These interactions likely stabilize non-helical conformations, and thus are expected to have an indirect and negative effect on phosphoprotein binding affinities. The second group of interactions (green) are positively correlated with both medium and high helix contents. Examples of such interactions include residues that directly form the binding site for phosphoprotein X-domain ($P_{XD)}$ such as A447-M501. Such interactions observed in MD simulations may suggest a competition for the folded MoRE binding site between $P_{XD}$ and intramolecular interactions between regions B and D. Interestingly this group also contains local salt-bridges within region B, such as K441-K449, which suggests that such interactions may allosterically promote helix formation. The third group of interactions (blue) are defined by positive correlations towards the medium helix substate, but low or negative correlations to the high and low helix substates. These interactions, such as R405-D484 observed in the S3 conformational state, or A409-T469 in S2, may stabilize folding intermediates of the MoRE region, thus reducing free-energy barriers and increasing binding/unbinding rates of $P_{XD}$.

**S23: MoRE helicity in all-atom MD simulations**

The most prominent secondary structure element in $N_{TAIL}$ is the MoRE region (residues 484-502) that according to the NMR chemical shifts determined by Gely et al[50], folds into a transient α-helical structure for up to 50% of time. This transient helix is stabilized and folds into four-helix bundle when binding to Measles phosphoprotein X domain ($P_{XD}$) which is an important part of the viral replication process of measles.

There have been previous simulation studies that focused on the helicity in the MoRE region and the coupled binding and folding of the MoRE-$P_{XD}$ complex. Robustelli *et al.*[52] used the Amber99SB-disp force field to perform a single 100 μs simulation of the MoRE region, and used two metrics to compare to NMR experiments; the RMSD between measured and computed Cα chemical shifts (RMSD: 0.52 ppm) and Pearson correlation of the same data (R: 0.92). Wang *et al.*[53] performed 96 130 ns temperature replica exchange simulations using the

CHARMM22* force field for a total sampling time 12.4 μs on MoRE. They only provided Pearson correlation (0.76) for the chemical shift profiles, which appeared to be in good agreement with the measured profiles. The two studies had differences in details, but both agreed on the approximately 20% helix content in the MoRE region, considerably more helical content (~50%) at the $P_{XD}$ binding site (491-501) than the region preceding it (482-490). They both suggested a strong induced fit character of coupled binding and folding mechanism.

Our two best wild-type $N_{TAIL}$ (401-525) simulations were performed using CHARMM36m-OPC force field (RMSD: 0.47 ppm, R: 0.83) and the Amber99SB-disp (RMSD: 0.36 ppm, R: 0.84) simulation provided by Piana *et al*. The former set of simulations were performed in six replicates using different starting structures for 5-10 μs length (for a total of 41 μs), whereas the latter was a single 30 μs trajectory, which was divided into three parts and analyzed separately for determining sampling uncertainty (Table S8). Both of these simulations of the full $N_{TAIL}$ predict transient helical structures of the MoRE region, with similar 16-18% helix content. The two force fields also agree in that the $P_{XD}$ binding site (491-501) has a higher content than the preceding segment as shown in Fig. S31, but the difference is considerably smaller for the Charmm36M-OPC simulations than for Amber99SB-disp. This difference appears to be a force-field family dependent difference, because similar distributions are shown in the helicity profiles of Wang and Robustelli as well.

As far as we can tell, no NMR data is available for the PET variants, but the measured CD spectra of W2 variant indicate a change in the secondary structure composition compared to the wild type construct (likely with a small decrease of the helix content). This difference may be caused by a combination of the different ionic strength, the absence of the N-terminal hexa-histidine tag, and the Y518W point mutation.

Simulation trajectories C3W2 and C4W2 variants using CHARMM36m-OPC show a smaller helix content (~10%) in the MoRE, with a much lower helicity at the 482-490 MoRE segment, which is compatible with the W2 CD spectrum. In contrast Amber99SB-disp simulations predict an increased helix content (~29%) in the same region. We also note that helix folding dynamics in the CHARMM36m forcefield appears to be considerably slower than in Amber, and the average secondary structure content did not fully converge within the 20 μs simulation trajectories (showing a slow upwards trend in the second half of the simulations).

#### B. Supporting Figures

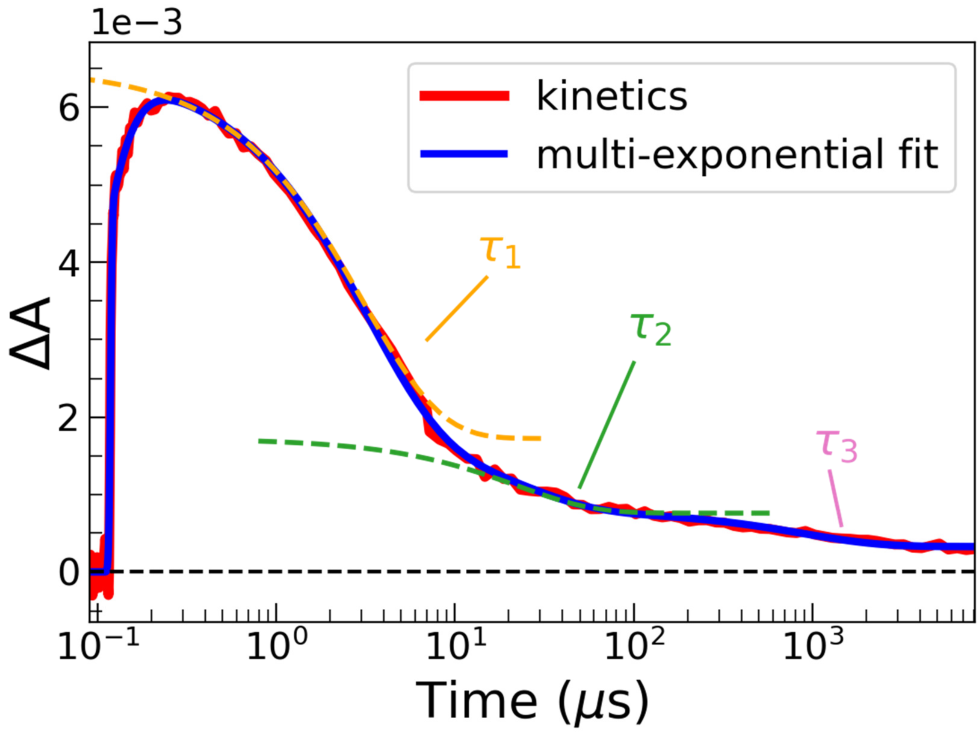

**Fig. S1.** Example of PET experiment raw data (red) with a multi-exponential fit (blue). ΔA (the transient absorption measured at 458 nm) is proportional to the W triplet state population, which decays in time. Dashed curves represent individual exponential decays contributing to the fit. The main component of the signal $\tau_1$ (orange) corresponds to the observed triplet state relaxation time from the intact protein, in the presence of the C quencher. The smaller components at longer times ($\tau_2$ , $\tau_3$,…) correspond to photoproducts, as shown in Supporting Fig. S2**.**

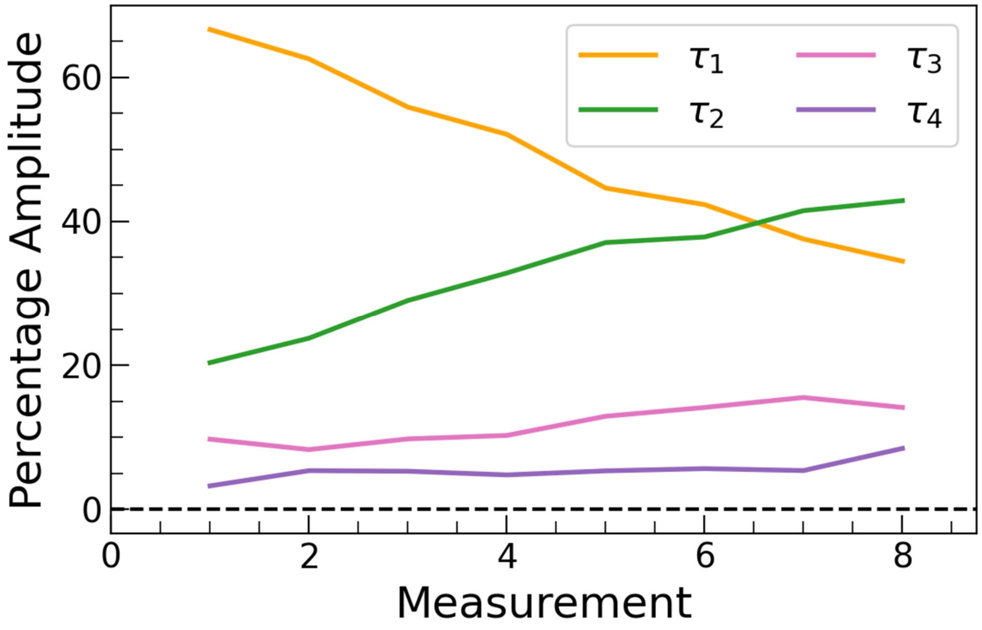

**Fig. S2.** The percentage amplitude of the multi-exponential fit components, as a function of UV pulse irradiation (each measurement corresponds to 256 UV pulses). The largest amplitude corresponds to the main component $\tau_1$ (6.37 μs, orange), and decreases steadily as a function of UV laser irradiation, indicative of photo-damage. The next significant component ($\tau_2 =$ 48.1 μs, green) starts at much lower amplitude and increases steadily, indicating that this is a photoproduct. $\tau_3$ (1020 μs, pink) and $\tau_4$ ($1 \times 10^6$ μs, purple) components follow suit and are much smaller. Sample conditions: 482C-518W in 15 mM Tris, 150 mM NaCl, pH 7.6.

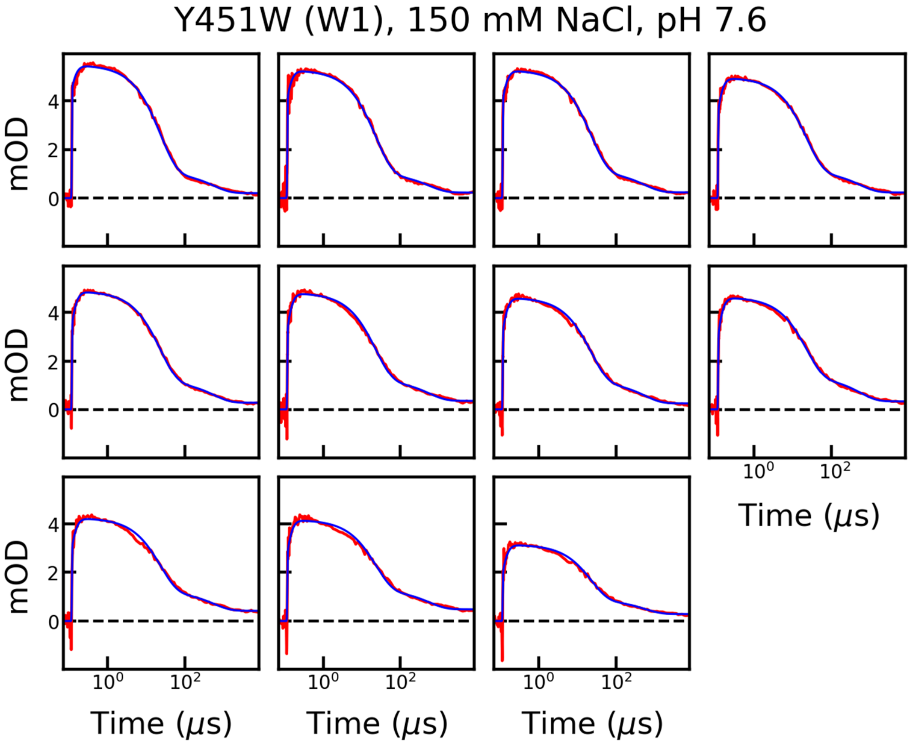

**Fig. S3.** PET experiment raw data (red) with multi-exponential fit (blue) for each measurement of Y451W (W1). Experimental conditions: 15 mM Tris, 150 mM NaCl, 1 mM TCEP, pH 7.6. To avoid excessive photodamage, only measurements for which the main component contributed 35% or more to the amplitude are used.

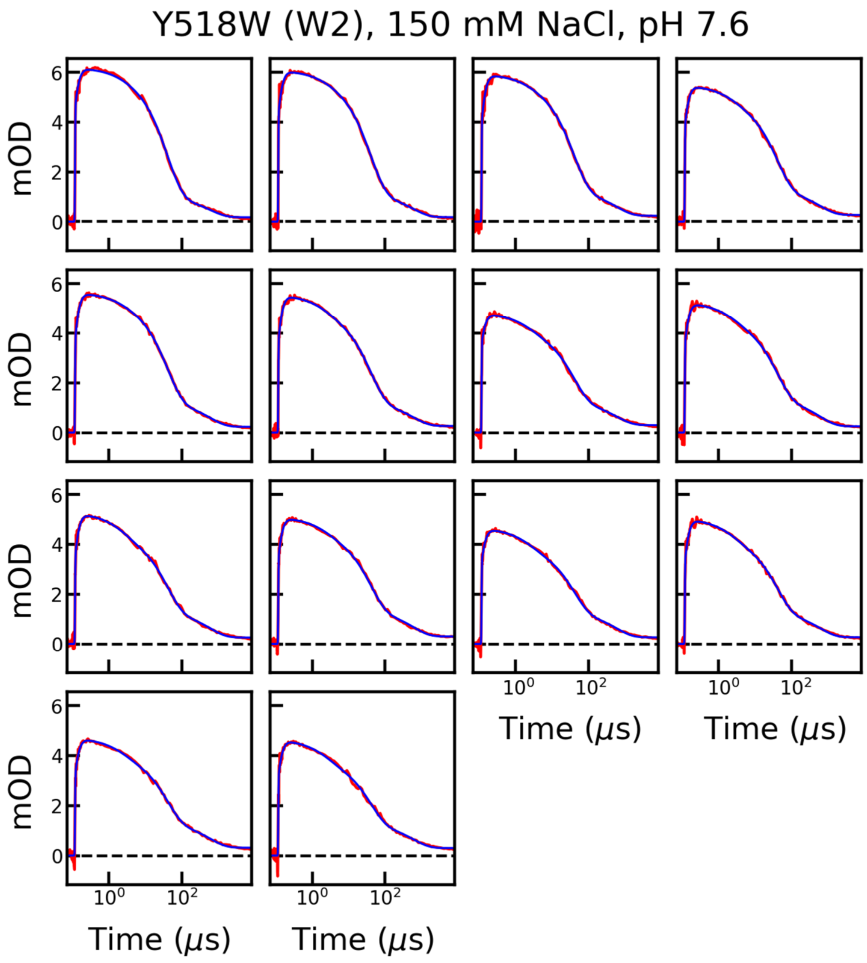

**Fig. S4.** PET experiment raw data (red) with multi-exponential fit (blue) for each measurement of Y518W (W2). Experimental conditions: 15 mM Tris, 150 mM NaCl, 1 mM TCEP, pH 7.6. To avoid excessive photodamage, only measurements for which the main component contributed 35% or more to the amplitude are used.

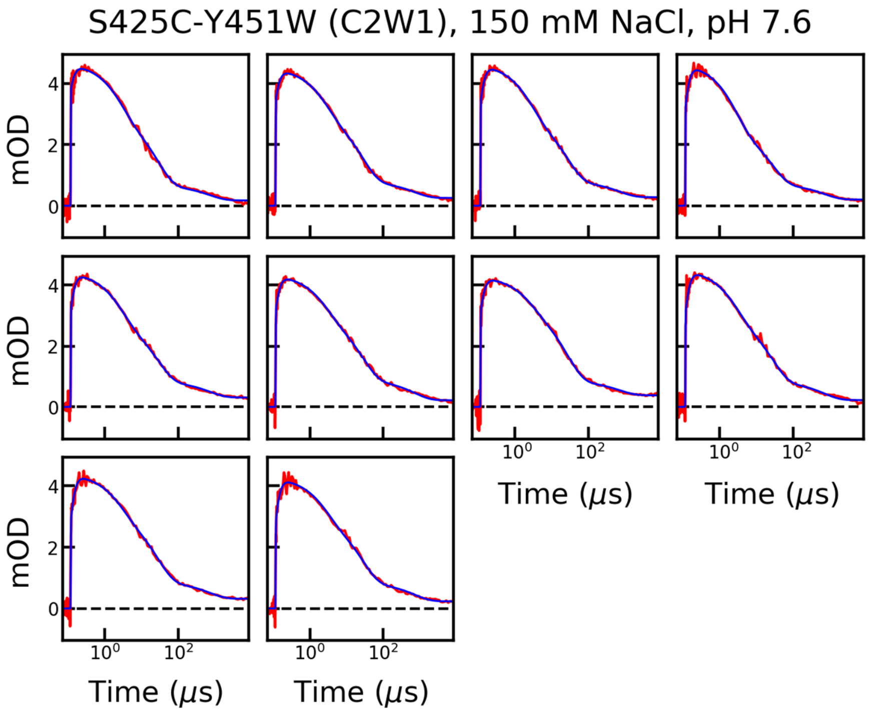

**Fig. S5.** PET experiment raw data (red) with multi-exponential fit (blue) for each measurement of S425C-Y451W (C2W1). Experimental conditions: 15 mM Tris, 150 mM NaCl, 1 mM TCEP, pH 7.6. To avoid excessive photodamage, only measurements for which the main component contributed 35% or more to the amplitude are used.

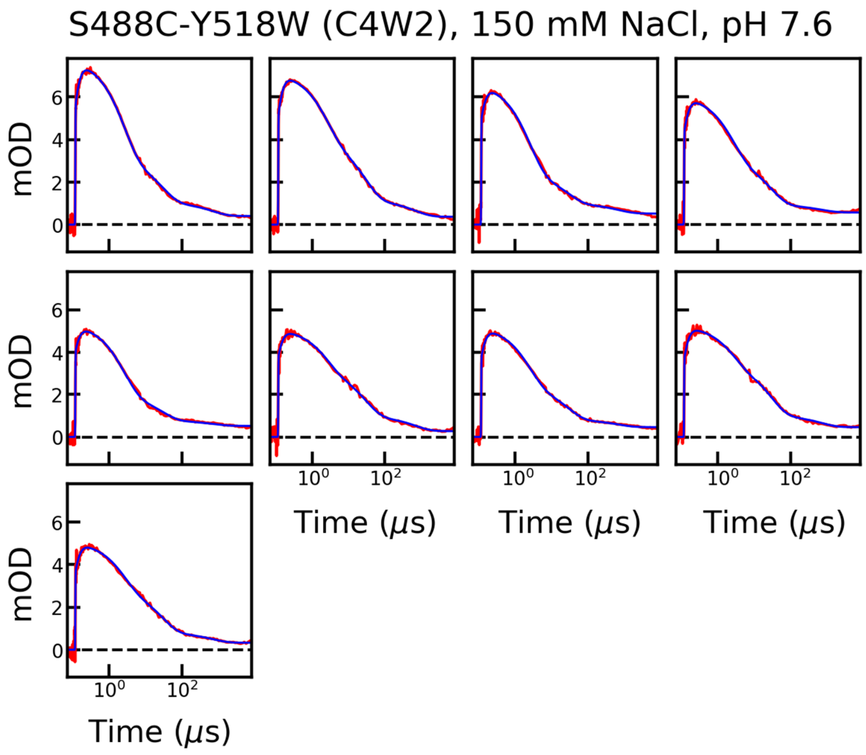

**Fig. S6.** PET experiment raw data (red) with multi-exponential fit (blue) for each measurement of S488C-Y518W (C4W2). Experimental conditions: 15 mM Tris, 150 mM NaCl, 1 mM TCEP, pH 7.6. To avoid excessive photodamage, only measurements for which the main component contributed 35% or more to the amplitude are used.

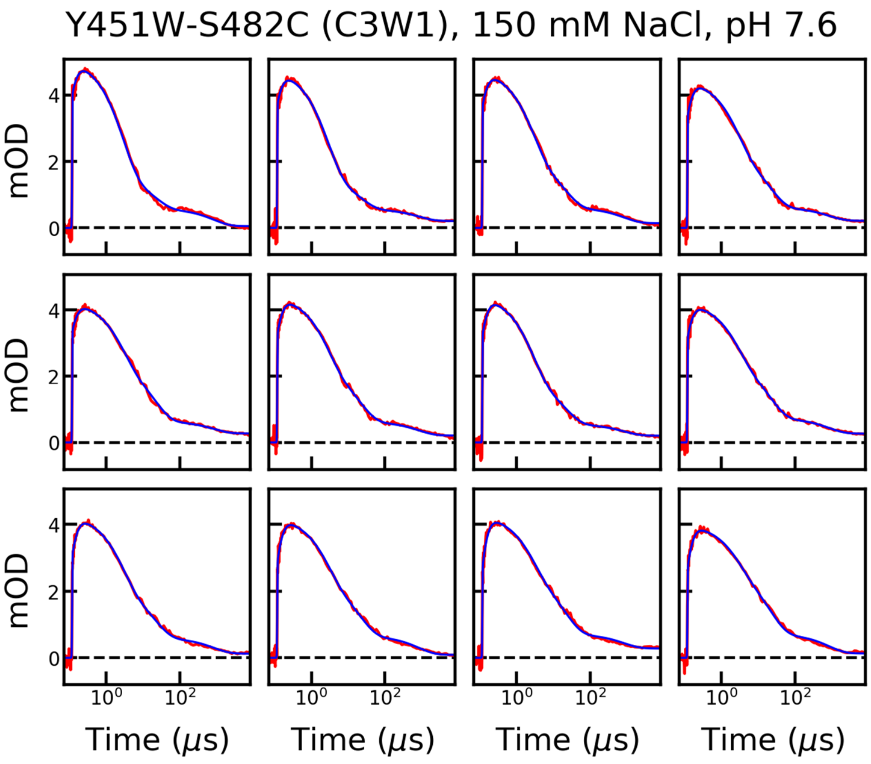

**Fig. S7.** PET experiment raw data (red) with multi-exponential fit (blue) for each measurement of Y451W-S482C (C3W1). Experimental conditions: 15 mM Tris, 150 mM NaCl, 1 mM TCEP, pH 7.6. To avoid excessive photodamage, only measurements for which the main component contributed 35% or more to the amplitude are used.

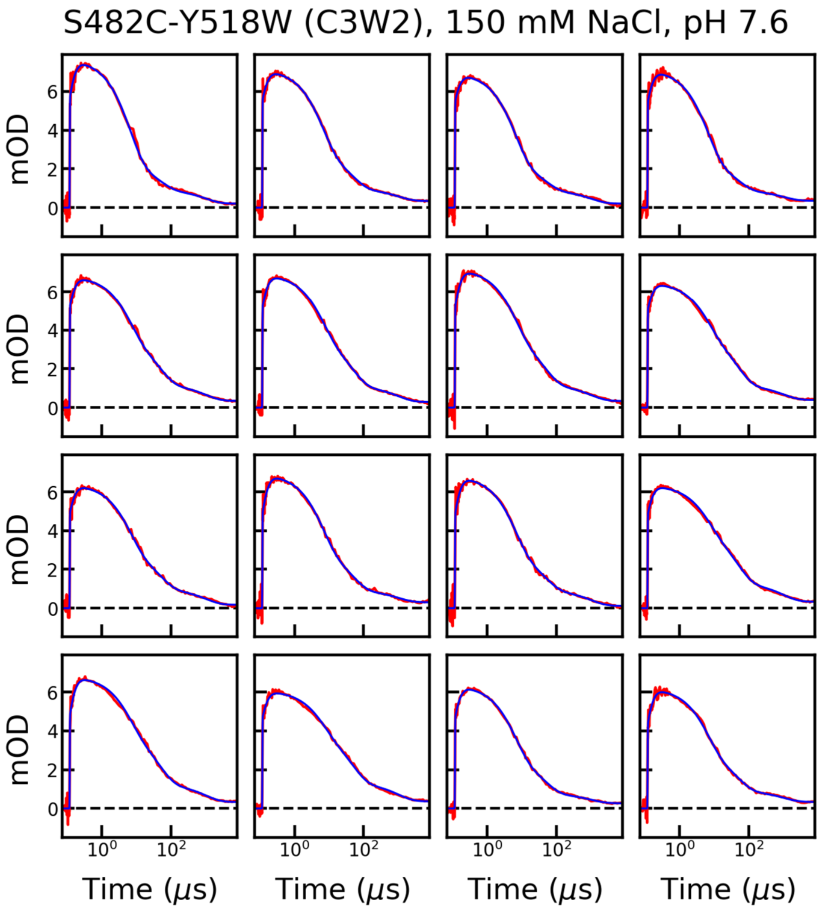

**Fig. S8.** PET experiment raw data (red) with multi-exponential fit (blue) for each measurement of S482C-Y518W (C3W2). Experimental conditions: 15 mM Tris, 150 mM NaCl, 1 mM TCEP, pH 7.6. To avoid excessive photodamage, only measurements for which the main component contributed 35% or more to the amplitude are used.

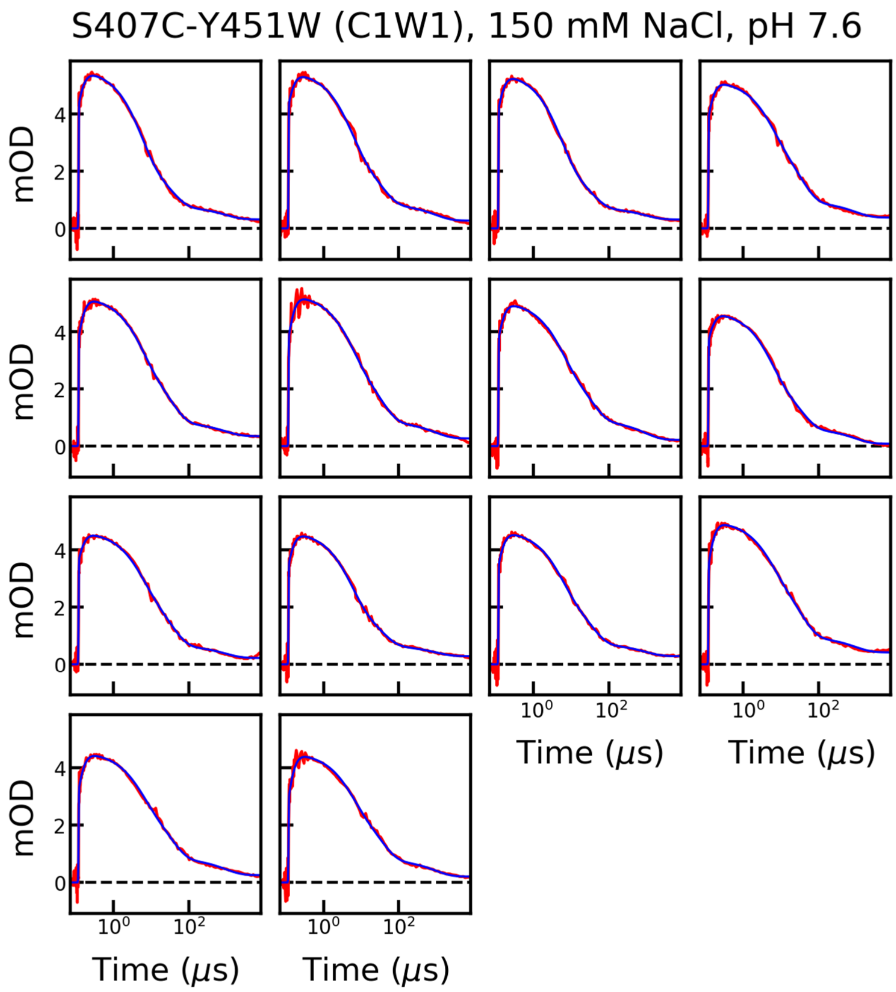

**Fig. S9.** PET experiment raw data (red) with multi-exponential fit (blue) for each measurement of S407C-Y451W (C1W1). Experimental conditions: 15 mM Tris, 150 mM NaCl, 1 mM TCEP, pH 7.6. To avoid excessive photodamage, only measurements for which the main component contributed 35% or more to the amplitude are used.

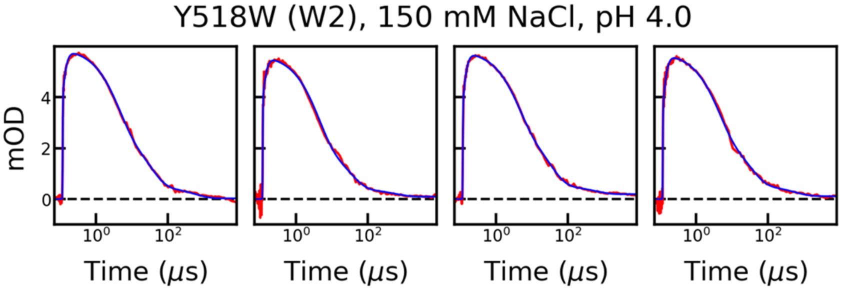

**Fig. S10.** PET experiment raw data (red) with multi-exponential fit (blue) for each measurement Y518W (W2). Experimental conditions: 20 mM NaAc, 150 mM NaCl, 1 mM TCEP, pH 4.0. To avoid excessive photodamage, only measurements for which the main component contributed 35% or more to the amplitude are used.

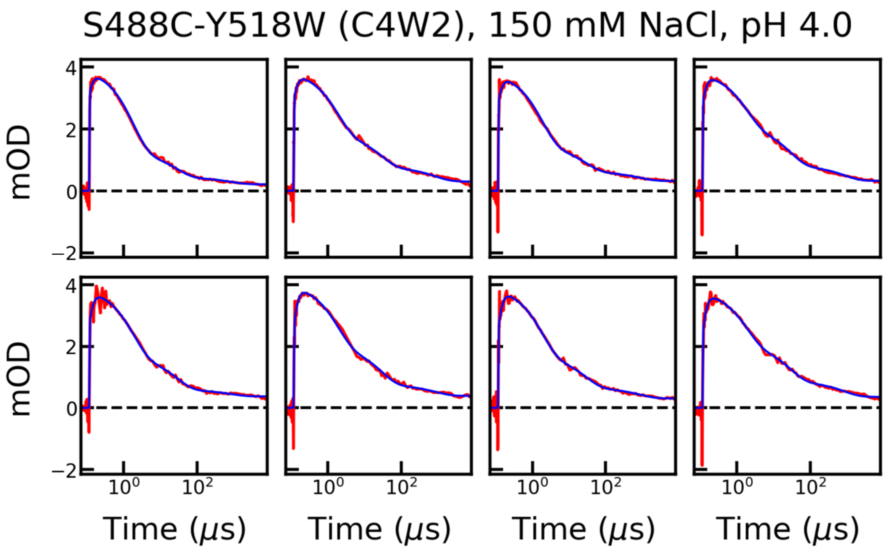

**Fig. S11.** PET experiment raw data (red) with multi-exponential fit (blue) for each measurement of S488C-Y518W (C4W2). Experimental conditions: 20 mM NaAc, 150 mM NaCl, 1 mM TCEP, pH 4.0. To avoid excessive photodamage, only measurements for which the main component contributed 35% or more to the amplitude are used.

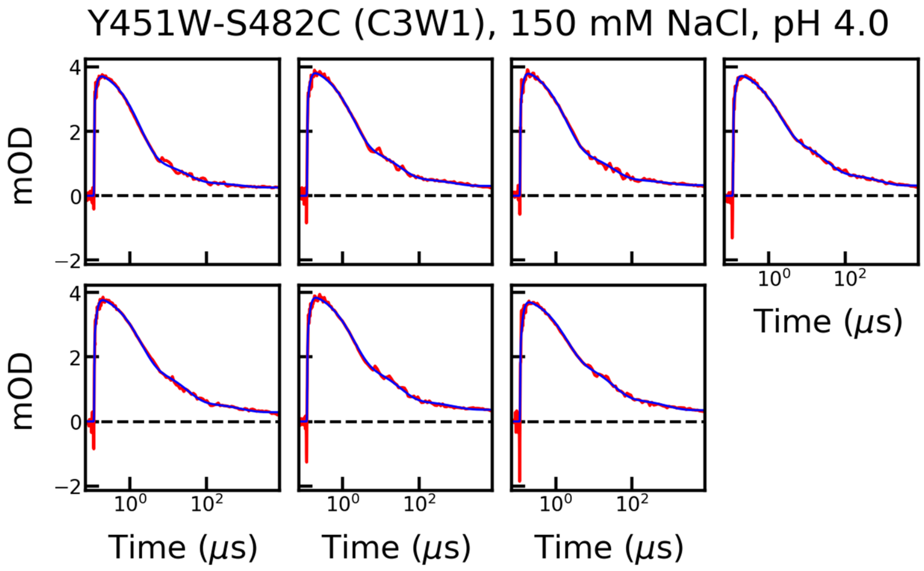

**Fig. S12.** PET experiment raw data (red) with multi-exponential fit (blue) for each measurement of Y451W-S482C (C3W1). Experimental conditions: 20 mM NaAc, 150 mM NaCl, 1 mM TCEP, pH 4.0. To avoid excessive photodamage, only measurements for which the main component contributed 35% or more to the amplitude are used.

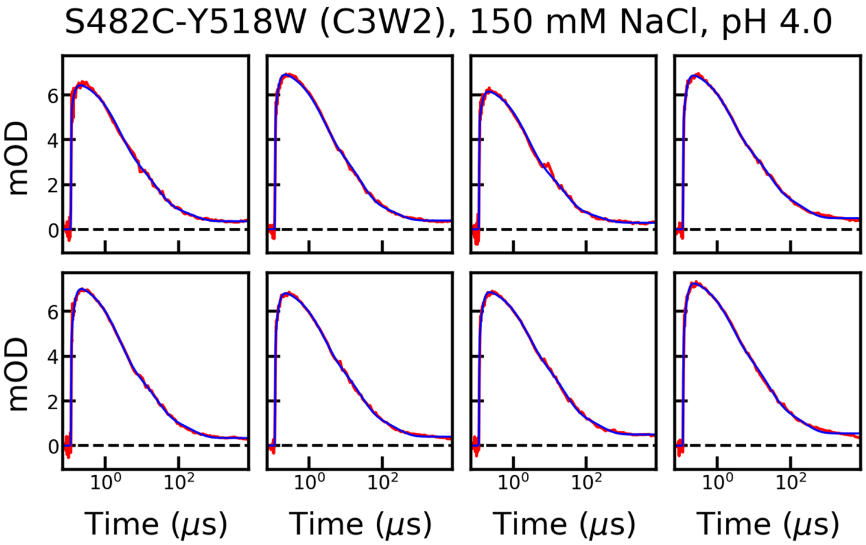

**Fig. S13.** PET experiment raw data (red) with multi-exponential fit (blue) for each measurement of S482C-Y518W (C3W2). Experimental conditions: 20 mM NaAc, 150 mM NaCl, 1 mM TCEP, pH 4.0. To avoid excessive photodamage, only measurements for which the main component contributed 35% or more to the amplitude are used.

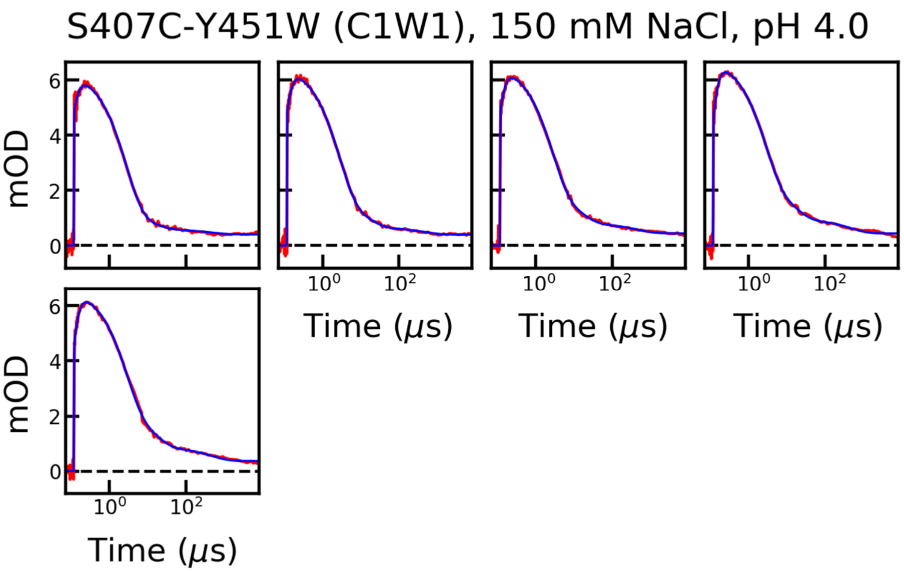

**Fig. S14.** PET experiment raw data (red) with multi-exponential fit (blue) for each measurement of S407C-Y451W (C1W1). Experimental conditions: 20 mM NaAc, 150 mM NaCl, 1 mM TCEP, pH 4.0. To avoid excessive photodamage, only measurements for which the main component contributed 35% or more to the amplitude are used.

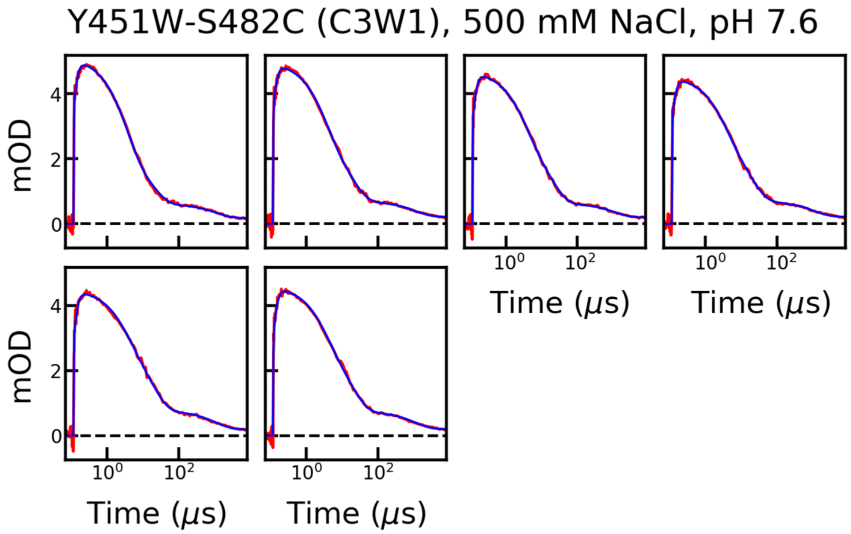

**Fig. S15.** PET experiment raw data (red) with multi-exponential fit (blue) for each measurement of Y451W-S482C (C3W1). Experimental conditions: 15 mM Tris, 500 mM NaCl, 1 mM TCEP, pH 7.6. To avoid excessive photodamage, only measurements for which the main component contributed 35% or more to the amplitude are used.

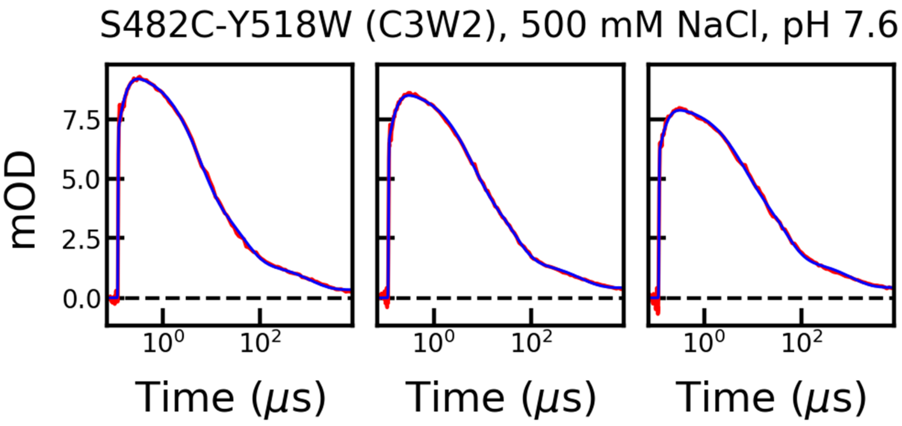

**Fig. S16.** PET experiment raw data (red) with multi-exponential fit (blue) for each measurement of S482C-Y518W (C3W2). Experimental conditions: 15 mM Tris, 500 mM NaCl, 1 mM TCEP, pH 7.6. To avoid excessive photodamage, only measurements for which the main component contributed 35% or more to the amplitude are used.

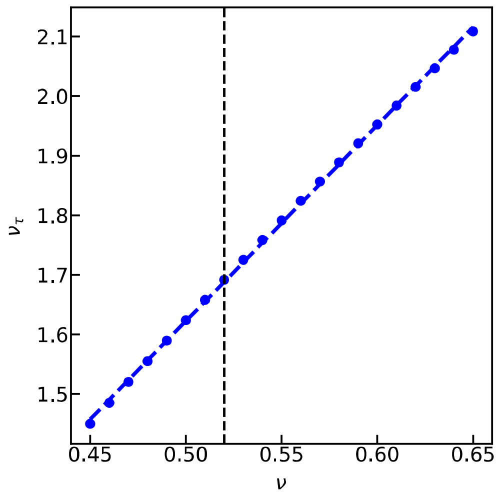

**Fig. S17.** Linear dependence between the C-W quenching time scaling exponent $\nu_\tau$ and the Flory (distance) scaling exponent $\nu$. $\nu_\tau$ was calculated for different values of $\nu$, using the SAW-$\boldsymbol{\nu}$ model (blue points). The black dashed line indicates the value of $\nu$ for Ntail at pH 7.6. The linear best fit function (blue dashed line) $\nu_\tau = 3.292 \times \nu - 0.024$ can be used to describe the relation between the two scaling exponents.

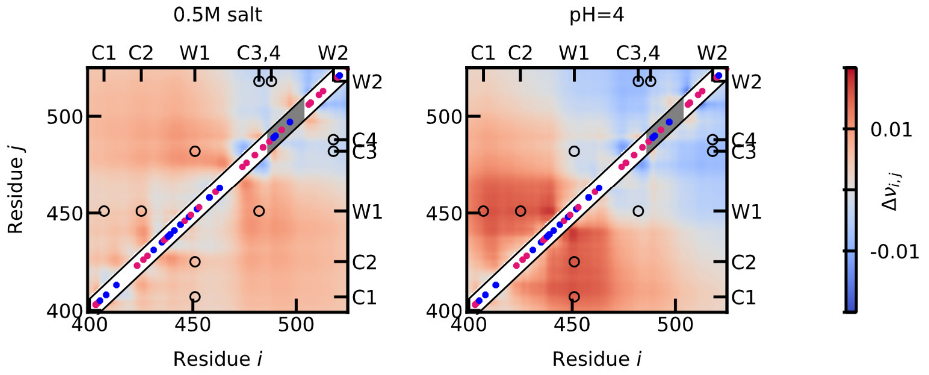

**Fig. S18.** A) The difference of the pairwise scaling exponents ($\nu_{i,j}$) between the CG simulations at 0.5M salt (Left) and pH 4.0 (Right) and the simulations at the reference condition (0.15 M salt and pH 7.6). The pairwise scaling exponents are calculated from the pairwise distances ($R_{i,j}$) as $R_{i,j} = b|i-j|^{\nu}$, with $b$=0.55nm[12,17]. The diagonal shows the charged amino acid positions with positively charged amino acids in blue and negatively charged amino acids in magenta. The MoRE is highlighted in grey.

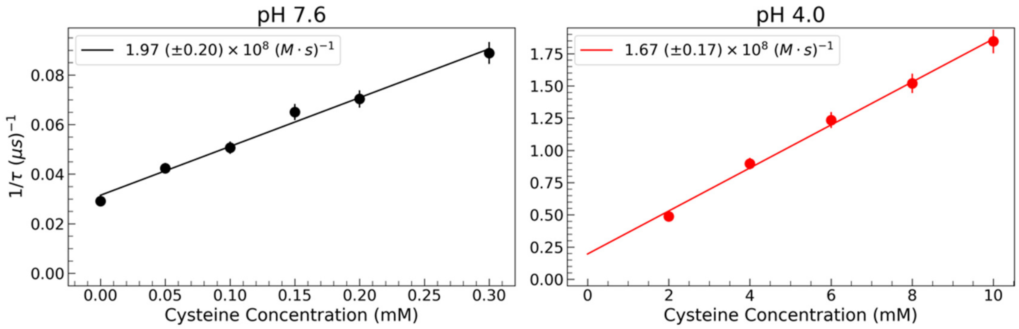

**Fig. S19.** Bimolecular quenching rates at pH 7.6 (black) compared to pH 4.0 (red). Note the different scales on both the x- and y-axes. The calculated rate at pH 7.6 is 1.97 (±0.20) × $10^8$ $(M•s)^{-1}$ compared to 1.67 (±0.17) × $10^8$ $(M•s)^{-1}$, which are the same within experimental error.

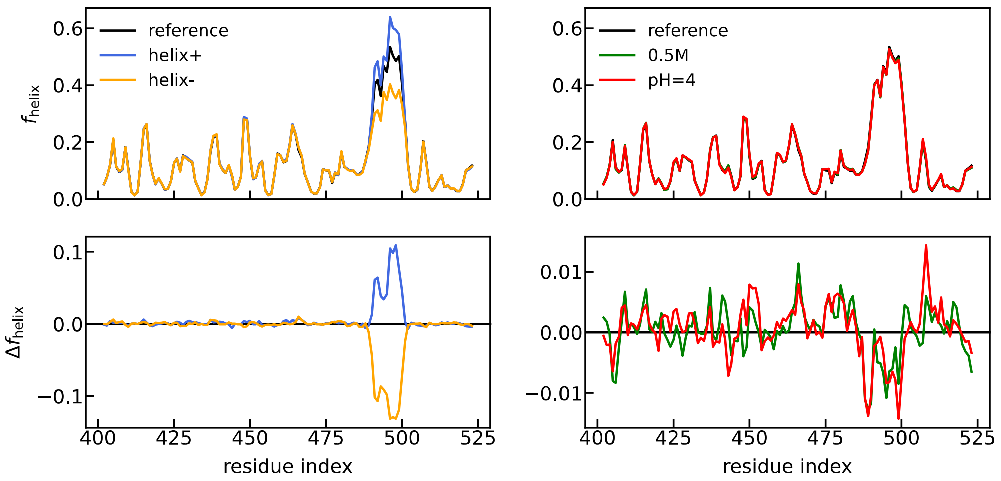

**Fig. S20.** Top left: Helical propensities of the CG model when varying a free parameter $\epsilon_{dih}$ controlling the stability of the helical minimum so as to introduce more helix (helix+ in blue with $\epsilon_{dih} = 0.4k\ cal/mol$) or less helix (helix- in orange with $\epsilon_{dih} = 0.1\ kcal/mol$ ). Top right: Helical propensities of the CG model at 0.5M salt (green) and pH=4 (red). The reference condition at 0.15 M salt, neutral pH and $\epsilon_{dih} = 0.2\ kcal/mol$ is shown in black. The corresponding difference of helical propensities to the reference condition is shown in the bottom.

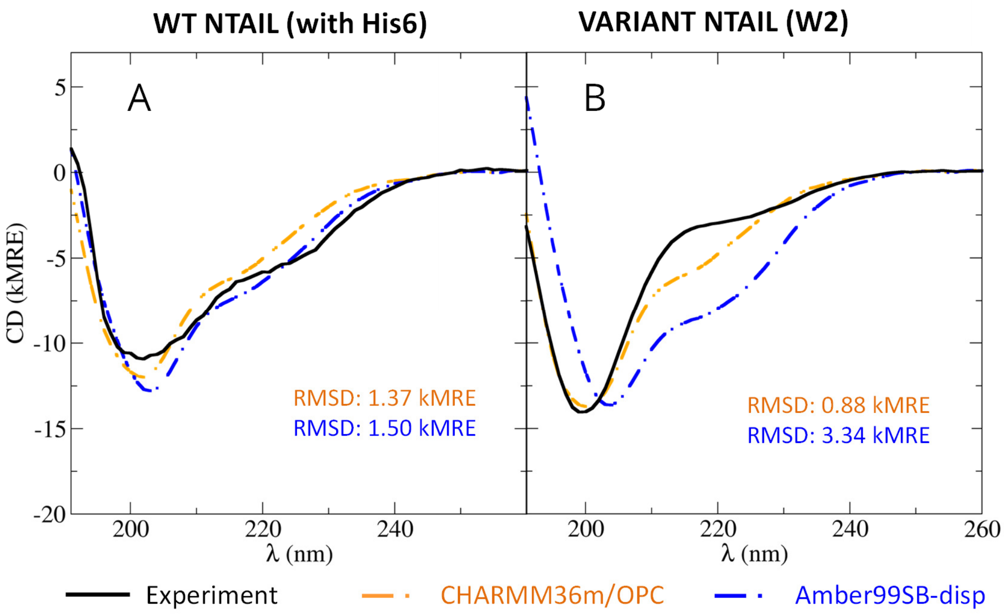

**Fig. S21** Predicted CD spectra calculated from the wild type (A) and variant (B) MD simulations. CD spectra predicted from CHARMM36m-OPC simulations are shown as orange dashed lines, whereas CD predictions from AMBER99SB-disp simulations blue dashed lines. The root-mean-squared deviation (RMSD) from the measured CD spectra (black solid lines) are shown in a corresponding color.

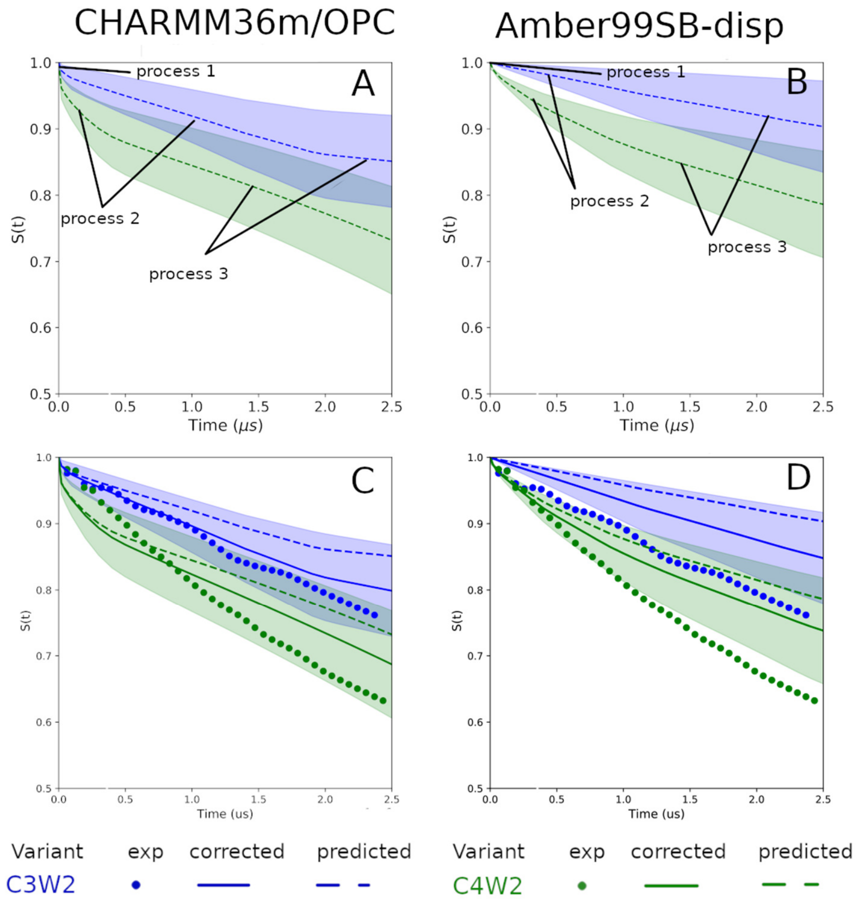

**Fig. S22.** PET experiment curves calculated from the CHARMM36m-OPC (left) and AMBER99SB-disp (right) force fields are shown as dashed lines, with their computational uncertainty depicted as shaded areas. Estimated PET data of C3W2 (S482C-Y518W) and C4W2 (S488C-Y518W) labelling positions are shown in blue and green, respectively. The uncorrected PET experiment curves in (A) and (B) indicate several dynamic processes present in all MD simulations with different rate-limiting steps that change the decay rate of the survival probabilities (*S(t)*). In panels (C) and (D) the computed decays are corrected for the estimated unquenched decay (solid lines), and compared to the measured decay curves (dots). The unquenched decay of 39.4 μs was estimated from relaxation decay of Y518W single variant as shown in Supporting Table S2.

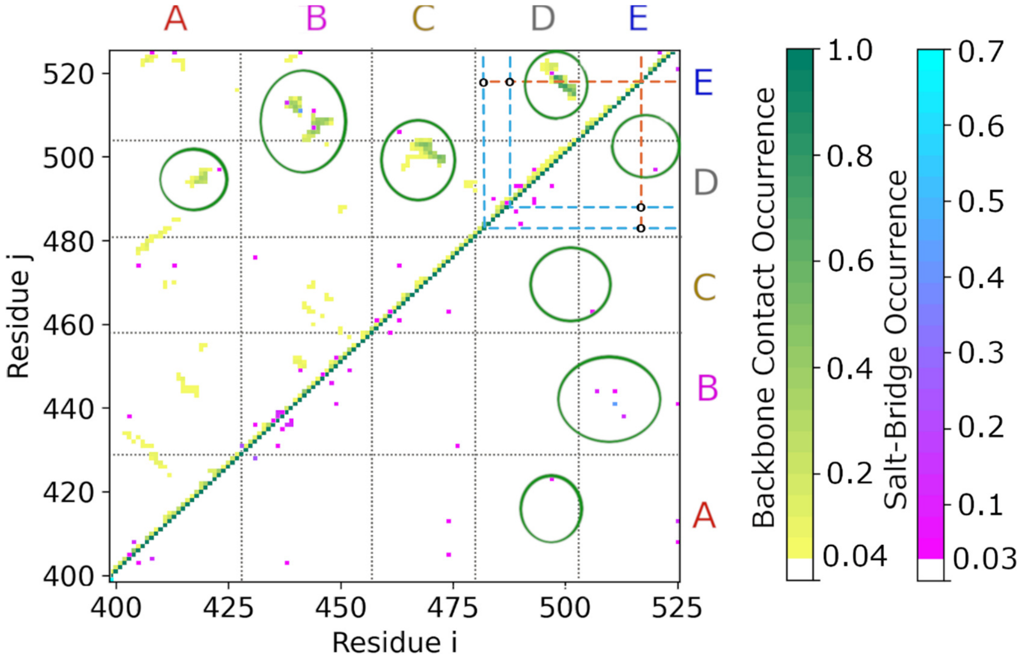

**Fig. S23.** Contact map for the S1 state shows Cα contact probabilities (above the diagonal, yellow to green) and salt bridge probabilities (above and below the diagonal, purple to blue). Cysteine acceptor positions C3 and C4 (residues 482 and 488) as well as tryptophan donor position W2 (residue 518) are indicated by cyan and orange dashed lines, respectively, and hypothetical C-W contacts are indicated by black circles. The boundaries of identified interaction regions A-E and are separated by black dotted lines on the contacts maps. The most prominent interactions in each state are encircled.

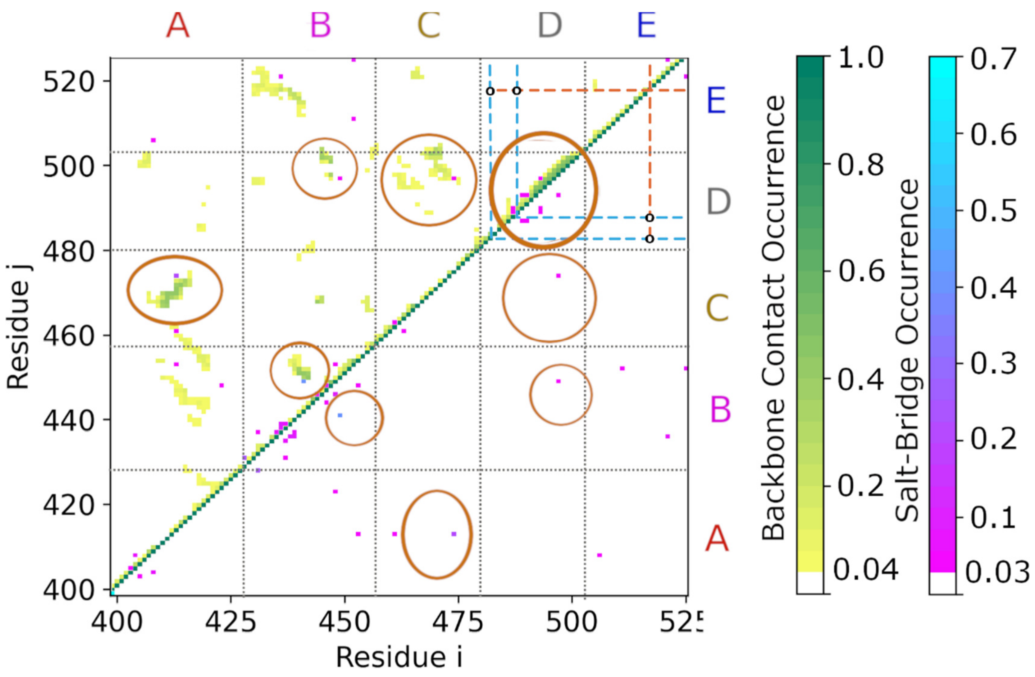

**Fig. S24.** Contact map for the S2 state shows Cα contact probabilities (above the diagonal, yellow to green) and salt bridge probabilities (above and below the diagonal, purple to blue). Cysteine acceptor positions C3 and C4 (residues 482 and 488) as well as tryptophan donor position W2 (residue 518) are indicated by cyan and orange dashed lines, respectively, and hypothetical C-W contacts are indicated by black circles. The boundaries of identified interaction regions A-E and are separated by black dotted lines on the contacts maps. The most prominent interactions in each state are encircled.

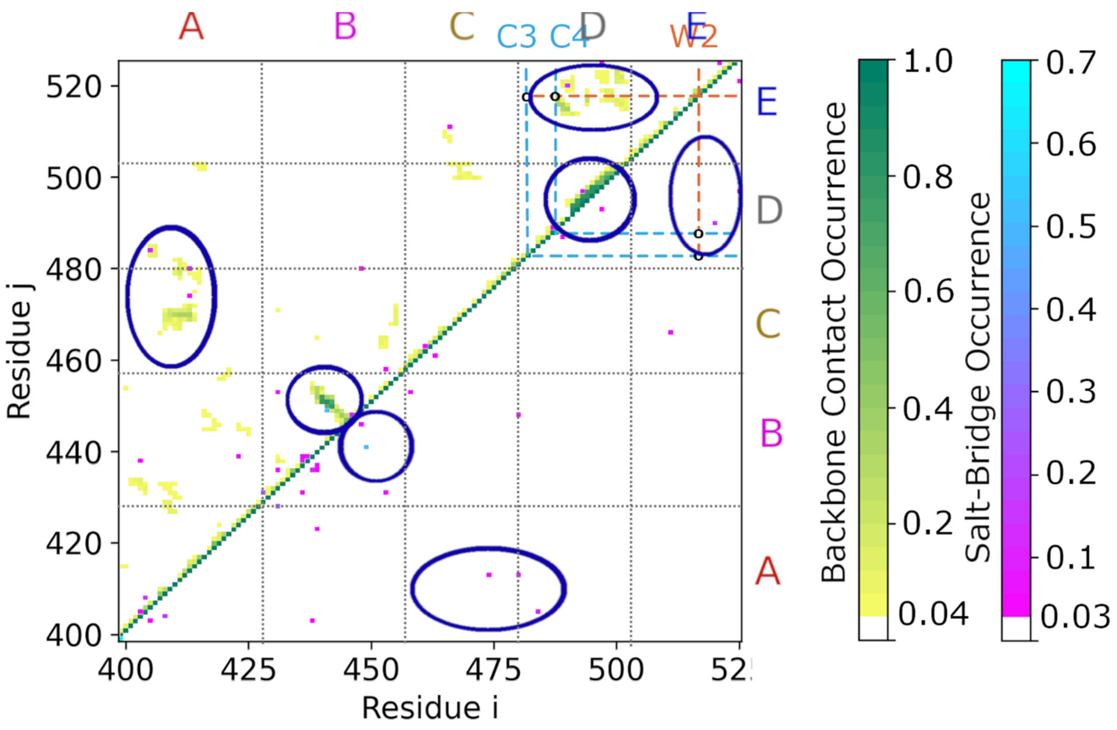

**Fig. S25.** Contact map for the S3 state shows $C_\alpha$ contact probabilities (above the diagonal, yellow to green) and salt bridge probabilities (above and below the diagonal, purple to blue). Cysteine acceptor positions C3 and C4 (residues 482 and 488) as well as tryptophan donor position W2 (residue 518) are indicated by cyan and orange dashed lines, respectively, and hypothetical C-W contacts are indicated by black circles. The boundaries of identified interaction regions A-E and are separated by black dotted lines on the contacts maps. The most prominent interactions in each state are encircled.

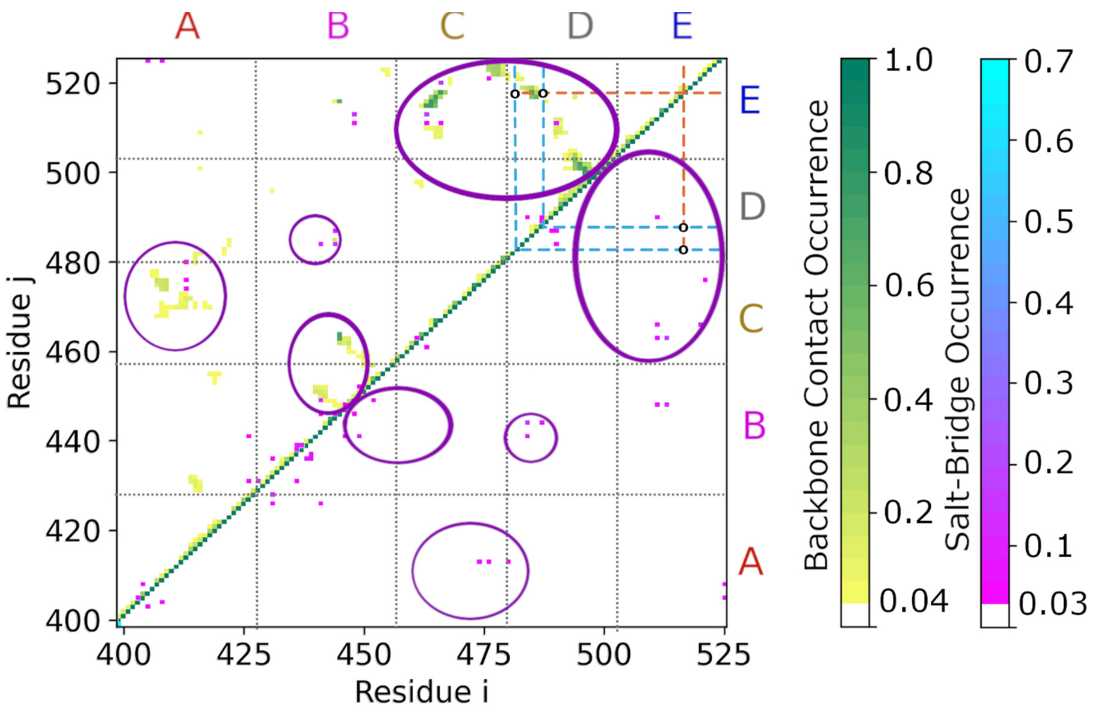

**Fig. S26.** Contact map for the S4 state shows Cα contact probabilities (above the diagonal, yellow to green) and salt bridge probabilities (above and below the diagonal, purple to blue). Cysteine acceptor positions C3 and C4 (residues 482 and 488) as well as tryptophan donor position W2 (residue 518) are indicated by cyan and orange dashed lines, respectively, and hypothetical C-W contacts are indicated by black circles. The boundaries of identified interaction regions A-E and are separated by black dotted lines on the contacts maps. The most prominent interactions in each state are encircled.

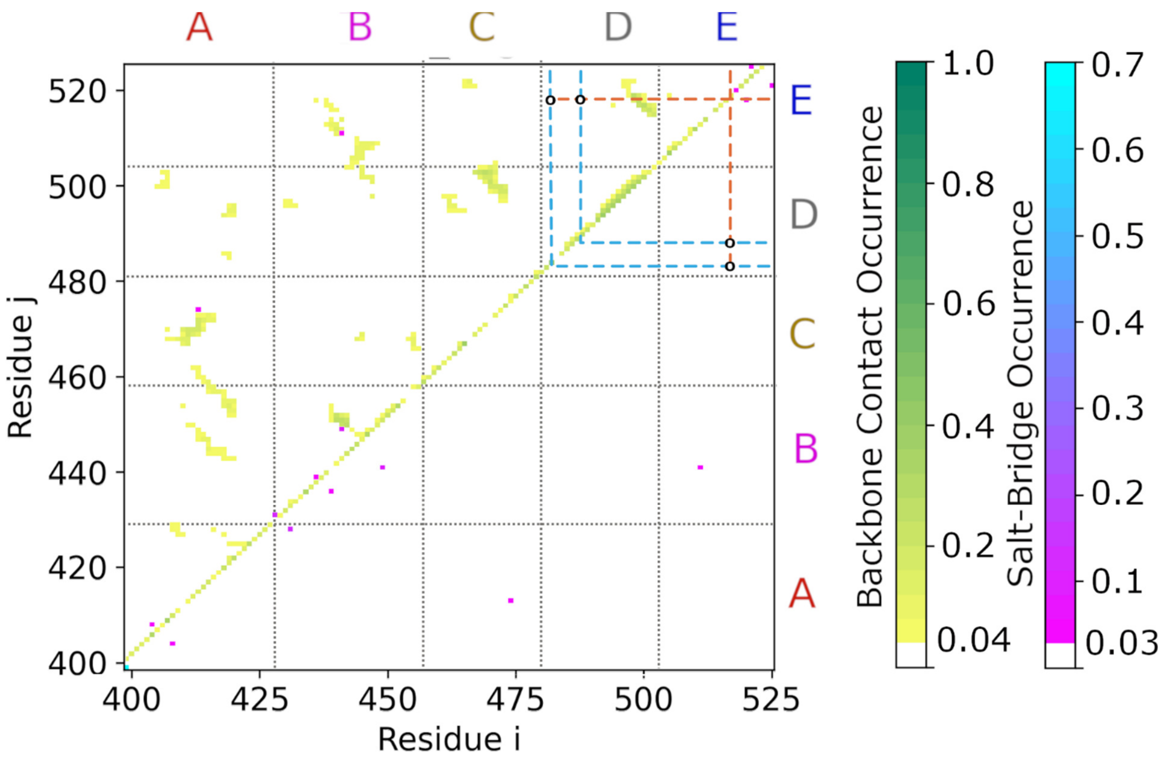

**Fig. S27.** Averaged contact map for all Charmm36m-OPC simulations. Contact map shows Cα contact probabilities (above the diagonal, yellow to green) and salt bridge probabilities (above and below the diagonal, purple to blue). Cysteine acceptor positions C3 and C4 (residues 482 and 488) as well as tryptophan donor position W2 (residue 518) are indicated by cyan and orange dashed lines, respectively, and hypothetical C-W contacts are indicated by black circles. The boundaries of identified interaction regions A-E and are separated by black dotted lines on the contacts maps.

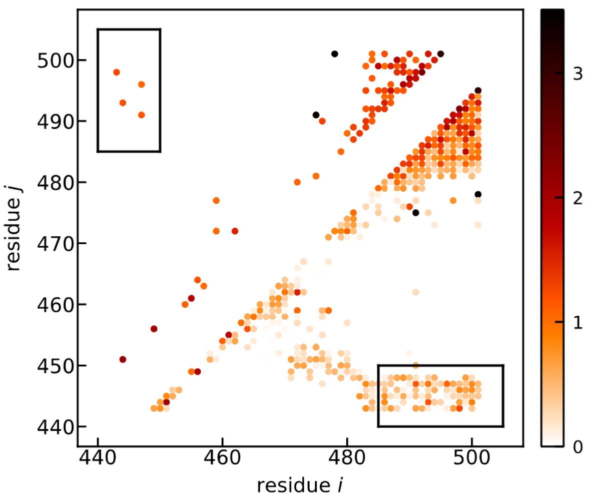

**Fig. S28.** Coupling score from using EVcouplings analysis. The region of the plot above the diagonal shows only the amino acid pairs with scores larger than 1. The region below the diagonal shows all pairs with scores larger than 0. The black box highlights the amino acid pairs between region B (residues 440-450) and region D (485-505)

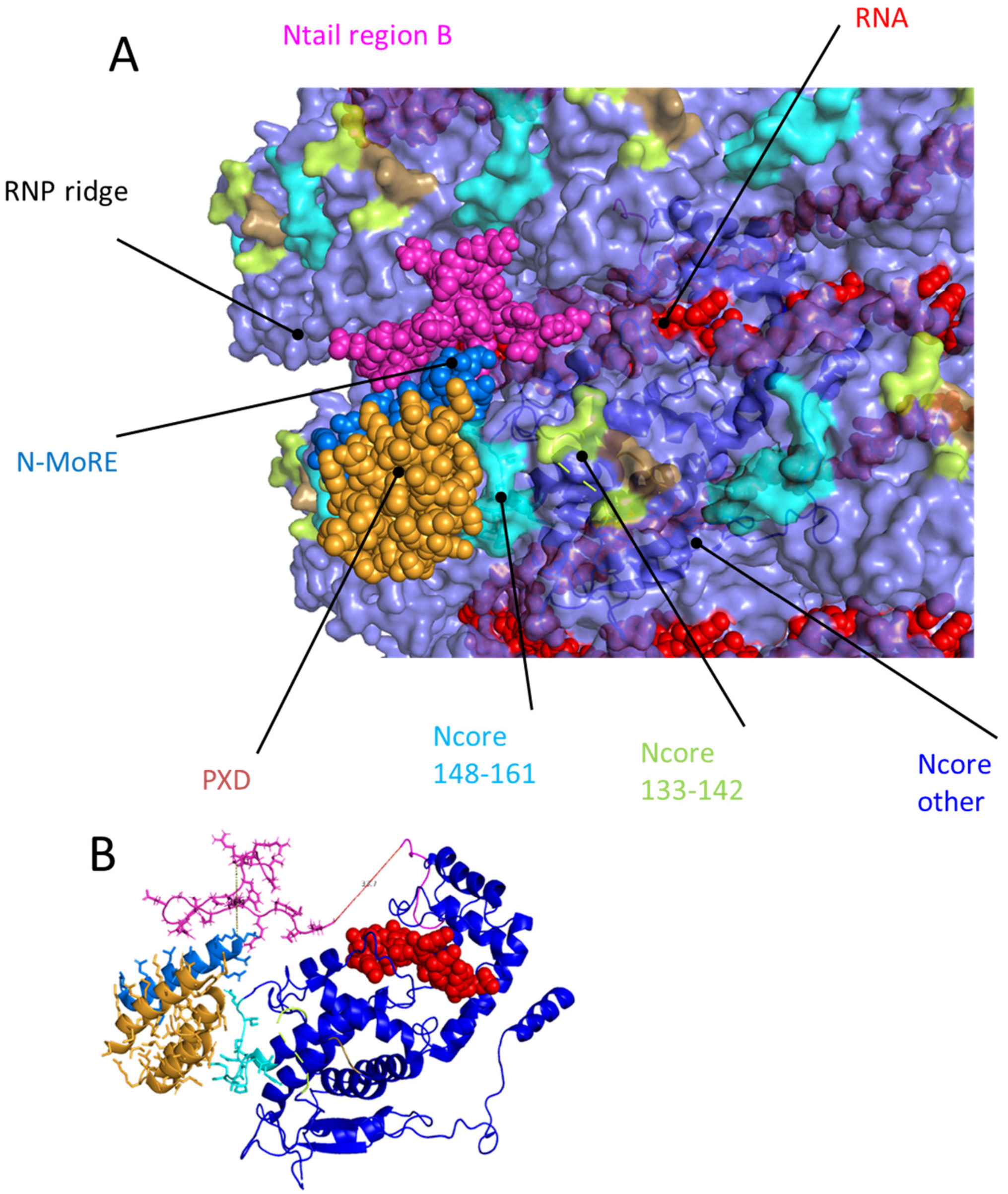

**Fig. S29**. Illustrative model of $N_{TAIL}$ in the nucleocapsid RNA-binding ridge. A) surface and sphere representation of the RNA binding ridge. B) Cartoon representation of the RNA binding ridge with only one N protomer displayed. The model shows how the $N_{TAIL}$ MoRE-PXD complex can bind $N_{CORE}$ at the RNA binding ridge, with sufficient space in the ridge to facilitate $N_{TAIL}$ intramolecular interactions between MoRE and region B. Note that Region B in this model is 3.5 nm away from the $C_{ARM}$ region of $N_{CORE}$, which may be bridged by the 28 amino acids of region A. This model was generated manually, based on the Cryo-EM structure of the nucleocapsid (pdb code: 4uft). The available structure of the $P_{XD}$-MoRE complex (pdb code: 1to6) was docked to the $P_{XD}$-$N_{CORE}$ binding site. Finally, a conformation of Ntail region B which contacted MoRE in the MD simulations was added into the nucleocapsid ridge.

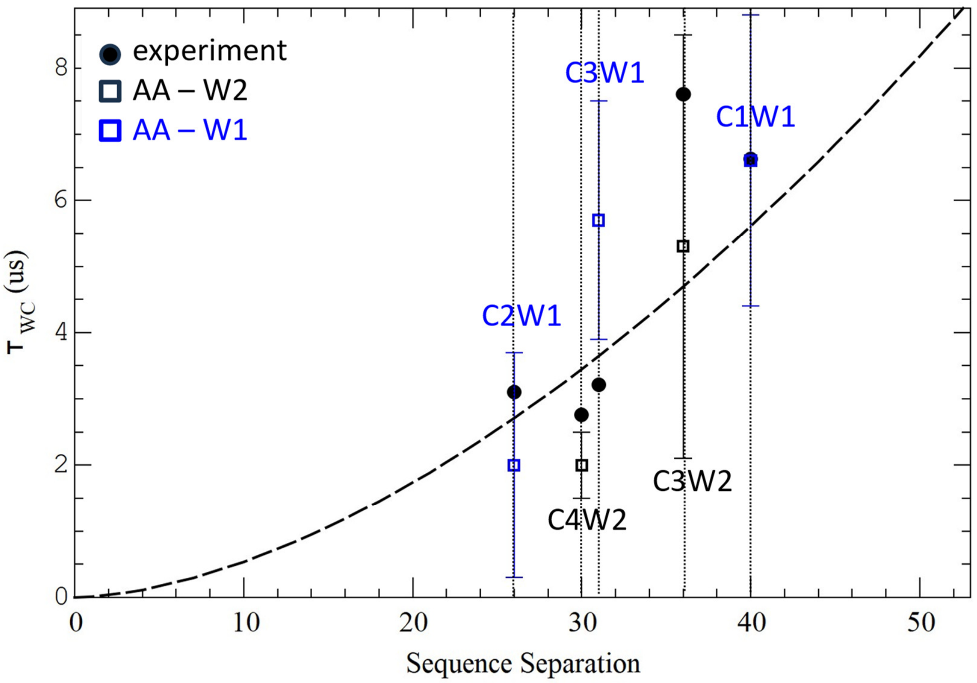

Fig. S30. Comparison between estimated C-W relaxation times for $N_{TAIL}$ PET variants obtained from W triplet-state relaxation experiments at the reference condition (circles, pH 7.6 and 150 mM NaCl concentration), and all-atom molecular dynamics (AA, squares) as function of donor-acceptor sequence separation. The estimated error of the estimated relaxation times is indicated by bars. The homopolymer model fitted to the measured relaxation times (Fig. 1D) is shown as a black dashed line. The AA relaxation times for the C1W1, C2W1, and C3W1 variants (blue squares) were estimated based S-Y contact distances with a cutoff distance 0.6 nm as described in Supplementary Methods S17.

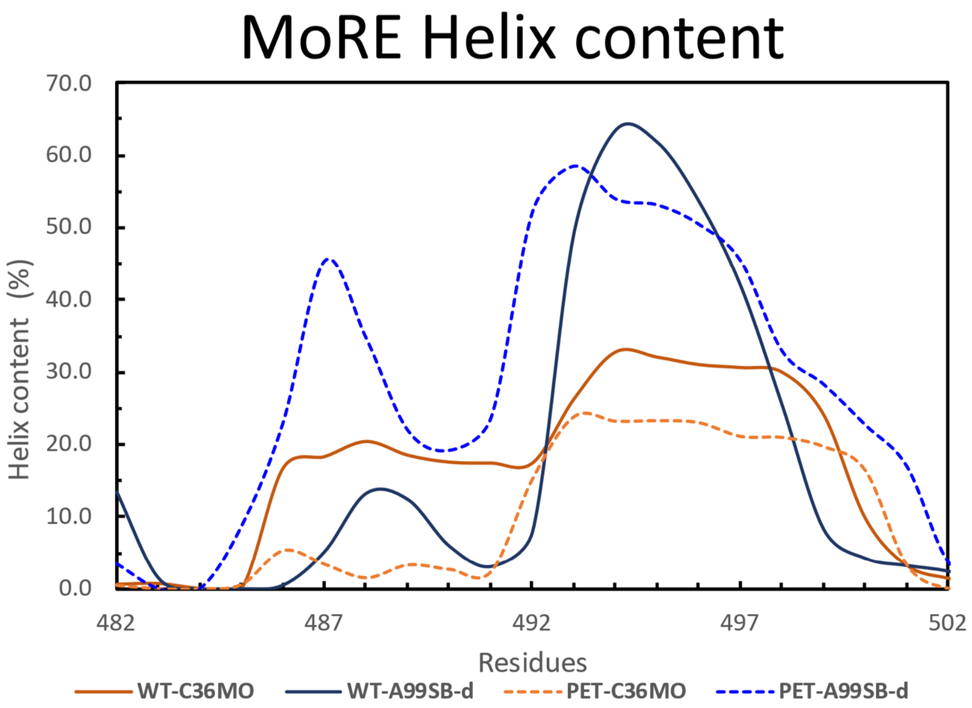

Forcefield WT-MoRE WT -total PET -MoRE PET -total
C36M-OPC 17% 3% 10% 2%
A99SB-disp 18% 8% 29% 12%

Fig S31: More Helix content in AA simulations. Top: The average helix content for residues in the MoRE region for WT (solid lines) and PET variant (dashed lines) simulations using the CHARMM36m-OPC (C36MO, orange) and Amber99SB-disp (A99SB-d, blue) force fields are shown. Bottom: the average helix content for MoRE region as a whole, and the full $N_{TAIL}$ (total) are shown.

#### C. Supporting Tables

**Table S1**. $N_{TAIL}$ variants and sequences used in the PET experiments. Residues in magenta denote acidic residues, blue letters indicate basic residues, cysteines are shown in cyan and tryptohans are highlighted in orange.

| Variant | Sequence |
|---|---|
| Wild type | GS 401TTEDKISRAV GPRQAQVSFL HGDQSENELP RLGGKEDRRV<br>441KQSRGEARES YRETGPSRAS DARAAHLPTG TPLDIDTASE<br>481SSQDPQDSRR SADALLRLQA MAGISEEQGS DTDTPIVYND<br>521RNLLD |
| Y518W (W2) | GS 401TTEDKISRAV GPRQAQVSFL HGDQSENELP RLGGKEDRRV<br>441KQSRGEARES YRETGPSRAS DARAAHLPTG TPLDIDTASE<br>481SSQDPQDSRR SADALLRLQA MAGISEEQGS DTDTPIVWND<br>521RNLLD |
| Y451W (W1) | GS 401TTEDKISRAV GPRQAQVSFL HGDQSENELP RLGGKEDRRV<br>441KQSRGEARES WRETGPSRAS DARAAHLPTG TPLDIDTASE<br>481SSQDPQDSRR SADALLRLQA MAGISEEQGS DTDTPIVYND<br>521RNLLD |
| S425C-Y451W (C2W1) | GS 401TTEDKISRAV GPRQAQVSFL HGDQCENELP RLGGKEDRRV<br>441KQSRGEARES WRETGPSRAS DARAAHLPTG TPLDIDTASE<br>481SSQDPQDSRR SADALLRLQA MAGISEEQGS DTDTPIVYND<br>521RNLLD |
| S488C-Y518W (C4W2) | GS 401TTEDKISRAV GPRQAQVSFL HGDQSENELP RLGGKEDRRV<br>441KQSRGEARES YRETGPSRAS DARAAHLPTG TPLDIDTASE<br>481SSQDPQDCRR SADALLRLQA MAGISEEQGS DTDTPIVWND<br>521RNLLD |
| Y451W-S482C (C3W1) | GS 401TTEDKISRAV GPRQAQVSFL HGDQSENELP RLGGKEDRRV<br>441KQSRGEARES WRETGPSRAS DARAAHLPTG TPLDIDTASE<br>481SCQDPQDSRR SADALLRLQA MAGISEEQGS DTDTPIVYND<br>521RNLLD |
| S482C-Y518W (C3W2) | GS 401TTEDKISRAV GPRQAQVSFL HGDQSENELP RLGGKEDRRV<br>441KQSRGEARES YRETGPSRAS DARAAHLPTG TPLDIDTASE<br>481SCQDPQDSRR SADALLRLQA MAGISEEQGS DTDTPIVWND<br>521RNLLD |
| S407C-Y451W (C1W1) | GS 401TTEDKICRAV GPRQAQVSFL HGDQSENELP RLGGKEDRRV<br>441KQSRGEARES WRETGPSRAS DARAAHLPTG TPLDIDTASE<br>481SSQDPQDSRR SADALLRLQA MAGISEEQGS DTDTPIVYND<br>521RNLLD |

**Table S2.** Measurements, conditions, and fitting parameters obtained after global analysis of the transient absorption data for each $N_{TAIL}$ variant studied. Only measurements for which the amplitude $A_1$ of the main decay ($\tau_1$) contributed 35% or more to the total signal amplitude were included in the global fit (see Supporting Methods S2 for more details). $|i\text{-}j|$ refers to the sequence separation between the tryptophan and cysteine for each variant. Solvent conditions only include the variable conditions (see section Supporting Methods S2 for full solvent details). The blue highlighted columns denote the main decay $\tau_1$, used to obtain $\tau_{CW}$ values reported in main text, and shown in Figure 1D (from $\tau_{CW}^{-1} = \tau_1^{-1} - \tau_o^{-1}$, where $\tau_o$ is the main decay measured on variants containing only W, highlighted in blue). There is a ±5% error on the $\tau_{CW}$ values. The amplitudes $(A_1, A_2, \dots A_5)$ are the averages of the percentage contribution of the first measurement of each sample for each variant, before significant photodamage began. The $\tau_5$ component with its negative percentage amplitude $A_5$ is the rise time.

| Variant | \|i-j\| | Solvent Conditions | Number of Measurements | $\tau_1$ (µs) | $A_1$ (%) | $\tau_2$ (µs) | $A_2$ (%) | $\tau_3$ (µs) | $A_3$ (%) | $\tau_4$ (s) | $A_4$ (%) | $\tau_5$ (ns) | $A_5$ (%) | $\tau_{CW}$ (µs) |
|---|---|---|---|---|---|---|---|---|---|---|---|---|---|---|
| Y451W (W1) | -- | pH 7.6 150mM NaCl | 11 | 23.0 | 81 | -- | -- | 554 | 15 | 1 | 4 | 34 | -22 | -- |
| Y518W (W2) | -- | pH 7.6 150mM NaCl | 14 | 39.4 | 78 | 2.43 | 6 | 605 | 13 | 1 | 3 | 30 | -26 | -- |
| | | pH 4.0 150mM NaCl | 4 | 4.06 | 61 | 31.6 | 32 | 554 | 6 | 1 | 1 | 28 | -30 | -- |
| S425C-Y451W (C2W1) | 26 | pH 7.6 150mM NaCl | 10 | 2.73 | 57 | 27.7 | 29 | 710 | 7 | 1 | 7 | 31 | -25 | 3.10 |
| S488C-Y518W (C4W2) | 30 | pH 7.6 150mM NaCl | 9 | 2.58 | 62 | 29.5 | 23 | 694 | 7 | 1 | 8 | 35 | -24 | 2.76 |
| | | pH 4.0 150mM NaCl | 8 | 1.84 | 68 | 28.8 | 21 | 849 | 6 | 1 | 6 | 29 | -16 | 3.38 |
| Y451W-S482C (C3W1) | 31 | pH 7.6 150mM NaCl | 12 | 2.82 | 43 | 21.8 | 40 | 842 | 13 | 1 | 4 | 42 | -24 | 3.21 |
| | | pH 4.0 150mM NaCl | 7 | 1.88 | 71 | 31.9 | 18 | 812 | 4 | 1 | 7 | 19 | -27 | 3.52 |
| | | pH 7.6 500mM NaCl | 6 | 3.33 | 57 | 19.0 | 31 | 1120 | 9 | 1 | 4 | 28 | -30 | 3.89 |
| S482C-Y518W (C3W2) | 36 | pH 7.6 150mM NaCl | 16 | 6.37 | 56 | 48.1 | 30 | 1020 | 9 | 1 | 5 | 44 | -25 | 7.60 |
| | | pH 4.0 150mM NaCl | 8 | 2.26 | 49 | 23.6 | 36 | 233 | 10 | 1 | 5 | 25 | -26 | 5.11 |
| | | pH 7.6 500mM NaCl | 3 | 5.08 | 52 | 43.0 | 33 | 1060 | 12 | 1 | 4 | 47 | -26 | 5.83 |
| S407C-Y451W (C1W1) | 40 | pH 7.6 150mM NaCl | 14 | 5.15 | 45 | 33.0 | 39 | 930 | 10 | 1 | 6 | 44 | -26 | 6.63 |
| | | pH 4.0 150mM NaCl | 5 | 2.71 | 81 | 15.7 | 9 | 481 | 3 | 1 | 7 | 37 | -21 | 8.19 |

**Table S3**. Force field comparison of wild type $N_{TAIL}$ to experimental data. The table shows the deviation between measured observables (SAXS intensities, CD spectra, and NMR chemical shifts) and the same observables predicted from simulation ensembles of particular forcefileds (FF, see Supporting Methods S8). The experimental radius of gyration was determined based on the measured SAXS curve using CRYSOL[48]. Experimental SS composition was estimated from the CD spectrum using SESCA bayes, and the experimental CD deviation is calculated by comparing the predicted spectrum of the estimated SS composition to the measured CD spectrum. A structural ensemble of MEV-$N_{TAIL}$ from the protein ensemble databank (PED) fitted to match measured chemical shifts is also shown for comparison. The exceptionally good (SD-HQ) and maximum acceptable (SD-max) deviations for the considered observables for a high-quality ensemble are shown on the bottom.

| **Summary** | **SAXS data** | | **CD spectra** | **Cα shifts** | **Cβ shifts** | **SS content (%)** | | |
|---|---|---|---|---|---|---|---|---|
| **System/ FF** | **χ (A.U.)** | **Rg (nm)** | **RMSD (kMRE)** | **RMSD (ppm)** | **RMSD (ppm)** | **Alpha** | **Beta** | **Other** |
| **Exp.** | 0.00 | 2.75 | 0.87 | 0.00 | 0.00 | 3.3 | 13.6 | 83.1 |
| **PED*** | 0.61 | 3.28 | 1.46 | 0.29 | 0.40 | 7.3 | 2.5 | 90.4 |
| **C36M-OPC** | 0.63 | 2.99 | 1.37 | 0.54 | 0.36 | 3.3 | 6.5 | 90.2 |
| **A99B-disp** | 0.79 | 2.73 | 1.50 | 0.44 | 0.44 | 7.9 | 2.9 | 89.2 |
| **A03-ws** | 0.49 | 2.91 | 2.88 | 0.66 | 0.49 | 16.0 | 6.2 | 77.8 |
| **A99sb-ws** | 1.67 | 2.63 | 2.98 | 0.43 | 0.35 | 16.7 | 4.9 | 78.4 |
| **C36M** | 2.29 | 2.07 | 0.95 | 0.56 | 0.49 | 4.6 | 7.0 | 88.4 |
| **C36M-Tip4** | 2.64 | 1.92 | 1.08 | 0.61 | 0.55 | 3.9 | 7.8 | 88.3 |
| **C22*-Tip3S** | 2.57 | 1.95 | 0.65 | 0.51 | 0.46 | 6.1 | 6.6 | 87.2 |
| **C22*-OPC** | 1.93 | 2.19 | 0.75 | 0.58 | 0.51 | 5.4 | 7.1 | 87.5 |
| **SD-HQ** | 1.0 | 0.3 | 1.1 | 0.21 | 0.42 | 4.0 | 6.0 | 9.0 |
| **SD-max** | 2.0 | 0.6 | 3.1 | 0.75 | 0.62 | 8.0 | 12.0 | 18.0 |

**Table S4**. Force field comparison of NTAIL variants to experimental data. The table shows the deviation between measured observables (C-W quenching times due to PET relaxation and CD spectra) and the same observables predicted from simulation ensembles of particular force fields (FF, see Supporting Methods S8). The experimental C-W quenching times $\tau_{CW}$ are calculated from direct fits to the measured decay curves for each variant, and corrected for the natural decay $\tau_0$ without the C quencher (as in Supporting Table S2). Experimental SS composition was estimated from the CD spectrum using SESCA bayes, and the experimental CD deviation is calculated by comparing the predicted spectrum of the estimated SS composition to the measured CD spectrum. The uncertainty of relaxation times shows the standard deviation of the posterior relaxation time distribution calculated from a Bayesian fit to the bootstrapped relaxation curves of all-atom MD simulation.

| Summary | PET predictions | | CD spectra | Compactness | SS content (%) | | | Ensemble |
|---|---|---|---|---|---|---|---|---|
| System/FF | C3W2 (μs) | C4W2 (μs) | RMSD (kMRE) | Rg (nm) | Alpha | Beta | Other | size |
| **experiment** | 7.6 | 2.8 | 0.708 | 2.72 | 0.9 | 1.6 | 97.5 | - |
| **C36M-OPC** | 5.3 (3.3) | 2.0 (0.5) | 0.884 | 2.49 | 1.9 | 7.1 | 90.9 | 87785 |
| **A99SB-disp** | 13.3 (3.2) | 4.7 (1.0) | 3.431 | 2.26 | 11.7 | 3.2 | 85.2 | 103945 |

**Table S5.** Normalized pointwise mutual Information analysis of $N_{TAIL}$ variant simulation trajectories from the CHARMM36m/OPC simulations. Prominent non-local interactions of Fig. 4 of main manuscript were analyzed to determine their correlation with PET donor-acceptor contact formation (both with C3W2 or C4W2 contacts). The residue numbers of the contact pair (residue 1 and residue 2), residue and region codes are also indicated in the table. Finally, the NPMI scores for conserved contacts found during our Coevolution analysis are also shown on the bottom.

| **S1 interactions** | | | | **NPMI** | |
|---|---|---|---|---|---|
| **Residue 1** | **Residue 2** | **residue** | **Regions** | **C3W2** | **C4W2** |
| 413 | 525 | R-D | A/E | -0.31 | -0.44 |
| 441 | 511 | K-E | B/E | -0.44 | -0.25 |
| 444 | 507 | R-E | B/E | -0.42 | -0.24 |
| 497 | 520 | R-D | D/E | -0.38 | -0.19 |
| 420 | 495 | L-L | A/D | -0.42 | -0.56 |
| 472 | 501 | M-L | C/D | -0.29 | -0.31 |
| 500 | 518 | A-W | D/E | -0.39 | -0.30 |
| **S2 interactions:** | | | | | |
| 409 | 469 | A-T | A/C | -0.02 | 0.16 |
| 413 | 476 | R-D | A/C | 0.30 | 0.03 |
| 441 | 449 | K-E | B/B | 0.03 | 0.28 |
| 447 | 501 | A-M | B/D | -0.36 | -0.42 |
| 497 | 453 | R-E | B/D | -0.14 | -0.01 |
| **S3 interactions:** | | | | | |
| 405 | 484 | R-D | A/D | 0.08 | 0.49 |
| 413 | 480 | R-E | A/D | 0.27 | 0.40 |
| 441 | 449 | K-E | B/B | 0.03 | 0.28 |
| 490 | 520 | R-D | D/E | -0.04 | 0.70 |
| **S4 interactions:** | | | | | |
| 413 | 476 | R-D | A/C | 0.30 | 0.03 |
| 444 | 487 | R-D | B/D | 0.29 | -0.37 |
| 463 | 513 | R-E | C/E | 0.42 | -0.42 |
| 490 | 511 | R-E | D/E | 0.31 | -0.21 |
| 465 | 516 | A-I | C/E | 0.45 | -0.47 |
| **Coevolution:** | | | | | |
| 444 | 493 | R-D | B/D | -0.06 | -0.20 |
| 443 | 498 | S-L | B/D | -0.04 | -0.10 |
| 447 | 491 | A-L | B/D | -0.03 | -0.17 |
| 447 | 496 | A-L | B/D | 0.00 | -0.14 |

**Table S6.** Normalized pointwise mutual Information analysis of $N_{TAIL}$ variant simulation trajectories from the CHARMM36m/OPC simulations. Prominent non-local interactions of Fig. 4 of main manuscript were analyzed to determine their correlation with the helical conformations of $N_{TAIL}$ MoRE. The helical conformations are defined by the fraction of MoRE residues in helical conformations; low (H-low) below 0.2, medium (H-med) between 0.2-0.5, and high (H-high) above 0.5. The residue numbers of the contact pair (residue 1 and residue 2), residue and region codes are also indicated in the table. Finally, the NPMI scores for prominent contacts involving the phosphoprotein XD binding site are also shown on the bottom. Contacts that have strong negative correlations with helix content in the MoRE are shown in red, ones that are positively correlated with medium helix content only are shown in blue, ones that are positively correlated with helix content are shown in green.

| **Residue 1** | **Residue 2** | **Res. type** | **Regions** | | **NPMI** | |
|---|---|---|---|---|---|---|
| **#S1 specific:** | | | | **H-low** | **H-med** | **H-high** |
| 413 | 525 | R-D | A/E | 0.07 | -0.61 | -0.42 |
| 441 | 511 | K-E | B/E | 0.10 | -0.34 | -0.05 |
| 444 | 507 | R-E | B/E | 0.11 | -0.50 | -0.29 |
| 497 | 520 | R-D | D/E | 0.04 | -0.08 | -0.50 |
| 420 | 495 | L-L | A/D | -0.01 | 0.04 | -0.12 |
| 472 | 501 | M-L | C/D | 0.06 | -0.27 | -0.41 |
| 500 | 518 | A-W | D/E | 0.07 | -0.19 | -0.51 |
| #S2 specific: | | | | | | |
| 409 | 469 | A-T | A/C | -0.08 | 0.17 | -0.41 |
| 413 | 476 | R-D | A/C | 0.01 | -0.50 | -0.08 |
| 441 | 449 | K-E | B/B | -0.01 | 0.77 | 0.16 |
| 447 | 501 | A-M | B/D | -0.70 | 0.56 | 0.44 |
| 497 | 453 | R-E | B/D | 0.03 | -0.30 | -0.12 |
| #S3 specific: | | | | | | |
| 405 | 484 | R-D | A/D | -0.32 | 0.33 | -0.44 |
| 413 | 480 | R-E | A/D | 0.30 | 0.03 | 0.08 |
| 441 | 449 | K-E | B/B | -0.01 | 0.77 | 0.16 |
| 490 | 520 | R-D | D/E | 0.03 | 0.28 | 0.49 |
| #S4 specific: | | | | | | |
| 413 | 476 | R-D | A/C | 0.06 | -0.41 | -0.36 |
| 444 | 487 | R-D | B/D | 0.06 | -0.53 | -0.35 |
| 463 | 513 | R-E | C/E | -0.02 | 0.05 | -0.40 |
| 490 | 511 | R-E | D/E | 0.04 | -0.16 | -0.32 |
| 465 | 516 | A-I | C/E | 0.08 | -0.48 | -0.45 |
| #XD-bind: | | | | | | |
| 447 | 498 | A-L | B/D | -0.62 | 0.57 | 0.47 |
| 474 | 497 | D-R | B/D | 0.06 | -0.18 | -0.56 |
| 491 | 514 | S-T | D/E | -0.21 | 0.34 | -0.49 |
| 498 | 523 | L-L | D/E | 0.03 | -0.01 | -0.56 |
| 501 | 516 | M-I | D/E | 0.13 | -0.25 | -0.48 |
| 501 | 522 | M-N | D/E | -0.21 | 0.35 | -0.52 |

**Table S7**. MoRE binding residue (on the left) interactions in the conformational states (S1-S4) identified in Fig. 3 of the main manuscript. The residue numbers of significant interaction partners, the highest contact formation probability of those partners, and region code for those residues are indicated in each state. We note that MoRE residues are located in region D of $N_{TAIL}$.

| MoRE Residue | S1 partners<br>Resid, Pmax, region | S2 partners<br>Resid, Pmax, region | S3 partners<br>Resid, Pmax, region | S4 partners<br>Resid, Pmax, region |
|---|---|---|---|---|
| **S491** | ---- | ---- | 514, 0.32, E<br>519-522, 0.18, E | 509-512, 0.15, E |
| **L495** | 417-420, 0.34, A | 463-465, 0.08, C | ----- | 501-503, 0.71, D |
| **L498** | 518-519, 0.73, E | 447, 0.35, B<br>472-473, 0.14, C<br>461-463, 0.11, C | 517-519, 0.17, E | ----- |
| **M501** | 515-517, 0.62, E<br>469-471, 0.41, C | 445-447, 0.37, B<br>406-407, 0.25, A<br>470, 0.14, C | 521-524, 0.34, E<br>515-517, 0.17, E | 495-497, 0.74, D |
| **W518** | 498-500, 0.75, D | 433-438, 0.25, B | 488, 0.58, D<br>498-499, 0.17, D<br>494, 0.25, D | 465-467, 0.74, C<br>484-486, 0.70, D |

**Table S8**. Further details on all-atom molecular dynamics (AA) simulations and force field comparison. For particular forcefileds (FF, see Supporting Methods S8), the number replicate trajectories (Ntraj) started from different initial conformations, the trajectory length range (length) in microsecond, and the number of frames used in the final ensemble analyzed.

| System/ FF | Ntraj | length (us) | No. of frames |
|---|---|---|---|
| $N_{TAIL}$ wild type (with His6 tag) | | | |
| C36M-OPC | 6 | 5-10 | 41230 |
| A99B-disp | 3 | 10 | 30000 |
| A03-ws | 3 | 2-3 | 7531 |
| A99sb-ws | 5 | 2-5 | 17460 |
| C36M | 5 | 1-3 | 10440 |
| C36M-Tip4 | 3 | 1-3 | 7524 |
| C22*-Tip3S | 3 | 4-7 | 17233 |
| C22*-OPC | 3 | 4-7 | 16607 |
| $N_{TAIL}$ Variants (C3W2, C4W2) | | | |
| C36M-OPC | 2X3 | 6-20 | 87790 |
| A99B-disp | 2X3 | 13-20 | 103961 |

**Table S9**. Coarse-grained (CG) molecular dynamics simulation details, including simulation conditions, simulation length, radius of gyration ($R_g$), and scaling exponent ($\nu$) computed from the trajectories.

| **Conditions** | **length (equilibration)** | $R_g$ **(nm)** | **scaling exponent (** $\nu$ **)** |
|---|---|---|---|
| 0.15M salt, pH=7.6, $k_h$=0.2 kcal/mol | 5μs (100ns) | 2.971 (3) | 0.5382 (2) |
| 0.5M salt | 5μs (100ns) | 3.035 (2) | 0.5436 (2) |
| pH=4 | 5μs (100ns) | 3.015 (4) | 0.5414 (3) |
| helix+, $k_h$=0.4 kcal/mol | 5μs (100ns) | 3.014 (1) | 0.5415 (3) |
| helix-, $k_h$=0.1 kcal/mol | 5μs (100ns) | 2.945 (3) | 0.5360 (2) |
